# Complex responses of soil prokaryotes, fungi and protists to prairie restoration on retired agricultural lands

**DOI:** 10.1101/2024.10.11.617895

**Authors:** Micaela Tosi, Kevin MacColl, Dasiel Obregón, Andrew S. MacDougall, Hafiz Maherali, Kari Dunfield

## Abstract

Restoring native ecosystems on marginal croplands has many benefits but the impacts on belowground biodiversity are less clear, in part because the limiting factors regulating soil biota are complex and poorly described. Here, we studied how grassland prairie restoration of marginal croplands affected the diversity and composition of soil microbiota on 5 conventional farms from Ontario, Canada. Soil samples (0-15 cm) were collected from annually cultivated fields and adjacent planted perennial grassland where cultivation and chemical inputs had ceased several years previously. Following DNA extraction, we estimated bacterial and fungal abundance using quantitative PCR, and microbial diversity of prokaryotes, fungi and protists using amplicon high-throughput sequencing. Under both land uses, prokaryotic communities were dominated by Proteobacteria, Actinobacteria and Acidobacteria, fungal communities by Ascomycota, and protist communities by Rhizaria (TSAR), Evosea (Amoebozoa) and Chlorophyta (Archaeplastida). Prairie restoration did not have a consistent effect on soil microbial abundance, richness or evenness, which responses varied across farms. Microbial genetic and taxonomic community composition (*i.e.*, sequence variant and genus level) were affected by land use, farm and the interaction between these two factors. Generally, prairie soils had higher relative abundance of Latescibacterota, Desulfobacterota, Acidobacteriota and Glomeromycota, and lower of Deinococcota, Chytridiomycota and Amoebozoa_X. In terms of differentially abundant fungal genera, prairies promoted more fungal plant symbionts, less saprotrophs and no plant pathogens. Interkingdom networks revealed changes in potential microbe-microbe associations with prairie restoration, with only 8 associations in common between land uses. The relationship between soil microbial diversity and physicochemical properties varied across microbial groups, diversity metrics and land uses. Our results evidence the complexity associated with restoring soils from agricultural land to natural ecosystems, with unspecified farm-specific factors (*e.g.*, soil type, prairie species, management history) strongly modulating the response of different microbial groups and variables.

## 1- Introduction

Driven by the need to support a growing population, the cultivation of the world’s arable lands may increasingly threaten ecosystem health and sustainability (Foley et al., 2005). Conventional agricultural practices are characterized by reduced plant diversity and plant cover, high physical disturbance from tillage and other equipment, and high external inputs of mineral fertilizers and pesticides. These practices alter the soil environment and its living communities, with consequences including organic carbon losses, nutrient imbalances, increased risk of erosion, increased greenhouse gas emissions and biodiversity loss (Trivedi et al., 2016; Dudley and Alexander, 2017; Yang et al., 2021). Environmental concerns become particularly pressing as the global footprint of annual cultivation widens. According to the Food and Agriculture Organization (FAO) of the United Nations (UN), land use destinated to crops has increased ∼15% between 1961 and 2019, now covering >30% of the earth’s terrestrial surface and resulting in land and soil degradation and water scarcity (FAO, 2022).

To some extent, the negative environmental consequences of agriculture can be alleviated by increasing ecological intensification via sustainable management practices, but financial and logistic factors can sometimes limit the application of such practices (Wezel et al., 2014; Kleijn et al., 2019). In the case of highly degraded or marginal farmlands, retirement and restoration of native ecosystems might be a more suitable way to recover the provision and spillover of ecosystem services while maintaining economically viable farms (Lamb et al., 2016; Yang et al., 2020; MacDougall et al., 2024). For example, in the study sites evaluated here, restoration of prairie grasslands increased the abundance and diversity of terrestrial arthropods, while also increasing the abundance of beneficial (*e.g.*, pollinators and pest predators) as opposed to herbivorous species (Dolezal et al., 2022). Restoration is also expected to benefit soil health as a consequence of reduced disturbance, changes in plant cover (coverage, diversity), and changes in the quantity and quality of plant inputs (Mariotte et al., 2018). Such changes can reduce erosion, improve soil structure (especially aggregation), increase soil organic carbon and reduce nutrient losses (Baer et al., 2002, 2015; Chandrasoma et al., 2016; Rosenzweig et al., 2016; De et al., 2020; Li et al., 2021). However, the response of soil properties and soil health could be slow and dependent on location-specific factors such as management history, soil type, topographic position and climate (Matamala et al., 2008; De et al., 2020; Mazzorato et al., 2022; Kimmell et al., 2023).

Soils host a vast biodiversity that has a crucial role in nutrient and carbon cycling, soil structure and plant health, and thus the provision of ecosystem services (Anthony et al., 2023). As such, soil organisms are likely to play a crucial role as facilitators of ecosystem restoration, for example by changing edaphic conditions and facilitating the establishment of the re-introduced vegetation (*e.g.*, via plant mutualistic fungi or bacteria) (Harris, 2009; Coban et al., 2022; Graham and Knelman, 2023). Despite this, and the increasing efforts towards ecosystem restoration (*e.g.*, UN Decade on Ecosystem Restoration, decadeonrestoration.org), soils and soil organisms have been largely neglected in ecosystem restoration studies (Callaham et al., 2008; Farrell et al., 2020). This is partly explained by methodological challenges to study soil organisms, as well as the difficulty to establish reference conditions or clear goals to monitor restoration success (Harris, 2009; Farrell et al., 2020). Recently, research on how soil biota responds to and contributes to restoration has intensified, with promising insights on microbial tools to aid restoration efforts (Robinson et al., 2023; Sáez-Sandino et al., 2023; Song, 2023).

Soil biota is expected to respond to restoration practices because they drastically modify the edaphic environment relative to agricultural fields (e.g., providing year-round plant cover, increasing root biomass, introducing more diverse rhizodeposits, and reducing physical disturbance). Based on these changes, prairie restoration could increase soil biodiversity, microbial and faunal biomass, functionality, and trophic complexity (Allison et al., 2005; Guo et al., 2019; Hu et al., 2020; Y. Guo et al., 2021). In a previous study, soil and root-associated arbuscular mycorrhizal fungi from prairie sites had higher richness and different community composition than those from adjacent crop soils (MacColl et al., 2024). However, other published data are not conclusive and evidence a complex response of soil communities to grassland restoration, that could depend on factors such as soil type and management (Smith et al., 2003; Bach et al., 2010; Strickland et al., 2017). Furthermore, the number of studies comparing soil microbiota from crops and adjacent restored prairies with metabarcoding approaches is limited, and little attention has been paid to larger organisms such as soil protists (Glaser et al., 2015; Santos et al., 2020; Romdhane et al., 2022). Because of the vast diversity and trophic modes comprised by soil protists, these organisms are highly relevant for the soil food web, biogeochemical cycling, microbial population regulation and plant health (Geisen et al., 2018). In recent years, high-throughput DNA sequencing tools have led to an improved knowledge of the diversity and function of soil biota, included these less studied groups (Burki et al., 2021). However, there are still many unanswered questions, including how these organisms respond to land use change and ecosystem restoration (Cotterill et al., 2008; Geisen et al., 2018).

Here, we assessed if restoration of marginal croplands to native prairie increased the diversity and/or altered the composition of soil prokaryotic, fungal and protist communities (hereby collectively referred to as “microbial communities” or “microbiota”, for simplicity). This was achieved through an observational study comparing soils from 5–9-year-old restored tallgrass prairie sites to those from adjacent croplands in different productive farms from southwestern Ontario, Canada. Soil microbial communities were studied using DNA high-throughput amplicon sequencing (metabarcoding) to analyze changes in their alpha and beta diversity, taxonomic composition and functional groups (fungi and protists only). Finally, using interkingdom network analyses (here, across the three studied microbial groups), we evaluated effects of prairie restoration on microbe-microbe associations within and across the three evaluated groups. Because prairie restoration increases plant cover and plant diversity (*i.e.*, more diverse inputs and root architectures), we expected soil microbial communities to be positively affected by this practice, leading to increased abundance and diversity. Changes in plant species composition, together with differences in the resulting edaphic environment, could also lead to clear shifts in microbial community composition after 5-9 years of restoration. The different microbial communities (prokaryotes, fungi and protists), encompassing different sizes, lifestyles and trophic modes, were expected to respond differently to the imposed environmental changes. Similarly, within each community, specific taxonomic and functional groups could differ in their sensitivity to these changes. Finally, since more diverse microbial communities could present more complex interaction networks (Wagg et al., 2019), this could be reflected in a higher number of microbe-microbe associations or correlations in prairie than crop soils.

## 2- Methods

### 2.1- Study area

This study was conducted on conventional farms from southwestern Ontario, Canada (Table 1, Figure S1), where retired agricultural fields had been restored to prairie grasslands as part of a Canadian farmer-led program named Alternative Land Use Services (ALUS, https://alus.ca/). The climate is temperate, with a mean annual temperatures of 8°C and average annual precipitation of 1036 mm (Dolezal et al., 2022; Mazzorato et al., 2022). The dominant soil types in the region span from sandy dominated luvisols to clay-dominated gleysols (Mazzorato et al., 2022). Farms were located along a SW-NE gradient, which was partially associated with differences in soil type and texture (*i.e.*, decreasing sand content towards the NE) (Table 1, Figure S1). All the agricultural sites were in a rotation of corn (*Zea mays*) and soybean (*Glycine max*), and planted with corn when sampled, except for Farm 2, which agricultural site was a fallow field after a sorghum (*Sorghum bicolor*) crop (Table 1). These sites received a ‘conventional’ input-intensive management, with tillage, application of synthetic fertilizers and use of pesticides. Prairie sites had been established within the previous 5-9 years from the time of sampling, on retired agricultural land that had been under the same management as the neighbouring agricultural fields.

**Table 1.** Description of farms and crop and prairie sites from SE Ontario, Canada, where soil samples were collected.

| Farm id. | Orig. id.* | Coordinates | Crop sites |  | Prairie sites |  |  |  |
| --- | --- | --- | --- | --- | --- | --- | --- | --- |
|  |  |  | Rotation | At sampling | Est. | Age (yrs) | Area (ha) | Dominant plant spp. |
| F1 | BO | 42.666 N, 80.704 W | Corn-Soybean | Corn | 2012 | 6 | 0.4 | Big bluestem, Golden rod |
| F2 | GI | 42.778 N, 80.682 W | Sorghum | Fallow | 2009 | 9 | 7.0 | Big bluestem, Japanese brome |
| F3 | CS | 42.878 N, 80.543 W | Corn-Soybean | Corn | 2009 | 9 | 2.4 | Big bluestem |
| F4 | VE | 42.877 N, 80.499 W | Corn-Soybean | Corn | 2013 | 5 | 1.0 | Switchgrass |
| F5 | BU | 42.891 N, 80.262 W | Corn-Soybean | Corn | 2010 | 8 | 5.0 | Big bluestem, Golden rod |

\* As published in MacColl *et al.*, 2024
Corn = *Zea mays* L., Soybean = *Glycine max* L., Sorghum = *Sorghum bicolor*
Big bluestem = *Andropogon gerardii*, Golden rod = *Solidago canadensis*, Japanese brome = *Bromus japonicus*, Switchgrass = *Panicum virgatum*
Crop site in F2 was in fallow but had *Digitaria* sp.

### 2.2- Soil sampling and physicochemical analyses

The collection of all soil samples was carried out during the summer, in late June 2018, within a period of 5 days to avoid biases due to seasonal variation in soil biota. In each of the five farm sites, we sampled two sites under different cover types, one under agricultural use and the other one with a restored prairie. For each of the sites, we randomly selected 5-m^2^ plots where we aseptically collected 12 soil cores (0-15 cm depth, 2 cm diameter). Soil cores within each plot were pooled and homogenized, resulting in a total of 50 soil samples (10 per farm, 5 per site). Samples were transported in coolers and, once in the lab, sub-samples for DNA analysis (∼0.25 g) were stored at –20°C until extraction. Another fraction of the soil samples was kept at 4°C for physicochemical analyses. Soil pH (water 1:2.5 v/v), textural fractions (laser diffraction method) were analyzed by A&L Canada Laboratories Inc., and organic matter (OM, Walkley-Black), extractable phosphorus (Olsen) and mineral nitrogen (ammonium plus nitrate, extracted with KCl) were analyzed by SGS Agri-Labs (Guelph, ON, Canada).

### 2.3- Soil DNA extraction, quantitative PCR and high-throughput sequencing

DNA extractions were carried out on ∼0.25 g equivalent dry soil samples using DNeasy PowerSoil™ Kit (Qiagen®, Valencia, CA) according to the manufacturer’s instructions. The concentration and quality of the extracted DNA was analyzed with NanoDrop 8000 (Thermo Scientific™) and 1% agarose gel electrophoresis.

Soil DNA samples were used to estimate bacteria and fungal abundance using real-time quantitative PCR (qPCR), as described in a previous work. Bacteria were quantified targeting the 16S rRNA gene with the 338F-518R primer set (Fierer and Jackson, 2005), whereas for fungi we targeted the 18S rRNA with the FR1-FF390 primer set (Vainio and Hantula, 2000). Amplification and quantification details are available in Tosi *et al*. (2022). Data were presented as gene copies per gram dry soil, as well as the fungi-to-bacteria ratio.

With the goal of evaluating microbial community composition and diversity, soil DNA was analyzed using high-throughput sequencing (HTS) at the Génome Québec (Montréal, Québec) facility. To target prokaryotes, we sequenced the V4 region of the 16S rRNA gene using primers 515F-Y (5’-GTGYCAGCMGCCGCGGTAA-3’) (Parada *et al*., 2016) and 806R (5’-GGACTACNVGGGTWTCTAAT-3’) (Apprill *et al*., 2015). Fungal communities were amplified via the first internal transcribed spacer (ITS1) of the ribosomal DNA using the primers ITS1f (5’-CTTGGTCATTTAGAGGAAGTAA-3’) and ITS2 (5’-GCTGCGTTCTTCATCGATGC-3’) (White *et al*., 1990). Protist communities were targeted by amplifying the 18S rRNA V4 region with primers 616*F (5’-TTAAARVGYTCGTAGTYG-3’) and 1132R (5’-CCGTCAATTHCTTYAART-3’) (Hugerth et al., 2014). The 16S rRNA and ITS primer pairs used here are those recommended by the Earth Microbiome Project, while the 18S rRNA protist primers had the best overall performance for the V4 region of the 18S rRNA (Vaulot et al., 2022). Amplicon sequencing was carried out on an Illumina MiSeq platform with a read length of 2 x 250 bp (PE250) for prokaryotes and fungi, and 2 x 300 bp (PE300) for protists. In all cases, amplification conditions were optimized by the sequencing facility, following protocols by the Earth Microbiome Project for 16S rRNA and ITS, and by Hugerth *et al*. (2014) for protists. Raw sequencing data was deposited under BioProject ID PRJNA1170895 in the Sequence Read Archive (SRA) of the National Centre for Biotechnology Information (NCBI).

### 2.4- Bioinformatics

Demultiplexed paired-end sequences obtained from the sequencing facility were analyzed using the bioinformatics platform QIIME 2 2023.2 (Bolyen et al., 2019). Before denoising, primer sequences of prokaryotic 16S rRNA reads were removed using q2-cutadapt (Martin, 2011). In the case of fungal reads, we used the ITSxpress plugin to trim off the conserved flanking regions of SSU and 5.8S which are comprised within the sequenced ITS fragment (Rivers *et al*., 018). The QIIME 2 plugin for DADA2 (q2-dada2) (Callahan *et al*., 2016) was used for denoising, dereplication, chimera filtering and merging, including a step to truncate the end of reads at a length that would allow to remove low quality fragments while allowing reads to merge. To assign taxonomy to the obtained amplicon sequence variants (ASVs) we used the plugin q2-feature-classifier (Bokulich et al., 2018), first training classifiers with fit-classifier-sklearn and then carrying out the taxonomic classification with classify-sklearn (Pedregosa et al., 2011). Taxonomic classifiers were trained using the databases Silva v. 138.1 (Quast *et al*., 2013) with 99% OTUs for prokaryotes and UNITE v. 9 with dynamic use of clustering thresholds for fungi (Abarenkov *et al*., 2020).

Protist data was processed by previously concatenating or joining length-trimmed paired-end reads to maximize the recovery of reads and taxa. Even though the obtained 18S rRNA fragment has been previously analyzed with merged paired-end reads (Oliverio et al., 2020), this fragment size can surpass the Illumina MiSeq PE300 limit for several protist groups, including some common soil taxa (*e.g.*, Amoebozoa, Excavata and Opisthokonta) (Vaulot et al., 2022). Similar concerns were recently raised by Mau et al. (2024). Our approach was chosen after comparing the performance of different suggested approaches (Dacey and Chain, 2021) (more details in Supplementary Methods). Briefly, after removing primers and length-trimming reads with cutadapt (Martin, 2011), reads were concatenated following protocols by Dacey and Chain (2021). These reads were then imported into QIIME2 and then input into q2-dada2 (Callahan *et al*., 2016) for denoising, dereplication and chimera filtering of reads. After this step, we obtained a total of 656,512 reads and 9645 ASVs before removing non-target sequences. Taxonomy was assigned using q2-feature-classifier classify-sklearn (Bokulich *et al*., 2018) with a taxonomic classifier trained using the PR^2^ (Protist Ribosomal Reference) database v.5.0.0 (Vaulot et al., 2022) (more details in Supplementary Methods). Non-target reads, only found in 16S rRNA (*i.e.*, chloroplast and mitochondria) and 18S rRNA sequencing, were removed before carrying out downstream analyses.

### 2.5- Functional classification of fungal and protist taxa

Functional groups were assigned to fungal and protist taxa only, as prokaryotes have no database that is both simplified and robust enough for these purposes. In the case of fungi, genera were assigned a primary and a secondary lifestyle using the FungalTraits database (Põlme et al., 2020). Protist genera were classified functionally using the database developed and applied by Xiong *et al*. (2021), followed by other resources whenever information was unavailable (Adl and Gupta, 2006; Adl et al., 2018; Schulz et al., 2019; Mazel et al., 2022). This classification includes the following categories: phagotroph, phototroph, parasite, mixotroph, saprotroph, plant pathogen and dinoflagellate. The phagotroph category comprises all protists consuming other organisms (bacteria, fungi, algae, other protists, animals) and sometimes small organic matter particles. Out of the 376 taxa (genera or lowest level available) detected in our dataset, we were able to assign function to 324 (86.2%) and probable function to 11 (2.9%). The remaining 41 taxa (10.9%) could not be assigned a trophic group either because it was not available or because taxonomic classification was partial.

### 2.6- Data analysis

Data was analyzed using QIIME 2 2023.2 and R v. 4.2.2 (R Core Team, 2022). Diversity analyses were carried out both at the ASV and genus level (here, genetic or fine-scale and taxonomic or coarse-scale), whereas differentially abundant taxa were explored at the genus and phylum level (or, in the case of protists, level 3 from the PR^2^ database). It is worth noting that in some cases ASVs cannot be classified up to the genus level, so these are clustered at the lowest taxonomic level available. These are counted as a single genus even though they could comprise multiple genera. Diversity metrics for ASVs and genera were calculated in QIIME 2 and R package ‘vegan’ (Oksanen et al., 2022), respectively. The ASV tables used to calculate alpha and beta diversity metrics (and to create the genus level table for the same purpose) were filtered to remove ASVs with very low abundance (<10 reads) and/or very low prevalence (<2 samples), and then rarefied to an even number of reads.

For alpha diversity, we calculated richness, evenness (Pielou’s index) and Shannon diversity index. Due to the highly contrasting responses between farm sites, we analyzed both land use and farm site effects, as well as their interaction, using ANOVA with generalized least squares (*gls*) in R package ‘nlme’ (Pinheiro et al., 2018). In each model, residuals were checked for normality and homoscedasticity. The same approach was used to analyze microbial abundance data (log gene copies g^−1^ dry soil) from qPCR.

To evaluate changes in genetic and taxonomic composition and beta diversity, we used two distance metrics: Jaccard (qualitative, based presence-absence) and Bray-Curtis (quantitative, based on relative abundance), with a focus on the former. Multivariate homogeneity of groups dispersions between land uses or farm sites was tested using *betadisper* in R package ‘vegan’. Land use, farm site and their interaction effects on community composition were tested using permutational multivariate analysis of variance (PERMANOVA) with function *adonis2* in R package ‘vegan’. Because interactions between land use and farm site were significant, we also carried out separate PERMANOVAs for each farm site. Distances between sites were visualized using principal coordinate analyses (PCoA), as well as plotting prairie-to-crop, within-site and between-farm distances. Venn diagrams from InteractiVenn (http://www.interactivenn.net/) were used to explore unique and shared ASVs and genera between land uses, and the multinomial species classification method (CLAM), with ‘vegan’ function *clamtest*, was used to explore potentially generalist and specialist ASVs and genera in each land use. To explore how restoration drives microbial compositional changes, we tested two scenarios: prairies either gain new ASVs while replacing existing ones or without replacing them. To test this, we partitioned beta diversity between crop and prairie microbial communities into turnover and nestedness (Baselga, 2012). High turnover supports the replacement scenario, while high nestedness suggests ASVs form a nested subset, indicating no replacement.

Changes in the relative abundance of taxonomic groups were analyzed at the phylum level and the genus level using differential abundance analyses, as they take into consideration the compositional nature of sequencing data (Gloor et al., 2017). Based on a comparative study by Nearing *et al*. (2022), differentially abundant taxa between crop and prairie soils were selected according to two reliable methods for compositional and sparse data: Analysis of Compositions of Microbiomes with Bias Correction (ANCOM-BC) (Lin and Peddada, 2020) and ANOVA-Like Differential Expression (ALDEx2) (Fernandes et al., 2013). ANCOM-BC was run with q2-composition in QIIME2 and ALDEx2 with function *aldex.t* in R package ‘ALDEx2’. Only taxa detected by both methods were considered to change significantly between land uses.

Relationships between the three microbial groups, and between them and soil physicochemical properties, were analyzed out with different approaches and in different data subsets (complete dataset, crop sites and prairie sites). Correlation coefficients were calculated to relate alpha diversity indices between microbial groups or with soil properties. Similarly, we used Mantel tests to evaluate correlations between community composition of the three different groups. To evaluate the contribution of farm, land use, edaphic and vegetation effects to the variability in microbial community composition, we carried out a variance partitioning analysis using the function *varpart* in ‘vegan’. We ran three separate analyses: one testing the contribution of soil physicochemical properties, land use and farm on all sites, and two separate analyses for prairie and crop sites evaluating the effects of soil properties, farm and plant species composition. Plant species composition effects were excluded from whole dataset analysis to avoid a high collinearity error, and from the crop sites analysis due to the high similarity between crop sites. Plant species composition data was introduced using correspondence analysis (CA) axes obtained with *cca* function in ‘vegan’. Seven CA axes were included, in order to match the number of soil variables (sand, clay, pH, OM, mineral N, extractable P and the C:P ratio). Finally, to test the influence of different soil physicochemical properties on community composition, we used the function best subset of environmental variables or *bioenv* in ‘vegan’. This analysis was used to find the subset of soil physicochemical variables with the maximum rank correlation with community dissimilarity matrices for each microbial group. Mantel tests, variance partitioning and *bioenv* were carried out on both Jaccard and Bray-Curtis distance matrices at the ASV level only.

Trans-kingdom association networks for each land use category were built with SPIEC-EASI (Sparse InversE Covariance estimation for Ecological Association and Statistical Inference) (Kurtz et al., 2015), as recommended by Matchado *et al*. (2021). SPIEC-EASI employs the CLR transformation to address compositionality and relies on a graphical model inference framework that presupposes sparsity in the underlying network (Kurtz et al., 2015). These analyses were carried out in the R package ‘SpiecEasi’ (Kurtz et al., 2023), using the *spiec.easi* function, which now includes a wrapper to work with multiple sequencing datasets from different taxa. We used the neighborhood selection framework (MB method) (Meinshausen and Bühlmann, 2006) with parameters: an nlambda (number of regularization levels) of 100, a lambda minimum ratio of 5 e^−1^, and a StARS criterion threshold of 0.01 in the pulsar parameters. We chose a conservative threshold to obtain sparse networks with fewer associations and less false positives, as our goal is to explore the most relevant microbe-microbe associations. The analysis was carried out using genus level tables, previously filtered by total abundance (min = 10 reads) and prevalence (min = 7 samples) within each dataset (land use and microbial group). Association networks were visualized and analyzed using Gephi v. 0.10. We compared parameters such as number of nodes and edges, nodes and edges in common, taxonomic groups involved in these interactions, as well as other descriptive network parameters. We also explored influential nodes or taxa (*i.e.*, ‘keystone’ taxa) based on their eigenvector centrality, which takes into account the number of edges or associations as well as quality of these edges, in terms of the importance of those neighboring nodes (Ruhnau, 2000; Obregon et al., 2023).

## 3- Results

### 3.1- Soil physicochemical properties

The studied farms presented a SW to NE soil texture gradient, with decreasing sand content from F1 to F4-F5 (Table 2, Figure S1). Land use effects on soil physicochemical properties varied across farms, with the exception of mineral N, which was consistently lower in restored prairie than crop soils (Table 2). One of the farms, F2, presented the lowest mineral N values (Table 2). Besides mineral N, overall land use effects were observed only for soil OM and pH, although among-farm variation was evident for these two variables (Table 2, S1). Texture fractions and extractable P showed only farm-specific land use effects (Table 2). One of the most distinctive farms was F5, where restored prairie soils had markedly lower soil OM, higher clay content, and higher extractable P (Table 2). Farm 4 was the only one where crop and prairie soils differed in texture, with the latter presenting higher sand and lower fine particles content (Table 2).

**Table 2.**
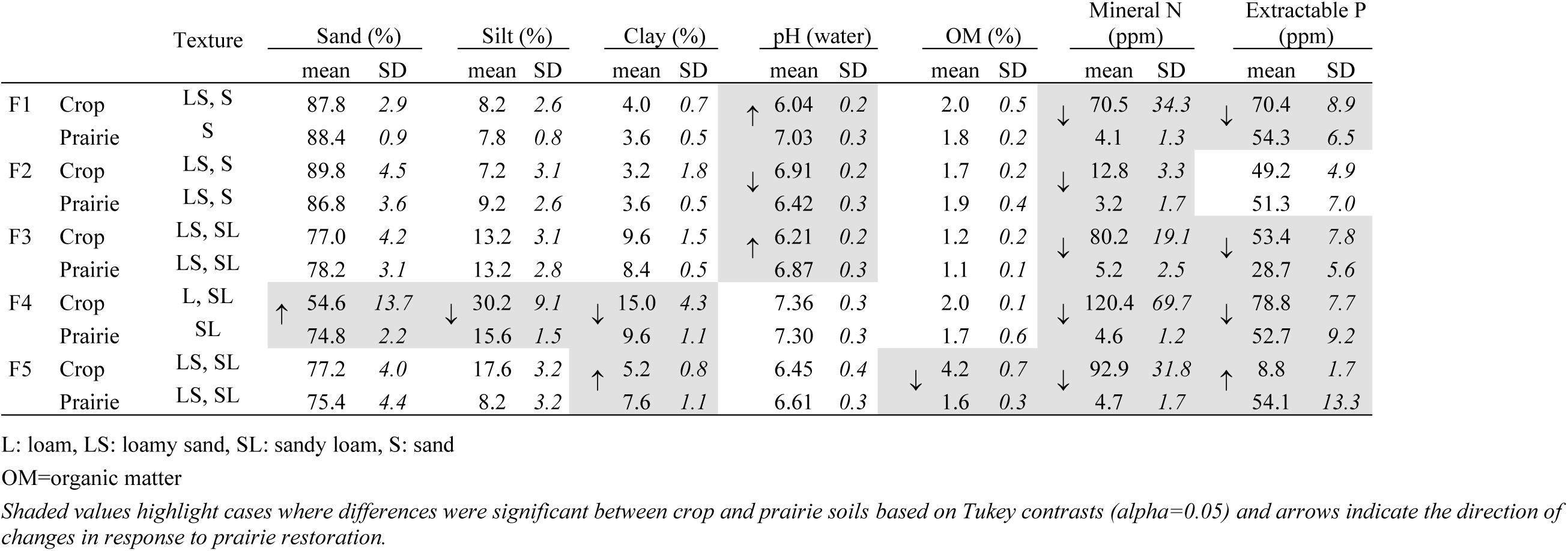
Soil physicochemical properties across farms and land uses. Mean and standard deviation (SD) values are shown.

### 3.2- Microbial abundance and alpha diversity

Soil bacterial abundance did not show a clear response to prairie restoration, except for two farms (F1 and F3) where it was higher in restored prairie sites compared to crop sites (Figure 1A, Table S1). In contrast, prairie restoration led to an average decrease in soil fungal abundance (from 8.5 × 10^7 to 6.1 × 10^7 copies g^−1^ dry soil in crop and prairie sites, respectively) and the fungi-to-bacteria ratio (F:B ratio) (Figure 1B, 1C, Table S1). However, when looking at results by farm, this difference was only significant in F2, where restored prairie soils had a 49% reduction in fungal abundance relative to crop soils (Figure 1B, 1C).

**Figure 1.**
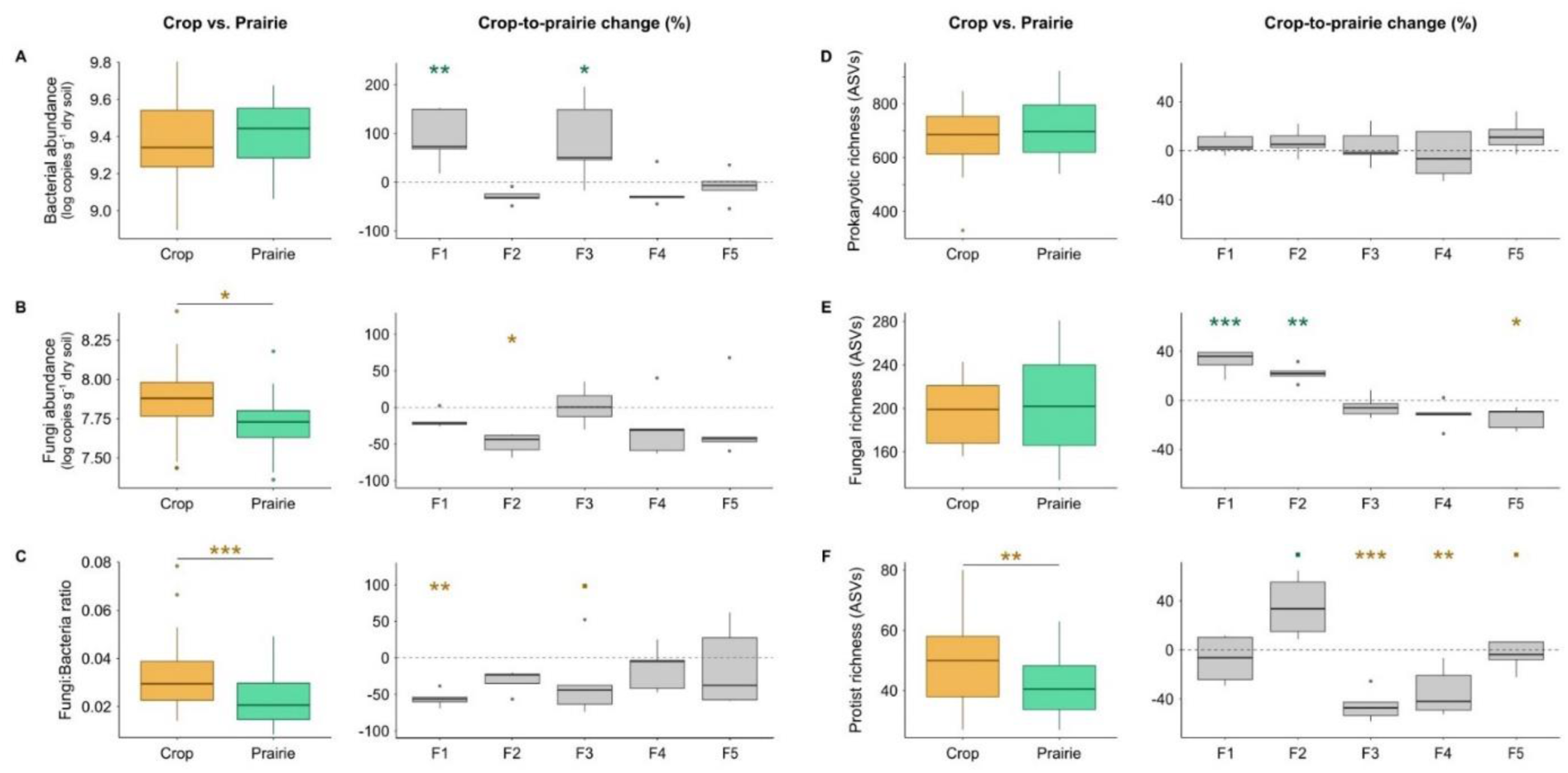
Response of soil microbial abundance from quantitative PCR and microbial genetic richness from high-throughput sequencing to prairie restoration. A-C) Soil bacterial abundance, fungal abundance and fungi:bacteria ratio. D-F) Genetic richness (ASV level) of soil prokaryotic, fungal and protists. For each variable, average values per land use and percentage change from crop to prairie sites are shown. *Crop-to-prairie change was calculated from each prairie plot relative to the crop site average of 5 plots (positive=higher in prairie, negative=higher in crop). In average value plots, asterisks represent significant changes based on ANOVA, whereas in percentage change plots, asterisks represent significant differences between prairie and crop soils based on Tukey contrasts (alpha=0.05)*.

After processing sequencing data as described in the methods, we recovered a total of 1,197,342 reads and 17,725 ASVs for prokaryotes, 2,612,111 reads and 5,411 ASVs for fungi, and 83,591 reads and 2,225 ASVs for protists (Table S2). Very low abundance and prevalence ASVs (<10 reads and <2 samples), which were removed for downstream analyses, comprised 66-76% of the total number of ASVs detected, but only 10-24% of the total reads (Table S2). When comparing restored prairie grasslands with conventional cropping systems, we did not find consistent changes in the alpha diversity of soil microbial prokaryotic, fungal and protist communities. Instead, the direction of changes strongly depended on the microbial group analyzed and the farm.

Prokaryotic genetic richness (ASVs) did not significantly differ between land uses, and this was consistent across farms (Figure 1D, Table S1). By contrast, taxonomic richness was on average slightly higher on crop than restored prairie soils (265 vs. 257, respectively, P=0.04), although tests by farm were not significant (Figure S3A). The crop-to-prairie changes in fungal genetic and taxonomic richness showed a pattern based on the geographic location and soil type of the farms (Figure 1E, Figure S3B, Table S1). The response of fungal genetic richness to prairie restoration was positive in F1 and F2 (32.6 and 21.9% increase, respectively, P<0.01), neutral in F3 and F4, and negative in F5 (–14.3%, P=0.019) (Figure 1E, Figure S3B). Protist genetic richness was on average lower in restored prairie sites than crop sites (41 vs. 49, respectively, P=0.005), but this response was only clearly detected in farms F3 and F4 (–45% and –34% change, respectively, P<0.01), while one of them (F2) showed the opposite trend (35% increase, P=0.066) (Figure 1F, Table S1). No changes in protist taxonomic richness were observed (Figure S3C). A complementary analysis examining individual phyla suggests that changes in richness could be strong within specific groups as opposed to the overall prokaryotic, fungal or protist community (Figure S4). For example, according to this exploration, prairie restoration could be increasing microbial richness within Acidobacteria and Basidiomycota, while decreasing it for Actinobacteria, Chloroflexi and Proteobacteria (Figure S4). Alpha diversity in terms of evenness was only sensitive for soil prokaryotic communities, showing a minor negative response to prairie restoration: genetic at F1 (–1.9%, P=0.013), and taxonomic at F1-F3 (–4.3, –2.9 and –2.9% change, respectively, P<0.01) (Figure S5).

We did not find any significant correlations between alpha indices from different groups, analyzing either the whole dataset, only crop sites or only restored prairie sites (data not shown). The distinct behaviour of alpha diversity across microbial groups is summarized in the PCA biplot of Figure S6. Alpha diversity metrics from different microbial groups were not correlated either, for the most part (data not shown). The only exception was a positive correlation between fungal evenness and protist richness, only amongst crop soils (Spearman r=0.41, P<0.05).

### 3.3- Beta diversity and composition shifts

For all microbial groups, we observed consistent shifts in genetic and taxonomic community composition in response to both land use and farm-specific factors, with an interaction between these variables (Figure 2, Table S3). Still, significant land use effects on microbial community composition were observed across all farms (Table S4), although with differences in prairie-to-crop distances among them (Figure 2A-2C, Figure S7). For example, crop communities in F5 and F2 sometimes clustered more closely with prairie communities than with other crop communities (Figure 2A-2C). Based on PERMANOVA R-squared values, land use explained a similar proportion of variance in genetic community composition for all microbial groups (4.6% for prokaryotes, 5.8% for fungi and 4.8% for protists), whereas farm effects explained 15.3, 18.3 and 21% for protists, prokaryotes and fungi, respectively (Figure 2G, Table S3). Results using Bray-Curtis dissimilarities (quantitative) were similar to those found using Jaccard distances (presence-absence), although with higher R-squared values (Figure 2G, Table S3).

**Figure 2.**
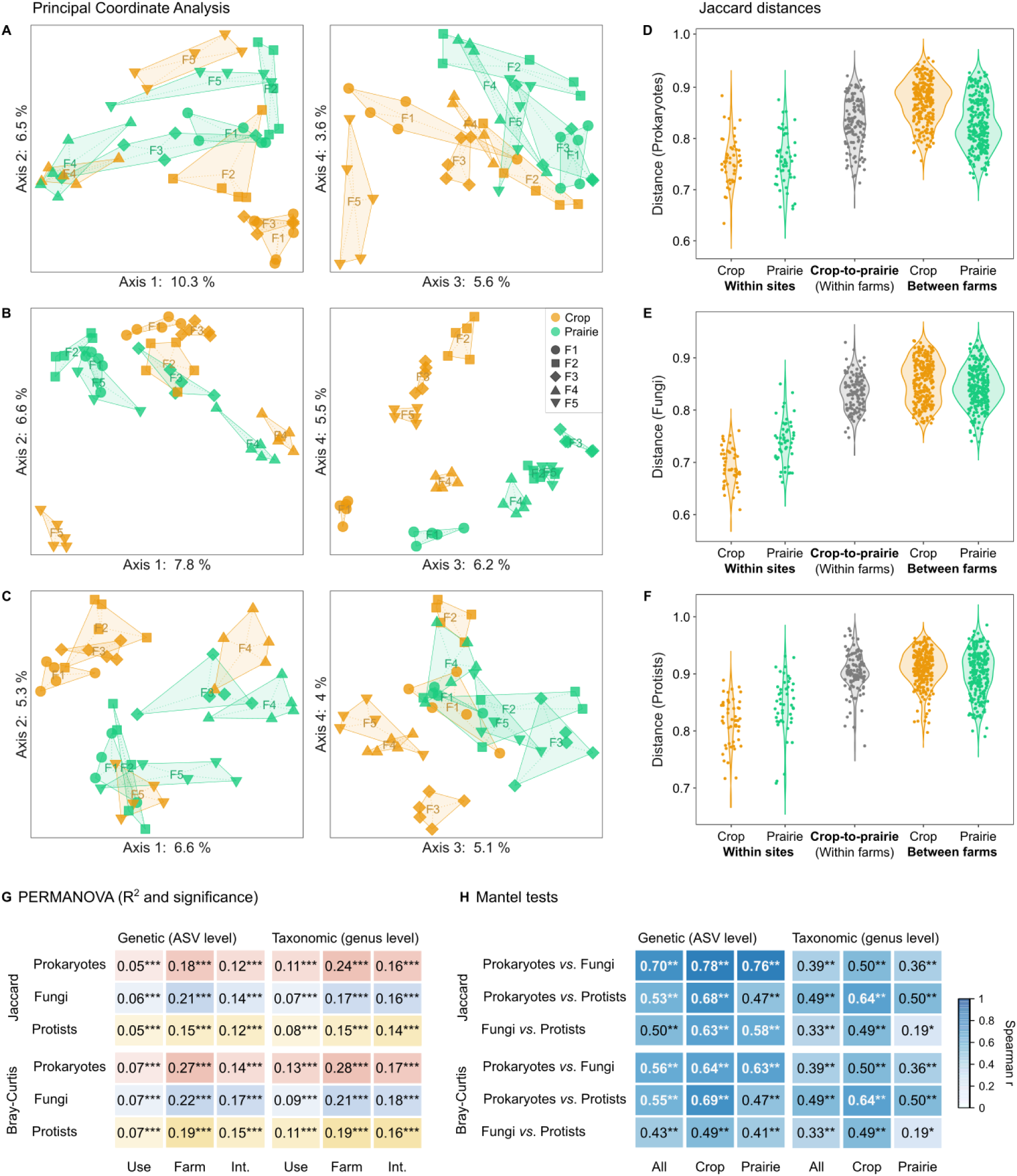
Response of measured soil microbial beta diversity (prokaryotes, fungi and protists) to prairie restoration. A-C) Principal Coordinate Analysis (PCoA) plots of Jaccard distances between sites for the first 4 axes (genetic or ASV level), D-F) Jaccard distances within crop and prairie sites, between crop and prairie sites (within each farm) and between crop and prairie soils from different farms (genetic or ASV level), G,H) Summary of PERMANOVA and Mantel test results for different microbial groups using two distances (Jaccard and Bray-Curtis) and two scales (genetic and taxonomic). In A-C, convex hulls enclose the 5 plots for each farm-use combination. Full PERMANOVA results are shown in Table S3.

In all microbial groups, distances within sites were always smaller than crop-to-restored prairie distances (within each farm) and distances between farms (within each land use) (Figure 2D-2F). All distances were overall higher for prokaryotes than for fungi or protists, although substantial variability was observed for all communities (Figure 2D-2F). For fungal and protist communities, within-site distances were smaller in crop than restored prairie communities (Figure 2E, 2F). On the other hand, for prokaryotic communities, between-farm distances were slightly larger among crop than restored prairie sites (Figure 2D), in agreement with *betadisper* test results (Table 3). Mantel tests found significant correlations between all three communities in terms of genetic and taxonomic community composition (Figure 2H). For genetic composition, the strongest correlations were found between prokaryotic and fungal communities, and crop sites showed higher correlations than restored prairie sites or the complete dataset (Figure 2H). On the other hand, the highest taxonomic composition correlations were between prokaryotic and protist communities, particularly amongst crop sites, and the lowest between fungal and protist communities, particularly amongst restored prairie sites (Figure 2H).

**Table 3.**
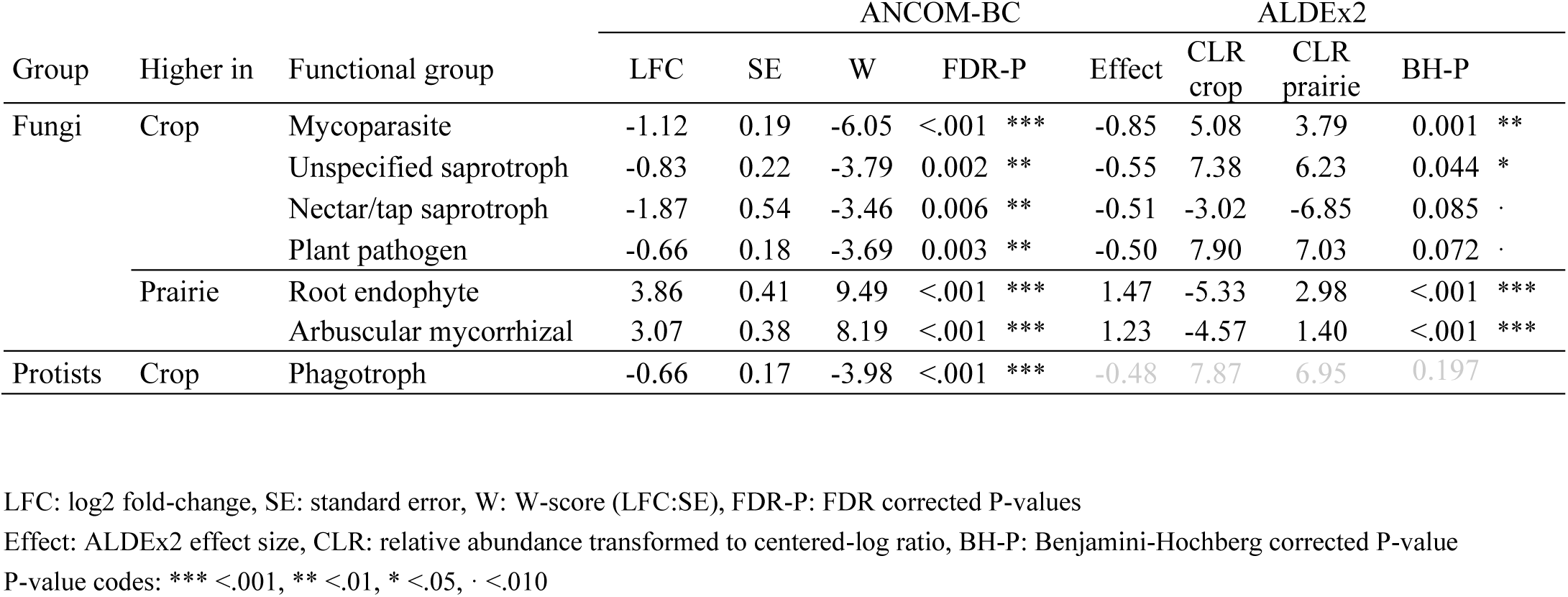
Changes in fungal lifestyles and protist trophic modes between land uses (crop vs. prairie) based on two differential abundance analyses (ANCOM-B and ALDEx2). *Only groups with corrected P-value < 0.10 are shown*.

For all microbial groups, genetic or ASV turnover in restored prairie relative to crop sites was higher and nestedness was lower than expected by chance, although with some farm-specific differences (Table S5). Prokaryotes and protists had higher ASV turnover only in F1, F3 and F4, whereas fungi had higher ASV turnover at all farm sites except F1. The higher turnover was consistent with the small proportion of shared ASVs between crop and restored prairie sites (Figure 3A). It is worth noting that the proportion of shared features was only small at the genetic level (14-23%) but increased markedly when looking at genera (67-80%) (Figure 3A, 3B). Moreover, both shared ASVs and genera were higher for prokaryotes, followed by fungi and protists. Crop sites had a higher number of unique and specialist protist and prokaryotic features than restored prairie sites, whereas the opposite was observed for fungi (Figure 3A-D). However, a high proportion of these unique features had very low prevalence, as can be seen when filtering out those present in less than 5 plots (Figure 3A, 3B). Most crop and restored prairie ‘specialists’ belonged to a few taxonomic groups: Proteobacteria (prokaryotes), Ascomycota (fungi), and Chlorophyta and Rhizaria (protists). Yet, notably, some taxa were only found among crop (*i.e.*, Crenarchaeota, Firmicutes and Myxococcota, Blastocladiomycota, Tubulinea) or restored prairie (*i.e.*, Acidobacteria, Bacteroidota and Desulfobacterota, Glomeromycota, Alveolata) ‘specialists’ (Figure 3C-3D, Figure S5).

**Figure 3.**
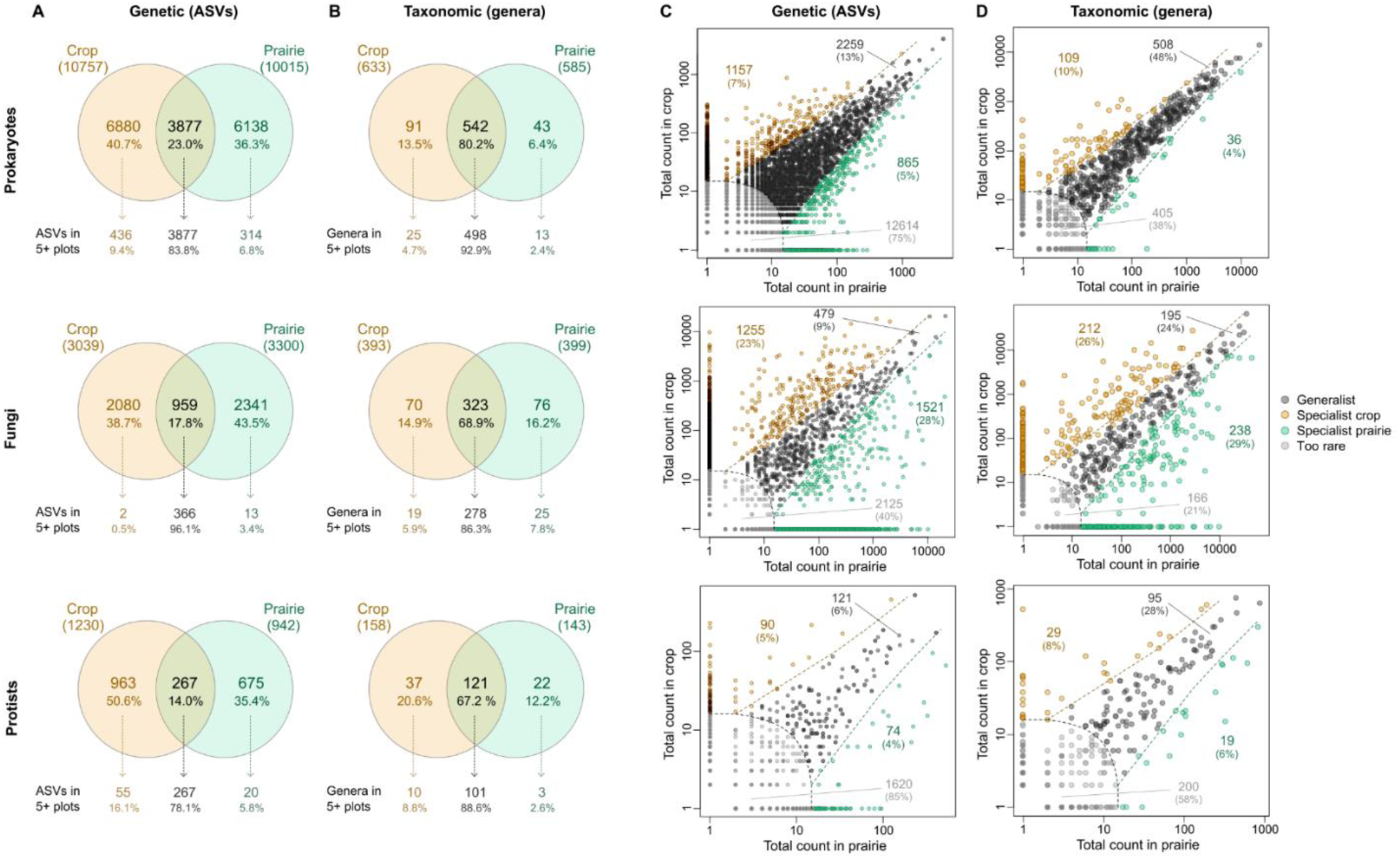
Distribution of soil microbial ASVs and genera (genetic or fine-scale and taxonomic or coarse-scale, respectively) across land uses. A) Venn diagrams showing shared and unique ASVs and genera by land use (including all features and only those present in 5 or more plots/samples). B) Analysis of specialist-generalist genera based on the multinomial species classification method (CLAM test). Nestedness analysis can be seen on Table S4.

### 3.4- Relative abundance of specific microbial taxa

In soil prokaryotic communities, three bacterial phyla comprised 64.8% of the total reads: Proteobacteria (24.8%), Actinobacteria (21.3%) and Acidobacteria (18.7%), with the latter being more dominant in restored prairie soils (Figure 4A). The dominant fungal phylum was Ascomycota (77.4%), especially in crop soils, followed by Mortierellomycota (10.0%) and Basidiomycota (6.6%) (Figure 4A). Lastly, soil protist communities were dominated by TSAR;Rhizaria (37.3%), particularly in restored prairie soils, followed by Amoebozoa;Evosea (24.8%) and Archaeplastida;Chlorophyta (21.3%) (Figure 4A). According to our differential abundance analyses, prairie restoration increased three bacterial phyla (Latescibacterota, Desulfobacterota, Acidobacteriota) and the fungal phylum Glomeromycota, but no major protist groups (Figure 4B). Methylmirabilota and Planctomycetota increases in restored prairie soils were only detected by ANCOM-BC (Figure 4B). On the other hand, the relative abundance of Deinococcota (Bacteria), Chytridiomycota (Fungi), and Amoebozoa_X (Protists) was decreased in restored prairie soils relative to crop soils (Figure 4B). ANCOM-BC also detected decreases in the archaeal Crenarchaeota, three fungal phyla (Ascomycota, Mucoromycota and Blastocladiomycota) and three protist groups (Amoebozoa;Evosea, Amoebozoa;Tubulinea, and Excavata;Discoba) (Figure 4B).

**Figure 4.**
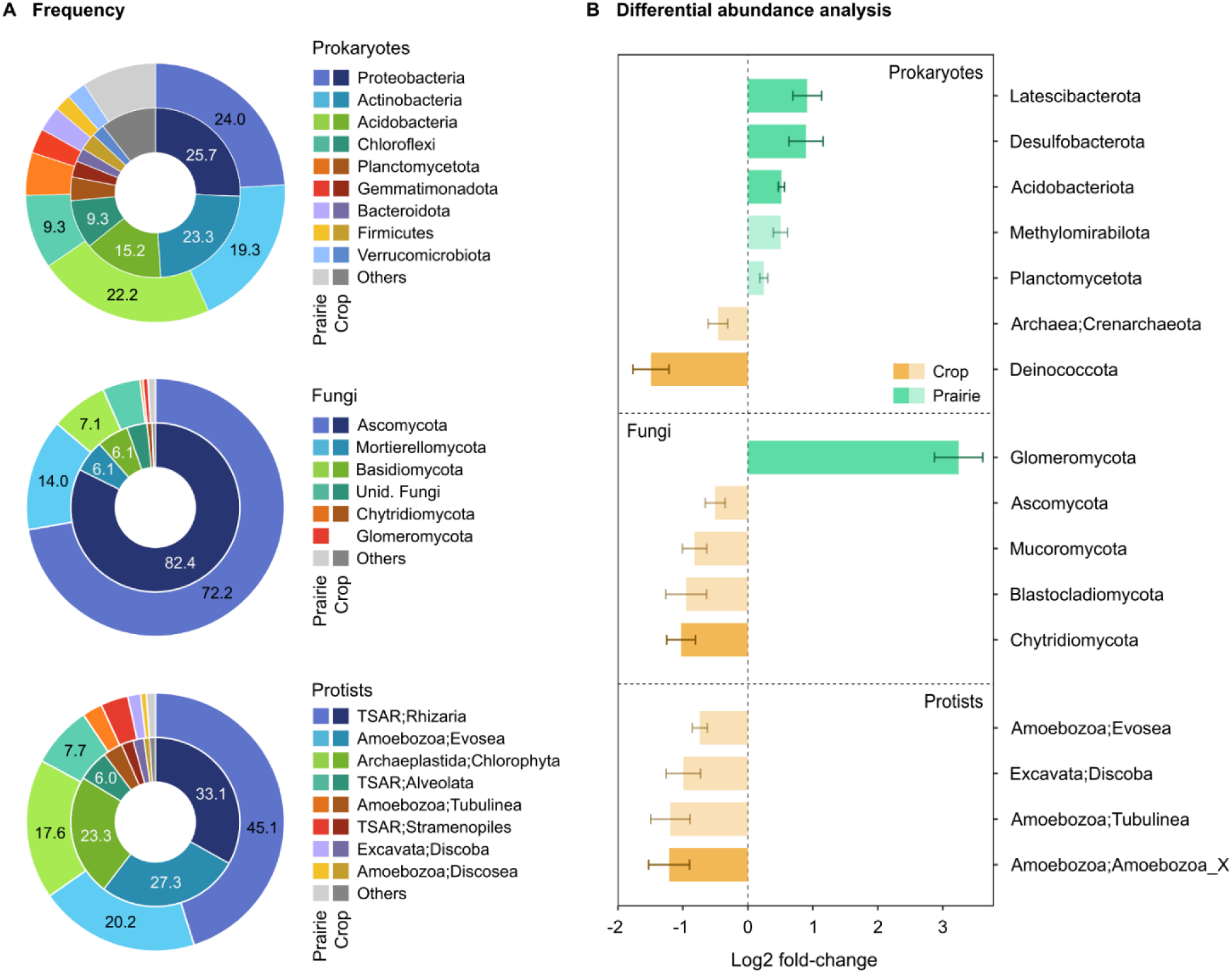
Changes in soil microbial phyla between crop soils and adjacent restored prairie soils. A) Overall frequency (%) of prokaryotic and fungal phyla and major protist groups in crop (inner donut) and prairie (outer donut) soils. B) Changes in relative abundance of microbial phyla or major protist groups between land uses (corrected P-value<0.05). In A, lower abundance groups are grouped in the “Others” category. In B, only sensitive taxa are shown, and light-shaded bars represent taxa only detected by ANCOM-BC, whereas darker-shaded bars represent taxa detected by both analyses (ANCOM-BC and ALDEx2).

At the genus level, there were a higher number of land use-sensitive taxa among fungi (34), followed by prokaryotes (32), and last by protists (11) (Figure 5A, 5B). Restored prairies favoured more prokaryotic genera than crop use, while the opposite was observed for fungi and protists (Figure 5A, 5B). In the case of prokaryotes, only 4 genera were promoted under crop systems, two of them ammonia oxidizers (*i.e.*, the archaeal *Candidatus Nitrocosmicus* and the bacterial *Nitrosospira*) (Figure 5B, Table S6). Of the 27 prokaryotic genera increased in restored prairie soils, most belonged to Proteobacteria (10) and Acidobacteria (9) (Figure 5B, Table S6). When looking at fungal communities, crop soils promoted 21 genera and restored prairie soils only 13, in both cases mostly from Ascomycota, followed by Basidiomycota (Figure 5B, Table S7). Crop soils also promoted two Chytridiomycota, whereas prairie restoration also promoted one Mortierellomycota. Finally, among protists we found 8 genera favoured under crop use and 3 under prairie restoration (Figure 4B, Table S7). More than half of the protist taxa favoured under crop belonged to Amoebozoa (2 Evosea, 2 Tubulinea and 1 Amoebozoa_X), whereas two of the groups promoted in restored prairie belonged to Vampyrellida (TSAR;Rhizaria;Cercozoa) and the third one to Excavata;Discoba (Figure 5B, Table S7).

**Figure 5.**
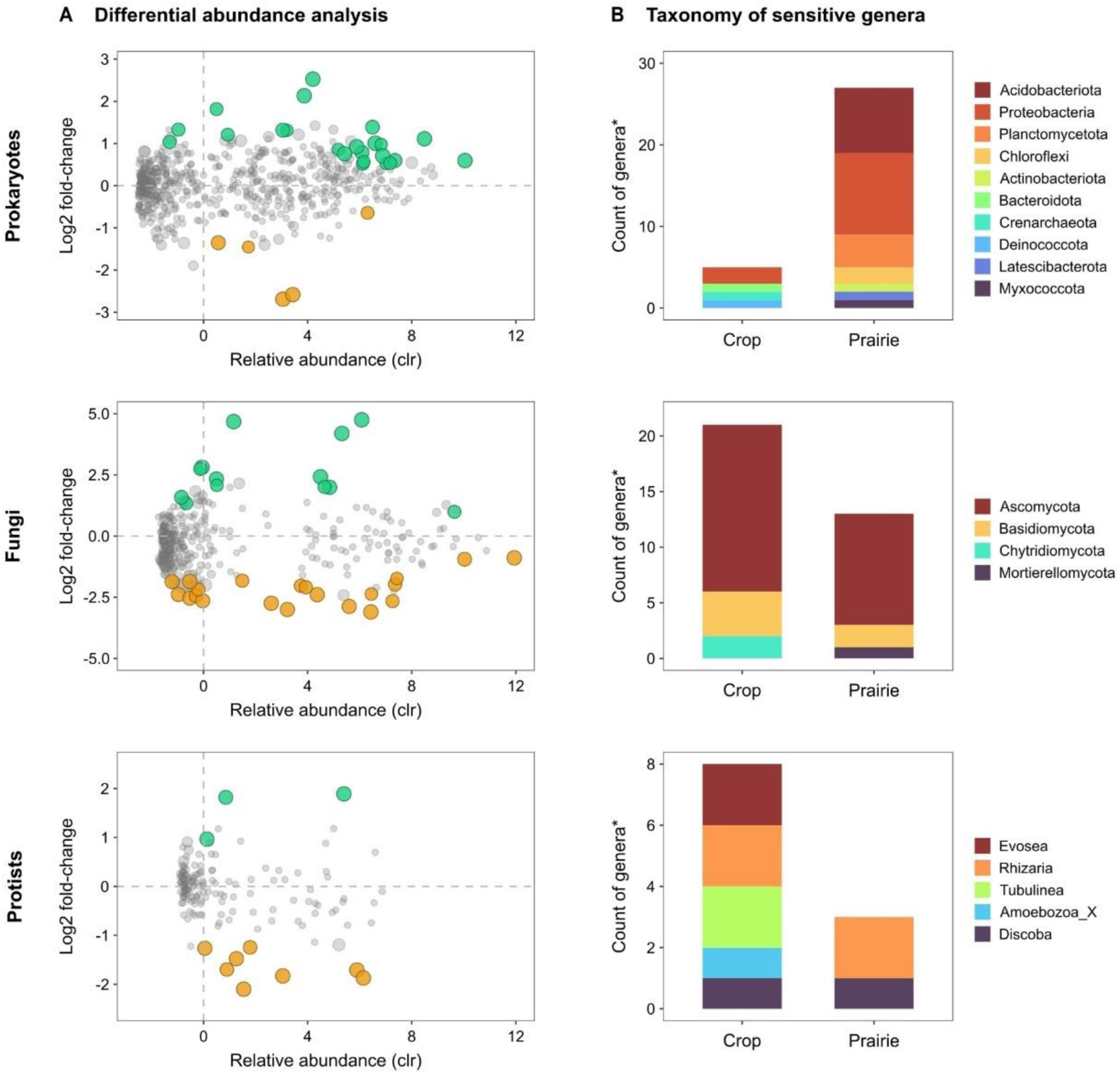
Changes in soil microbial genera between crop soils and adjacent restored prairie soils. A) Soil microbial genera showing changes as related to their relative abundance between land uses (from differential abundance analysis) and their overall relative abundance transformed to centered log ratio (clr). B) Taxonomy of sensitive genera from panel B as frequency of phyla or major protist groups (B). *Changes in relative abundance were assessed based on significance (corrected P-value<0.05) on two differential abundance analyses (ANCOM-BC and ALDEx2). Predicted function of sensitive fungi and protists can be found in Figure S5 and Tables S6 and S7*.

### 3.5- Predicted functional groups

The functional profile of fungal communities showed a dominance of saprotrophs in both crop and restored prairie soils (∼41.9 and 44.4%, respectively), although ∼34% of the taxa could not be assigned (Figure S7). Differential abundance tests found that restored prairie soils had higher relative abundance root endophytes and arbuscular mycorrhizal fungi, whereas crop soils had more mycoparasites and unspecified saprotrophs, plant pathogens and nectar/tap saprotrophs (Table 3). Among the fungal genera increasing in crop soils (see Figure 5), these were dominantly saprophytic, but also comprised 4 plant pathogens (*Fusarium*, *Bipolaris*, *Ophiosphaerella* and *Plectosphaerella*) and one mycoparasite (*Fusicolla*) (Figure S6, Table S7). Restored prairie soils also increased several saprophytic fungi, as well as one root endophyte (*Serendipita*) and one animal parasite (*Hirsutella*). Notably, only crops promoted dung saprotrophs (*Enterocarpus* and *Kernia*), while only restored prairies promoted wood saprotrophs (*Pyrenochaeta* and *Paraphaeosphaeria*).

Soil protist communities were markedly dominated by phagotrophs (∼70%), followed by phototrophs (∼20%) (Figure S7). Based only on ANCOM-BC, prairie restoration decreased the relative abundance of phagotrophs (Table 3). Besides this, protist functional groups responded more to farm effects than to land use, with farms in finer textures promoting phototrophs, especially in crop soils (Figure S6). All sensitive differentially abundant taxa (see Figure 5) were phagotrophs, regardless of whether they increased in crop or prairie use (Figure S7).

### 3.6- Interkingdom associations

Interkingdom network analyses revealed some differences between land uses in terms of the association between microbial taxa. Compared to restored prairie soils, crop soils presented a higher number of total nodes (469 vs. 513, respectively), especially prokaryotic, and unique nodes (154 vs. 198, respectively) (Figure 6A, 6B). Networks also had a higher number of edges, both negative and positive, in crop than restored prairie soils (Figure 6A). Consistently with the type of nodes, prokaryote-prokaryote associations were the most frequent, followed by prokaryote-fungus (Figure 6A). Notably, despite crop and restored prairie networks having 315 nodes in common (47.2%), only 8 edges occurred in both (Figure 6B): 6 bacteria-bacteria edges (positive: *Conexibacter*-*Terrabacter*, AD3-Acetobacteracea, AKAU4049-*Rubrobacter* and Pedosphaeraceae-Latescibacterota; negative: AD3-Vicinimibacteraceae, AKAU4049-*Burkholderia*_*Caballeronia*_*Paraburkholderia*), one negative bacterium-fungus (*Candidatus*_*Nitrocosmicus*-*Mariannaea*), and one positive fungus-fungus edge (*Pleosporales*-*Penicillium*).

**Figure 6.**
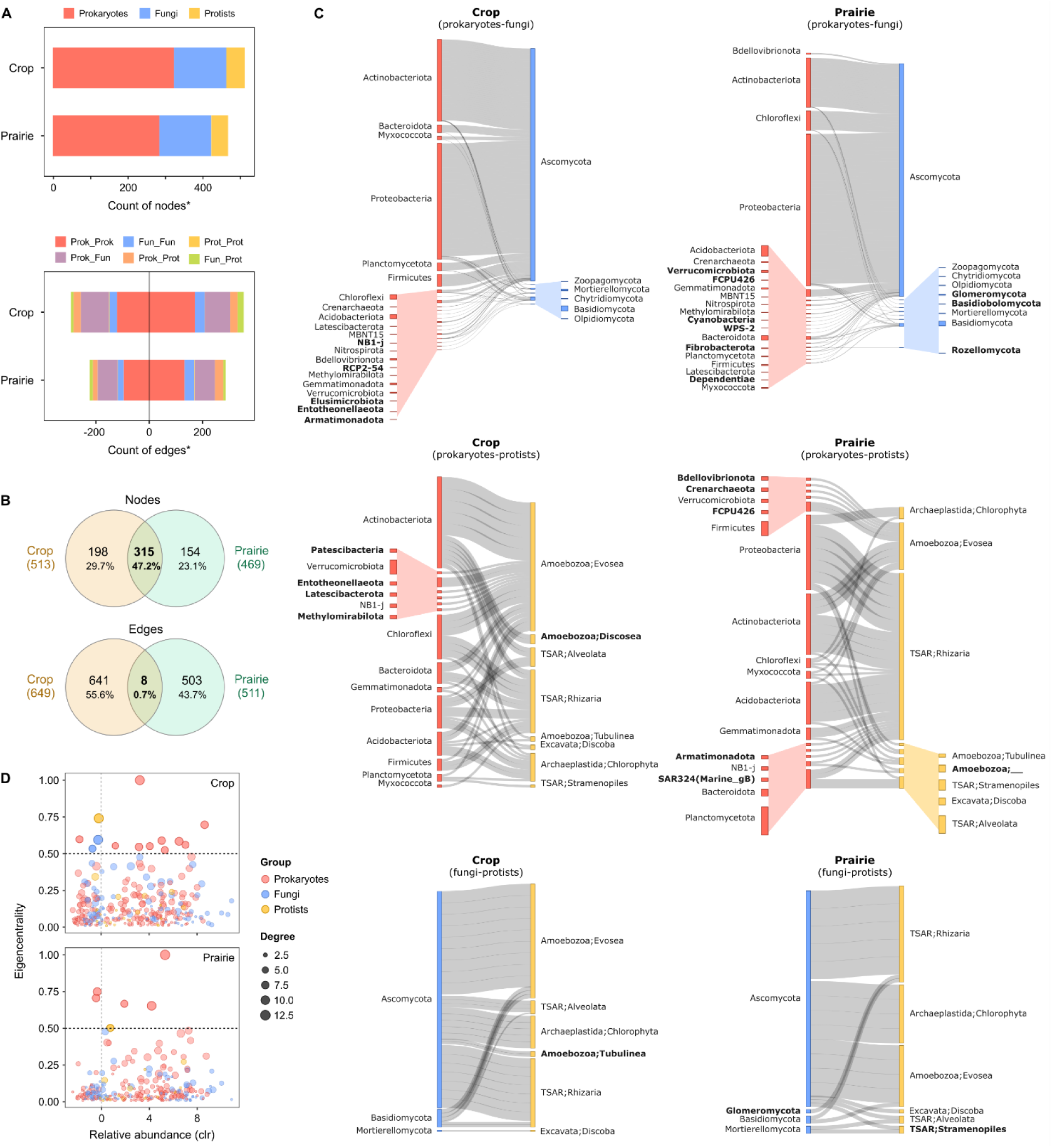
Overview of interkingdom network analysis of soil microbial genera from SPIEC-EASI comparing crop and prairie sites. A) Number of nodes and edges by microbial group, B) Nodes and edges in common between crop and prairie soils, C) Sankey plots summarizing interactions between taxa from different microbial groups according to their phyla or major protist group, D) Relationship between node relative abundance as centered log ratio (clr), degree (number of edges per node) and eigenvector centrality as a measure of keystone potential. In C, taxa names in bold show those present only in the network of that land use. In D, each point represents a node and those in darker shades (eigencentrality>0.5) were considered the most influential in each network. *Detailed information about influential nodes can be found in Table S8*.

Land use also determined differences in the major taxonomic groups involved in these interkingdom associations (Figure 6C). Prokaryote-fungus associations were mostly between Actinobacteriota or Proteobacteria and Ascomycota in both uses, although with a higher contribution of Actinobacteria in crop than restored prairie soils. Prokaryote-protist associations presented a more even contribution of prokaryotic taxa than other interactions, although Actinobacteriota, and in restored prairie soils Proteobacteria, were still quite frequent (Figure 6C). In these associations, the protist groups Evosea (Amoebozoa) and Rhizaria (TSAR) were the most frequent, with the latter being more important in restored prairie soils and vice versa. Notably, Chloroflexi was more in prokaryote-fungus links in restored prairie soils but prokaryote-protist links in crop soils. Fungus-protist associations also showed a dominance of Ascomycota and the protist groups Evosea and Rhizaria, with similarly important contribution of Chlorophyta in restored prairie soils. We also found several groups that were only involved in associations from only one land use (Figure 6C). In the restored prairie network, these included 6 prokaryotic phyla (*e.g.*, Verrucomicrobiota, Cyanobacteria, Fibrobacterota) and three fungal phyla (Glomeromycota, Basidiobolomycota and Rozellomycota) in prokaryote-fungus associations. Glomeromycota was also only present in fungi-protist associations of prairie but not crop soils. Five prokaryotic phyla (*e.g.*, Bdellovibrionota, Crenarchaeota and Armatimonadota) were also uniquely present in prairie prokaryote-protist associations.

Based on both degree and eigenvector centrality, which we used as a measure of keystone potential, crop soils had a higher number of influential nodes than prairie soils (24 vs. 12, respectively) (Figure 7D, Table S8). Most of these taxa were bacteria for both crop and prairie soils, and fungal keystone taxa were only present in crop soils (Figure 7D, Table S8). In the crop network, influential taxa were dominantly from Actinobacteria (*e.g.*, *Nonomuraea*, *Streptomyces, Geodermatophilus*) and Ascomycota (*e.g.*, *Aaosphaeria*, *Staphylotrichun*, *Pleotrichocladium*), but also included some members from other phyla (Proteobacteria, Acidobacteria, Planctomycetota, Chloroflexi, Verrucomicrobiota) (Table S8). Influential nodes in restored prairie soils include members from Proteobacteria (*e.g.*, Burkholderia), Chloroflexi (AD3), Gemmatimonadetes (AKAU4049) and Actinobacteria (*e.g.*, Rubrobacter). The nodes *Blastococcus* (Actinobacteria, Frankiales) and AD3 (Chloroflexi) were influential in both crop and prairie networks (Table S8).

**Figure 7.**
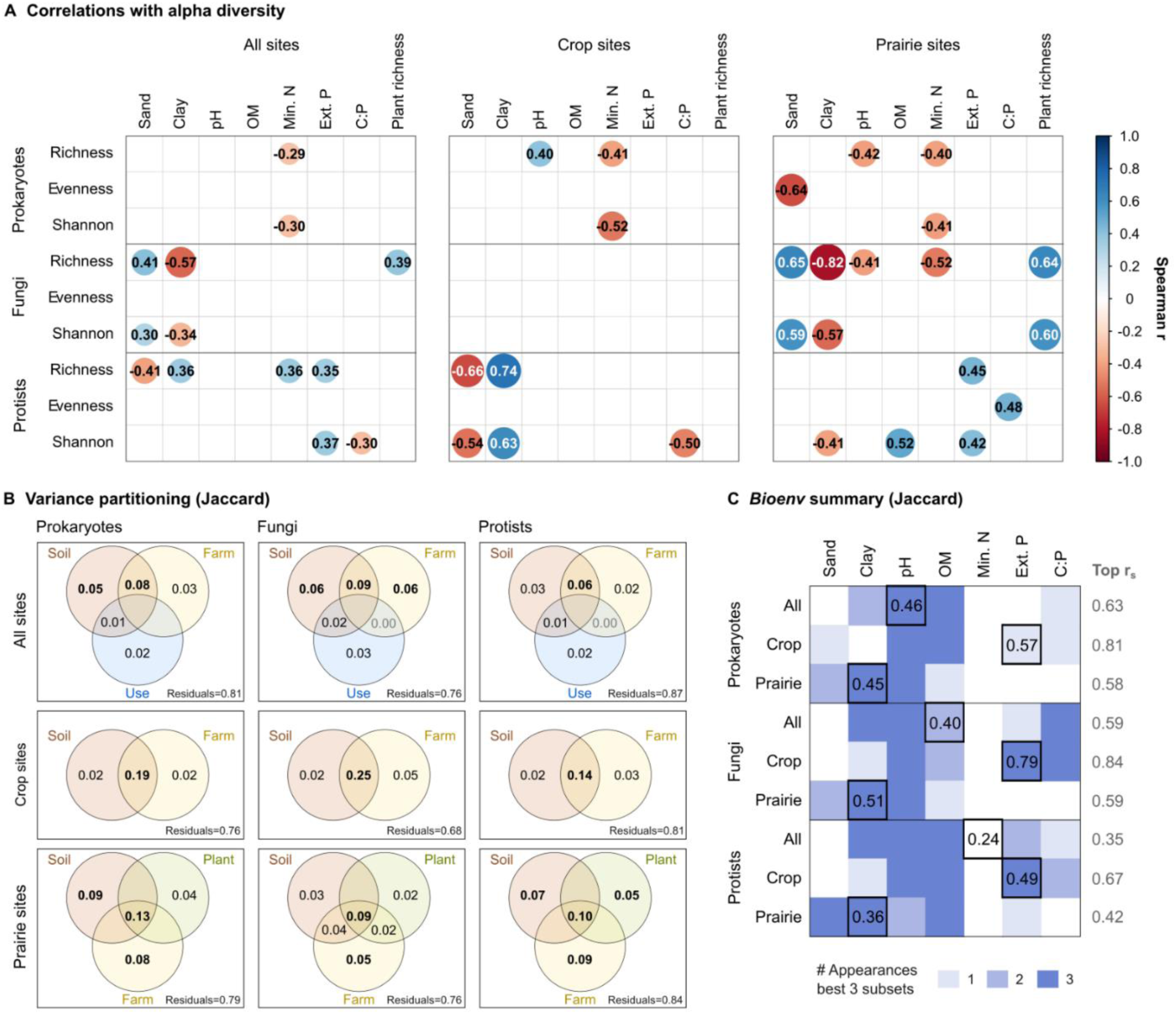
Relationship between soil microbial communities and soil environmental variables. A) Correlations between environmental variables and alpha diversity metrics in the complete dataset (all sites) and in each land use separately (crop sites and prairie sites). B) Variance partitioning analysis showing contributions (adjusted R-squared values) of land use, farm-specific effects, soil physicochemical variables and/or plant community composition to community composition, C) Summary of *bioenv* analysis showing soil physicochemical variables with the highest correlation with community dissimilarities (*i.e.*, appearing the subsets of variables with the highest correlations). In B, bold numbers highlight adjusted R-squared > 0.05. In C, variables highlighted with a thick border are those with the highest correlation on its own and the number represents the Spearman correlation value.

### 3.7- Relationship between soil microbiota and environmental variables

Correlations between alpha diversity metrics and environmental variables were limited, mostly low (|r_s_|<0.5), and more frequent in restored prairie soils (Figure 7A). Correlations suggested that fungal alpha diversity (richness and Shannon) was higher in coarser textures (*i.e.*, higher sand and lower clay content), especially in restored prairie sites. However, the opposite relationship was found for prokaryotic evenness in restored prairie sites and protist richness in crop sites. There was also a negative relationship between soil mineral N content and prokaryotic diversity (richness and Shannon), regardless of the land use. The same was observed for fungal richness but only in restored prairie soils. Soil OM and extractable P were only associated with protist diversity, with positive correlations in restored prairie soils. Plant richness, on the other hand, was only correlated with fungal diversity showing a positive association with richness and the Shannon index in restored prairie soils.

To relate environmental variables with microbial community composition (Jaccard distance), we first used variance partitioning, to compare the relative contribution of soil physicochemical properties, land use and other farm effects. For all microbial groups. Based on the adjusted R^2^, we found that the joint effect of soil properties and farm effects explained the highest proportion of the variation (Figure 7B). For prokaryotes and fungi, this was followed by the fraction explained uniquely by soil properties. And, in the case of fungi, the fraction explained by farm-specific factors also equally influential. We also carried out a similar analysis on crop and restored prairie sites separately, which revealed mainly joint effects of soil properties and farm-specific factors on crop microbial communities (Figure 7B). Similarly, in restored prairie sites it was the joint effect of soil, farm and plant composition that explained the highest fraction of variation in all microbial groups, followed by farm effects and, except for fungi, soil properties (Figure 7B).

To inquire on relative importance of different soil physicochemical variables, we explored which subsets of variables have the highest correlation with community composition (Jaccard) using *bioenv* (Figure 7C). Soil pH was one of the most influential variables for all microbial groups, being present in almost all the best subsets. After pH, the most important variables were soil OM and texture for prokaryotes and protists, and texture and C:P for fungi. There were some differences when analyzing each land use separately, such as extractable P becoming more influential in crop soils and soil texture in restored prairie soils. When looking at the complete dataset, the individual soil variables with highest influence were pH for prokaryotes (r_s_=0.46), OM for fungi (r_s_=0.40) and mineral N for protists (r_s_=0.24). Still, the association with community composition was never higher than the best combination of soil variables: r_s_=0.63 for prokaryotes, r_s_=0.59 for fungi, and r_s_=0.35 for protists.

## 4- Discussion

This study allowed us to characterize soil microbial communities from productive farms of southwestern Ontario using a metabarcoding approach. Overall, 5-9 years of tallgrass prairie restoration induced a clear shift in the composition of soil prokaryotic, fungal and protist communities, including changes in specific taxa and functional groups, but did not have a positive effect on their alpha diversity or abundance. The fact that prairie restoration effects were not consistent between microbial groups suggests they respond differently to the environmental changes imposed by this practice, likely due to ecological and adaptive differences in environmental preference, as well as differences in dispersal ability (*e.g.*, Grossmann *et al*. 2016; Chaudhary *et al*. 2022; King *et al*. 2022). Yet, based on the correlations in terms of community composition and interkingdom associations, there is some degree of co-response and/or interkingdom ecological interactions, such as predation of bacteria or fungi by protists (Glücksman et al., 2010; Degrune et al., 2024). The variability observed across farms reveals a complex response of soil microbiota to prairie restoration, possibly resulting from multiple interacting factors (*e.g.*, soil properties, vegetation, current and historical management) rather than any single explanatory driver. This complex response, with site-to-site variation, is known to affect restoration outcomes and predictability (Brudvig and Catano, 2021)

Due to the observational nature of this study, we cannot identify the precise cause behind farm-to-farm differences, but we were able to identify some potentially relevant factors modulating the effect of prairie restoration on soil microbiota. Soil type and physicochemical properties seem to be a relevant modulating factor based on our results and published data (Jangid et al., 2010; Sha et al., 2023a). As mentioned for soil OC, coarser textured soils as studied here (3.2-15% clay) could slow down the recovery of soil microbiota by offering them less protection and limiting aggregate formation (Bach et al., 2010; Baer et al., 2010). Besides, there could be an influence of persistent legacies in soil properties from agricultural activity, such as relatively higher P levels and bulk density (Mazzorato et al., 2022). Given the vertical variation in soil properties, as well as the different habitats and life strategies of microbial groups, responses could also vary with soil depth (Hu et al., 2020). On the other hand, prairie vegetation was linked with microbial diversity and composition (Figure 7) but did not seem to modulate the response of soil microbiota to restoration, as opposed as was previously found for bacterial diversity (Xu et al., 2022; Shu et al., 2023) and other properties (Sha et al., 2023a). Prairie management practices (*e.g.*, fertilization, burning) are also expected to drive microbial response (Smith et al., 2003; Jangid et al., 2010; Bach and Hofmockel, 2015; Hu et al., 2020), but unfortunately we could not gather this information for the studied farms. Management history in restored sites may also be a critical factor in determining the success of restoration, as it establishes different starting points. Space-to-time substitutions like this study pose an additional challenge, as there could be differences in soil type and prior management between crop and restored prairie sites. For example, the crop site in F5 had previously been an orchard, which could explain the high soil OC and contrasting responses in soil microbial communities.

It is also possible that restoration time was not enough to evidence clear and consistent changes in soil microbial communities, based on previous findings in restoration or abandonment chronosequences (Lozano et al., 2014; Baer et al., 2015; Barber et al., 2017; Sha et al., 2023a). Even after several years, restored prairie communities might not fully resemble those from native prairies (Jangid et al., 2010; Mackelprang et al., 2018). Besides, spatial properties of the restored prairies (*i.e.*, size, shape, location) could be relevant to restoration success. Spatial properties can affect the influence of neighboring croplands but also the proximity to propagules or source populations to recolonize the restored land, something particularly relevant in areas cultivated for several years (Baer et al., 2012). In our study, we cannot properly separate the effects of restoration time and prairie size because the older prairies (8-9 years) were also the largest (2.4-7 ha) (*i.e.*, F2, F3 and F5), but these farms evidenced greater prairie-to-crop shifts in fungal community composition (Figure 2B, Table 1). By contrast, F4 farm, with the youngest and one of the smallest restored prairie sites (but also the only one with switchgrass), generally showed the smallest distances between crop and prairie sites (Figure 2B, Table 1).

### 4.1- Recovered diversity and taxa in this study

We recovered a higher number of prokaryotic ASVs and genera, followed by fungi and lastly protists, similar to previous reports in grassland (Dassen et al., 2017) and alpine soils (Malard et al., 2022). A study by Romdhane *et al*. (2022) recovered more protist than fungal OTUs (97% similarity); this could be explained by the different protist primers, OTU clustering, but mostly by their decision to keep non-protist groups (*e.g.*, Metazoa) (Mau et al., 2024). The overall taxonomic composition of soil microbial communities was also consistent with the literature, with prokaryotic communities dominated by Proteobacteria, Actinobacteria and Acidobacteria (Delgado-Baquerizo et al., 2018). Regarding fungi, a dominance of Ascomycota has also been found in grasslands in a global survey by Egidi *et al*. (2019), contradicting earlier findings likely limited by the soil types and biomes surveyed (Tedersoo et al., 2014). Protist communities from both land uses were dominated by Rhizaria (TSAR), Evosea (Amoebozoa) and Chlorophyta (Archaeplastida), in agreement with studies using metatranscriptomics (Geisen et al., 2015) and metabarcoding (Leff et al., 2018). A study across 180 soils did not find a dominance of Amoebozoa (Oliverio et al., 2020), but we believe this was due to a loss of amoebozoan data when merging paired-end reads (Vaulot et al., 2022) (see Supplementary Methods).

### 4.2- Prairie restoration did not increase microbial abundance or diversity

Despite increases in plant cover and diversity after prairie restoration (*e.g.*, Dolezal et al., 2022), we did not observe significant shifts in soil microbial abundance or diversity even after nine years since last being cultivated (Figure 1). The responses we did detect depended more on the microbial group or farm being analyzed, with none of our five farms showing a consistently beneficial or detrimental effect of restoration on all microbial groups. Prairie restoration has previously been shown to promote soil bacterial and/or fungal biomass (Allison et al., 2005; Bach et al., 2010; Jangid et al., 2010; Baer et al., 2015), but here we only detected an average decrease in soil fungal abundance and the F:B ratio. Soil bacterial abundance increased with restoration at only two farms, possibly due to them showing increases in soil pH associated with restoration (Table 2). Our findings partially agree with those in a grassland restoration chronosequence, where phospholipid fatty acids (PLFA) fungal biomass decreased but bacterial biomass increased over time under restoration (Yang et al., 2023). These results are unexpected, considering soils usually become more dominated by fungi in later stages of succession, as a response to low nutrient inputs and higher root biomass with increased lignin and C:N ratio (Baer et al., 2002; Matamala et al., 2008; Maharning et al., 2009; Mazzorato et al., 2022). Indeed, we found restored grassland to have 20x more root biomass than adjacent crop field to depths of 60 cm (Noble et al., 2023). Other factors could be involved in our unexpected responses to restoration, especially relating to the differences in management among farms. For example, in the only farm (F2) showing a significantly higher fungal abundance in crop sites, there was also an increase in the frequency of the fungal genus *Cladorrhinum* (23.8% vs. 0.1-1.8% in all other sites). This could be explained by the sorghum cropping and fallow stage of this site, as *Cladorrhinum* can have different lifestyles (saprotroph, plant pathogen, plant mutualist) and was previously detected in sorghum cultivated soils (Carmarán et al., 2015).

Overall, microbial community richness was more sensitive to restoration than evenness, and fungi and protists were more sensitive than prokaryotes. However, alpha diversity metrics did not respond in unison to restoration, with positive, negative and neutral effects by farm and by microbial group. In other words, there was no particular microbial group responding consistently across sites, but also not a specific site where prairie restoration was consistently positive or negative for all groups. Such variability in how soil microbial richness and diversity responds to restoration or reduced land use intensity is consistent with other authors (Sha et al., 2023b), with decreases for bacteria (Barber et al., 2017, 2023; Mackelprang et al., 2018), increases for bacteria but not fungi (Y. Guo et al., 2021), or no clear changes (Labouyrie et al., 2023). Based on a brief exploration (Figure S3), it is possible that microbial diversity exhibits a phylum-specific response that is worth exploring further.

Despite the variability in response, there was some evidence that soil texture could be mediating farm-specific responses to restoration. For example, fungal richness, and to a lesser extent protist richness, only responded positively on farms with coarser textured soils (Figure 1, Table 1). In a grassland restoration chronosequence, Bach *et al*. (2010) found clearer responses of microbial PLFA profiles in finer textures but exploring soils with much higher silt and clay contents than ours (*i.e.*, 71-90%). It is worth considering that farm F5 could be disproportionately driving this trend, as the decrease in fungal richness was associated with the high soil OM and low extractable P of its crop site (Table 2). Prokaryotic evenness (genus level) also changed only in coarser textured soils, but with negative effects of prairie restoration (Figure S5), linked to a higher dominance of three Acidobacteriota (Pyrinomonadaceae_RB41, unidentified Vicinamibacteraceae and Vicinamibacterales) and the N_2_-fixing genus *Bradyrhizobium* (Proteobacteria).

### 4.3- Prairie restoration modified microbial community composition and functional groups

All microbial groups exhibited a clear shift in community composition in response to restoration, both in terms of presence-absence and relative abundance of ASVs and genera. Based on the higher turnover, lower nestedness and less shared and generalist features, we can conclude that changes in composition were mainly caused by ASVs in restored prairies replacing those previously present in crop soils (Figure 3, Table S5). As expected, this was less clear looking at microbial genera, where we found that fungi had more specialized taxa than prokaryotes and protists, consistently with findings by Malard *et al*. (2022) in alpine soils. Differences in soil microbial community composition have been previously detected between crop and native tallgrass prairie (Cornell et al., 2022, 2023), and between crops and restored prairies or grasslands (Allison et al., 2005; Jangid et al., 2010; Herzberger et al., 2014; Hu et al., 2020; Armbruster et al., 2021), with many of the latter using the lower resolution PLFA analysis. Still, restored prairie communities can sometimes be more similar to crop than native prairie communities (Mackelprang et al., 2018; Armbruster et al., 2021), which could explain the relative small proportion of community composition associated with restoration (Figure 7, Table 3). In this sense, further shifts could be expected with longer restoration time, as it may weaken legacy effects on soil properties and biota (Lozano et al., 2014; Cline and Zak, 2015; Barber et al., 2017; Yang et al., 2023). Comparing microbial groups, prokaryotes showed the strongest associations with other groups, while the weakest were between fungi and protists, similar to previous findings (Malard et al., 2022). Despite these similarities between groups, and the consistent compositional shifts across farms, we again observed group-specific and farm-specific effects in the magnitude and direction of these changes (Figure 2, Table 3).

Our results also demonstrate consistent restoration-associated changes in the frequency and relative abundance of some microbial taxa. Based on the differential abundance analyses, the bacterial phyla Latescibacterota, Desulfobacterota and Acidobacteriota increased with prairie restoration. Acidobacteriota is a widespread, abundant and largely unexplored phylum which encompasses some known heterotrophic organisms with a diverse carbohydrate utilization capacity and ecophysiology (Kielak et al., 2016; Kalam et al., 2020). This phylum has been previously associated with tallgrass prairie restoration (Barber et al., 2017), although it could decrease over time under grassland restoration (Yang et al., 2023). Latescibacterota and Desulfobacterota are commonly found in low abundance in soils and, notably, both groups happen to harbor bacteria with anaerobic metabolism: anaerobic fermentation in Latescibacterota (Youssef et al., 2015) and Fe(III)-reduction in Geobacteraceae (Megonigal et al., 2003), the most represented family in our study. Crop soils, on the other hand, had a higher relative abundance of Deinococcota (mostly Deinococcaceae), which were more abundant in biocrusts (Wang et al., 2024) and are characterized by their high resistance to UV radiation (Seshadri et al., 2023), consistent with the lower vegetation cover and higher exposure of crop soils to direct sunlight.

Prairie restoration also increased the relative abundance of the fungal phylum Glomeromycota [arbuscular mycorrhizal (AM) fungi] (Figure 4), in agreement with AM fungal spore density measurements in these sites (MacColl et al., 2024) and PLFA abundance in other tallgrass prairie restoration studies (Allison et al., 2005; Baer et al., 2015). As found here, soil AM fungi were found to be more responsive than other microbial groups in early stages of restoration (*i.e.*, 2-4 years) (Herzberger et al., 2014). This early response could stimulate vegetation development and ecosystem recovery (Mao et al., 2019) and boost carbon sequestration in soils (Frey, 2019). The increase in AM fungi in restored grassland compared to croplands could be explained by the elimination of tillage, higher plant cover and root biomass, and lower mineral N and P content from fertilization, which could increase plant carbon allocation to the fungal symbiont (Kabir, 2005; Wang et al., 2009; Barceló et al., 2020). Whereas in this study Glomeromycota were promoted evenly, a more detailed analyses of AM fungal communities on these soils showed family-level differences in relative abundance within this group (MacColl et al., 2024). Prairie restoration also favoured two other fungal genera with plant-growth promoting traits (*Serendipita* and *Mortierella*), consistently with previous findings (Verbruggen et al., 2014; Armbruster et al., 2021) (Table S7). The fact that prairie restoration favoured more symbiotrophs and less plant pathogens and saprotrophs (Table 3) is in agreement with previous studies on land perturbation (Labouyrie et al., 2023) and N and P fertilization (Lekberg et al., 2021). While the effect of disturbance may be more direct, Lekberg *et al*. (2021) suggest that nutrient content affects fungal guilds indirectly, via changes in plant communities. With prairie restoration we also observed a reduction in the relative abundance of chytrids (Chytridiomycota), which are zoosporic fungi with saprophytic and parasitic lifestyles (Hanrahan-Tan et al., 2023a). The environmental adaptation and stress tolerance found within this phylum (Gleason et al., 2004; Freeman et al., 2009; Hanrahan-Tan et al., 2023b) could explain their thriving in agricultural relative to restored prairie soils.

Soil protists are relatively understudied compared to prokaryotes and fungi, but there is evidence of their response to anthropogenic disturbances, mostly from specific taxa and/or using traditional quantification or identification approaches (Geisen et al., 2017, 2018). Phagotrophic protists were the dominant trophic group in all soils, consistent with previous reports (Oliverio et al., 2020; Santos et al., 2020), and the only group affected by land use (Figure S7, Table 3, S7). This suggests prairie restoration can affect higher trophic levels in the soil food web and may alter nutrient cycling via changes in microbial predation and nutrient release dynamics (*i.e.*, ‘microbial loop’) (Geisen et al., 2018). Crop systems favoured more protist taxa than restored prairies, in agreement with increases in this trophic group with increasing land use intensity (Schulz et al., 2019). Moreover, Santos *et al*. (2020) found lower phagotroph richness in grassland compared to crop soils. Yet, our findings conflict with reports of negative effects of mineral fertilization on protists, in particular consumers (Zhao et al., 2019, 2020; S. Guo et al., 2021). The conventional tillage practices applied in these crop systems may also shift protist community composition (Ma et al., 2024) by modifying soil structure, porosity and water dynamics, which can in turn affect protist taxa (Geisen et al., 2014). It is also possible that the adaptation of certain protists to crop or prairie conditions is associated with their adaptive strategies (*e.g.*, resistance structures) and specific feeding habits (Geisen et al., 2018). With the available data, we can only affirm that the protist taxa promoted in crops are mainly bacteriovores, although many can also consume other microorganisms, protists and small organic matter particles. By contrast, prairie restoration promoted two taxa in the order Vampyrellida (Table S8), which have been considered for their potential biocontrol by consuming other eukaryotes (Chandarana and Amaresan, 2022; Hess and Suthaus, 2022) and were previously negatively affected by fertilization (Zhao et al., 2019). These feeding habits are not consistent with our results in terms of bacterial and fungal abundance, but these complex responses may not be well represented by a single snapshot of soil communities, plus each taxon can have multiple types of prey and mycophagy may be more common than expected (Geisen et al., 2016; Adl et al., 2018).

### 4.4- Prairie restoration affected intra– and interkingdom microbial associations

To inquire further into the associations between microbial taxa, we carried out interkingdom network analyses. Even though network analyses have limitations to infer actual microbe-microbe interactions, especially at a coarse spatial scale (Berry and Widder, 2014; Carr et al., 2019; Goberna and Verdú, 2022), they are useful to infer co-occurrence and association patterns that could represent such interactions or common responses to biotic/abiotic factors. Notably, despite sharing almost 50% of nodes (taxa), crop and restored prairie networks only had 8 edges (associations) in common, revealing drastic changes in microbial association patterns. Unlike earlier studies (Morriën et al., 2017; Guo et al., 2019), the crop soils network also presented a higher number of taxa and associations than the restored prairie networks. Similar findings have been reported when evaluating land use intensity (Romdhane et al., 2022) and prairie-to-crop conversion (Cornell et al., 2023). Cornell *et al*. (2023) attributed the higher complexity of crop networks to lower diversity and biotic/abiotic homogenization (*i.e.*, increased niche sharing), none of which were clearly observed in our data. The most connected were mostly bacterial, in contrast to Romdhane *et al*. (2022), who found protists were the most connected group but also had the highest number of nodes. These potentially keystone taxa differed between land uses with one exception (Chloroflexi AD3), suggesting a role of land use intensity on which taxa are the most influential for the community (Banerjee et al., 2019).

### 4.5- Potential environmental drivers of microbial response to prairie restoration

The influence of soil physicochemical properties and plant species composition on microbial diversity and composition was both linked and independent from prairie restoration and farm-to-farm variability (Figure 7). Effects of soil properties and vegetation type on soil microbiota have been previously reported (Trivedi et al., 2016; Guo et al., 2019; Malard et al., 2022), sometimes surpassing those of land use (Kuramae et al., 2012). The influence of soil texture, organic matter and nutrient content on microbial communities, and pH specifically on prokaryotic communities, is consistent with the literature (Fierer, 2017). The fraction of community composition not explained by any environmental, land use or farm-specific factors was larger than previously reported, possibly due to the variability associated with observational studies. Yet, this could become less important over time under restoration (Guo et al., 2019). Notably, fungal communities seemed to be the most susceptible to the evaluated factors (*i.e.*, lower residual variability), consistently with the highest proportion of specialists. Fungi were also the only group with a positive response to plant species diversity in restored prairie sited, similar to an experimental study by Shen *et al*. (2021), who also found Basidiomycota to be responsive in terms of richness (Figure S3).

Most of the measured soil physicochemical properties did not clearly respond to prairie restoration. An exception was mineral nitrogen, which decreased in agreement with published studies (Baer et al., 2002; Mazzorato et al., 2022), likely due to lack of mineral fertilization. Our results in soil OC are consistent with those of Mazzorato *et al*. (2022) in these and other farms in the area, where overall increases with restoration were only small (+6%) and insignificant. These findings contradict reports of positive responses to grassland or prairie restoration (Baer et al., 2002, 2015; Rosenzweig et al., 2016; Li et al., 2021; Sha et al., 2023a). In some cases, carbon sequestration may take longer than the 5-9 years of restoration of these prairies (Matamala et al., 2008; De et al., 2020), although earlier responses have been observed in upper soil layers (*e.g.*, 0-5 cm) (Baer et al., 2002; Matamala et al., 2008; Li et al., 2019). This process is also modulated by factors such as management, climate, topography and soil type, which may surpass the importance of time since restoration (Matamala et al., 2008; De et al., 2020; Mazzorato et al., 2022; Kimmell et al., 2023), consistent with among-farm differences. In particular, soil texture may play a key role, as higher clay content provides more physicochemical protection and, thus, higher restorative potential for soil OC (Baer et al., 2010).

It is also worth noting that soil physicochemical properties were more variable across crop than prairie soils (Table 2), which could explain why crop sites presented higher beta dispersion of prokaryotic communities (Table 3), clearer changes in the composition of major protist groups (Figure S6) and higher variability in bacterial and fungal abundance (Figure 1A-C). Based on our results, the assumption that agriculture tends to ‘homogenize’ soil microbial communities (Montecchia et al., 2015; Peng et al., 2024) might depend on the studied sites.

## 5- Conclusions

Given the importance of soil microbes in ecosystem functioning, it is crucial to understand their role and potential for manipulation in restoration practices (Coban et al., 2022). During restoration, belowground communities may respond earlier than aboveground communities (van der Bij et al., 2017; Graham and Knelman, 2023), with beneficial microbes potentially facilitating the establishment of the re-introduced plant species (Farrell et al., 2020). However, our study reveals a highly complex response that opens doors to several questions to be answered before establishing restoration goals and developing microbiome manipulations (*e.g.*, inoculations). Future research efforts should aim at disentangling the complex interactions behind the response of soil biota to ecosystem restoration, for example by better understanding the drivers of microbial assembly, along with the environmental and management factors discussed above. More research is also needed to explore temporal/seasonal dynamics (*e.g.*, Jangid *et al*., 2010; Barber *et al*., 2017), different soil depths (*e.g.*, Hu *et al*., 2020) and the response of active fractions of the microbial community (*e.g.*, using RNA sequencing). This will also better characterize the relationship between the microbial dynamics and ecosystem services as restoration progresses. Finally, we encourage researchers to investigate spillover effects of restored prairie strips on neighboring crop sites (Kemmerling et al., 2022; Dutter et al., 2024), as well as benefits at the farm-scale.

## 7- Acknowledgements

We would like to thank Kamini Khosla and all the Kari Dunfield Lab and Hafiz Maherali Lab members that helped with sample collection and processing. Special thanks to all the participating farmers from the Alternative Land Use Services (ALUS) program.

## References

1. Adl, M.S., Gupta, V.V.S.R., 2006. Protists in soil ecology and forest nutrient cycling. Canadian Journal of Forest Research 36, 1805–1817. doi:10.1139/X06-056

2. Adl, S.M., Bass, D., Lane, C.E., Lukeš, J., Schoch, C.L., Smirnov, A., Agatha, S., Berney, C., Brown, M.W., Burki, F., Cárdenas, P., Čepička, I., Chistyakova, L., del Campo, J., Dunthorn, M., Edvardsen, B., Eglit, Y., Guillou, L., Hampl, V., Heiss, A.A., Hoppenrath, M., James, T.Y., Karnkowska, A., Karpov, S., Kim, E., Kolisko, M., Kudryavtsev, A., Lahr, D.J.G., Lara, E., Le Gall, L., Lynn, D.H., Mann, D.G., Massana, R., Mitchell, E.A.D., Morrow, C., Park, J.S., Pawlowski, J.W., Powell, M.J., Richter, D.J., Rueckert, S., Shadwick, L., Shimano, S., Spiegel, F.W., Torruella, G., Youssef, N., Zlatogursky, V., Zhang, Q., 2018. Revisions to the Classification, Nomenclature, and Diversity of Eukaryotes. Journal of Eukaryotic Microbiology 66, 4–119. doi:10.1111/jeu.12691

3. Allison, V.J., Miller, R.M., Jastrow, J.D., Matamala, R., Zak, D.R., 2005. Changes in Soil Microbial Community Structure in a Tallgrass Prairie Chronosequence. Soil Science Society of America Journal 69, 1412–1421. doi:10.2136/sssaj2004.0252

4. Anthony, M.A., Bender, S.F., van der Heijden, M.G.A., 2023. Enumerating soil biodiversity. Proceedings of the National Academy of Sciences 120, 2017. doi:10.1073/pnas.2304663120

5. Armbruster, M., Goodall, T., Hirsch, P.R., Ostle, N., Puissant, J., Fagan, K.C., Pywell, R.F., Griffiths, R.I., 2021. Bacterial and archaeal taxa are reliable indicators of soil restoration across distributed calcareous grasslands. European Journal of Soil Science 72, 2430–2444. doi:10.1111/ejss.12977

6. Bach, E.M., Baer, S.G., Meyer, C.K., Six, J., 2010. Soil texture affects soil microbial and structural recovery during grassland restoration. Soil Biology and Biochemistry 42, 2182– 2191. doi:10.1016/j.soilbio.2010.08.014

7. Bach, E.M., Hofmockel, K.S., 2015. Coupled Carbon and Nitrogen Inputs Increase Microbial Biomass and Activity in Prairie Bioenergy Systems. Ecosystems 18, 417–427. doi:10.1007/s10021-014-9835-8

8. Baer, S.G., Bach, E.M., Meyer, C.K., Du Preez, C.C., Six, J., 2015. Belowground Ecosystem Recovery During Grassland Restoration: South African Highveld Compared to US Tallgrass Prairie. Ecosystems 18, 390–403. doi:10.1007/s10021-014-9833-x

9. Baer, S.G., Heneghan, L., Eviner, V.T., 2012. Applying soil ecological knowledge to restore ecosystem services, in: Wall, D.H. et al. (Ed.), Soil Ecology and Ecosystem Services. OUP, Oxford, pp. 377–393.

10. Baer, S.G., Kitchen, D.J., Blair, J.M., Rice, C.W., 2002. Changes in ecosystem structure and function along a chronosequence of restored grasslands. Ecological Applications 12, 1688–1701. doi:10.1890/1051-0761(2002)012[1688:CIESAF]2.0.CO;2

11. Baer, S.G., Meyer, C.K., Bach, E.M., Klopf, R.P., Six, J., 2010. Contrasting ecosystem recovery on two soil textures: Implications for carbon mitigation and grassland conservation. Ecosphere 1, 1–22. doi:10.1890/ES10-00004.1

12. Banerjee, S., Walder, F., Büchi, L., Meyer, M., Held, A.Y., Gattinger, A., Keller, T., Charles, R., van der Heijden, M.G.A., 2019. Agricultural intensification reduces microbial network complexity and the abundance of keystone taxa in roots. ISME Journal 13, 1722–1736. doi:10.1038/s41396-019-0383-2

13. Barber, N.A., Chantos-Davidson, K.M., Amel Peralta, R., Sherwood, J.P., Swingley, W.D., 2017. Soil microbial community composition in tallgrass prairie restorations converge with remnants across a 27-year chronosequence. Environmental Microbiology 19, 3118–3131. doi:10.1111/1462-2920.13785

14. Barber, N.A., Klimek, D.M., Bell, J.K., Swingley, W.D., 2023. Restoration age and reintroduced bison may shape soil bacterial communities in restored tallgrass prairies. FEMS Microbiology Ecology 99, 1–12. doi:10.1093/femsec/fiad007

15. Barceló, M., van Bodegom, P.M., Tedersoo, L., den Haan, N., Veen, G.F. (Ciska), Ostonen, I., Trimbos, K., Soudzilovskaia, N.A., 2020. The abundance of arbuscular mycorrhiza in soils is linked to the total length of roots colonized at ecosystem level. PLOS ONE 15, e0237256. doi:10.1371/journal.pone.0237256

16. Baselga, A., 2012. The relationship between species replacement, dissimilarity derived from nestedness, and nestedness. Global Ecology and Biogeography 21, 1223–1232. doi:10.1111/j.1466-8238.2011.00756.x

17. Berry, D., Widder, S., 2014. Deciphering microbial interactions and detecting keystone species with co-occurrence networks. Frontiers in Microbiology 5, 1–14. doi:10.3389/fmicb.2014.00219

18. Bokulich, N.A., Dillon, M., Bolyen, E., Kaehler, B., Huttley, G., Caporaso, J., 2018. Q2-Sample-Classifier: Machine-Learning Tools for Microbiome Classification and Regression. Journal of Open Source Software 3, 934. doi:10.21105/joss.00934

19. Bolyen, E., Rideout, J.R., Dillon, M.R., Bokulich, N.A., Abnet, C.C., Al-Ghalith, G.A., Alexander, H., Alm, E.J., Arumugam, M., Asnicar, F., Bai, Y., Bisanz, J.E., Bittinger, K., Brejnrod, A., Brislawn, C.J., Brown, C.T., Callahan, B.J., Caraballo-Rodríguez, A.M., Chase, J., Cope, E.K., Da Silva, R., Diener, C., Dorrestein, P.C., Douglas, G.M., Durall, D.M., Duvallet, C., Edwardson, C.F., Ernst, M., Estaki, M., Fouquier, J., Gauglitz, J.M., Gibbons, S.M., Gibson, D.L., Gonzalez, A., Gorlick, K., Guo, J., Hillmann, B., Holmes, S., Holste, H., Huttenhower, C., Huttley, G.A., Janssen, S., Jarmusch, A.K., Jiang, L., Kaehler, B.D., Kang, K. Bin, Keefe, C.R., Keim, P., Kelley, S.T., Knights, D., Koester, I., Kosciolek, T., Kreps, J., Langille, M.G.I., Lee, J., Ley, R., Liu, Y.-X., Loftfield, E., Lozupone, C., Maher, M., Marotz, C., Martin, B.D., McDonald, D., McIver, L.J., Melnik, A. V., Metcalf, J.L., Morgan, S.C., Morton, J.T., Naimey, A.T., Navas-Molina, J.A., Nothias, L.F., Orchanian, S.B., Pearson, T., Peoples, S.L., Petras, D., Preuss, M.L., Pruesse, E., Rasmussen, L.B., Rivers, A., Robeson, M.S., Rosenthal, P., Segata, N., Shaffer, M., Shiffer, A., Sinha, R., Song, S.J., Spear, J.R., Swafford, A.D., Thompson, L.R., Torres, P.J., Trinh, P., Tripathi, A., Turnbaugh, P.J., Ul-Hasan, S., van der Hooft, J.J.J., Vargas, F., Vázquez-Baeza, Y., Vogtmann, E., von Hippel, M., Walters, W., Wan, Y., Wang, M., Warren, J., Weber, K.C., Williamson, C.H.D., Willis, A.D., Xu, Z.Z., Zaneveld, J.R., Zhang, Y., Zhu, Q., Knight, R., Caporaso, J.G., 2019. Reproducible, interactive, scalable and extensible microbiome data science using QIIME 2. Nature Biotechnology 37, 852–857. doi:10.1038/s41587-019-0209-9

20. Brudvig, L.A., Catano, C.P., 2021. Prediction and uncertainty in restoration science. Restoration Ecology 1–6. doi:10.1111/rec.13380

21. Burki, F., Sandin, M.M., Jamy, M., 2021. Diversity and ecology of protists revealed by metabarcoding. Current Biology 31, R1267–R1280. doi:10.1016/j.cub.2021.07.066

22. Callaham, M.A., Rhoades, C.C., Heneghan, L., 2008. A striking profile: Soil ecological knowledge in restoration management and science. Restoration Ecology 16, 604–607. doi:10.1111/j.1526-100X.2008.00490.x

23. Carmarán, C.C., Berretta, M., Martínez, S., Barrera, V., Munaut, F., Gasoni, L., 2015. Species diversity of Cladorrhinum in Argentina and description of a new species, Cladorrhinum australe. Mycological Progress 14, 94. doi:10.1007/s11557-015-1106-3

24. Carr, A., Diener, C., Baliga, N.S., Gibbons, S.M., 2019. Use and abuse of correlation analyses in microbial ecology. The ISME Journal 2647–2655. doi:10.1038/s41396-019-0459-z

25. Chandarana, K.A., Amaresan, N., 2022. Soil protists: An untapped microbial resource of agriculture and environmental importance. Pedosphere 32, 184–197. doi:10.1016/S1002-0160(21)60066-8

26. Chandrasoma, J.M., Udawatta, R.P., Anderson, S.H., Thompson, A.L., Abney, M.A., 2016. Soil hydraulic properties as influenced by prairie restoration. Geoderma 283, 48–56. doi:10.1016/j.geoderma.2016.08.001

27. Chaudhary, V.B., Aguilar-Trigueros, C.A., Mansour, I., Rillig, M.C., 2022. Fungal Dispersal Across Spatial Scales. Annual Review of Ecology, Evolution, and Systematics 53, 69–85. doi:10.1146/annurev-ecolsys-012622-021604

28. Cline, L.C., Zak, D.R., 2015. Soil microbial communities are shaped by plant-driven changes in resource availability during secondary succession. Ecology 96, 3374–3385. doi:10.1890/15-0184.1

29. Coban, O., de Deyn, G.B., van der Ploeg, M., 2022. Soil microbiota as game-changers in restoration of degraded lands. Science 375. doi:10.1126/science.abe0725

30. Cornell, C.R., Zhang, Y., Ning, D., Wu, L., Wagle, P., Steiner, J.L., Xiao, X., Zhou, J., 2022. Temporal Dynamics of Bacterial Communities along a Gradient of Disturbance in a U.S. Southern Plains Agroecosystem. MBio 13. doi:10.1128/mbio.03829-21

31. Cornell, C.R., Zhang, Y., Ning, D., Xiao, N., Wagle, P., Xiao, X., Zhou, J., 2023. Land use conversion increases network complexity and stability of soil microbial communities in a temperate grassland. ISME Journal 17, 2210–2220. doi:10.1038/s41396-023-01521-x

32. Cotterill, F.P.D., Al-Rasheid, K., Foissner, W., 2008. Conservation of protists: Is it needed at all? Biodiversity and Conservation 17, 427–443. doi:10.1007/s10531-007-9261-8

33. Dacey, D.P., Chain, F.J.J., 2021. Concatenation of paired-end reads improves taxonomic classification of amplicons for profiling microbial communities. BMC Bioinformatics 22, 493. doi:10.1186/s12859-021-04410-2

34. Dassen, S., Cortois, R., Martens, H., de Hollander, M., Kowalchuk, G.A., van der Putten, W.H., De Deyn, G.B., 2017. Differential responses of soil bacteria, fungi, archaea and protists to plant species richness and plant functional group identity. Molecular Ecology 26, 4085– 4098. doi:10.1111/mec.14175

35. De, M., Riopel, J.A., Cihacek, L.J., Lawrinenko, M., Baldwin-Kordick, R., Hall, S.J., McDaniel, M.D., 2020. Soil health recovery after grassland reestablishment on cropland: The effects of time and topographic position. Soil Science Society of America Journal 84, 568–586. doi:10.1002/saj2.20007

36. Degrune, F., Dumack, K., Ryo, M., Garland, G., Romdhane, S., Saghaï, A., Banerjee, S., Edlinger, A., Herzog, C., Pescador, D.S., García-Palacios, P., Fiore-Donno, A.M., Bonkowski, M., Hallin, S., van der Heijden, M.G.A., Maestre, F.T., Philippot, L., Glemnitz, M., Sieling, K., Rillig, M.C., 2024. The impact of fungi on soil protist communities in European cereal croplands. Environmental Microbiology 26, 1–9. doi:10.1111/1462-2920.16673

37. Delgado-Baquerizo, M., Oliverio, A.M., Brewer, T.E., Benavent-González, A., Eldridge, D.J., Bardgett, R.D., Maestre, F.T., Singh, B.K., Fierer, N., 2018. A global atlas of the dominant bacteria found in soil. Science 359, 320–325. doi:10.1126/science.aap9516

38. Dolezal, A.J., Esch, E.H., MacDougall, A.S., 2022. Restored marginal farmland benefits arthropod diversity at multiple scales. Restoration Ecology 30, 1–11. doi:10.1111/rec.13485

39. Dudley, N., Alexander, S., 2017. Agriculture and biodiversity: a review. Biodiversity 18, 45–49. doi:10.1080/14888386.2017.1351892

40. Dutter, C.R., Rutkoski, C.E., Evans, S.E., McDaniel, M.D., 2024. Contour prairie strips alter microbial communities and functioning both below and in adjacent cropland soils. Applied Soil Ecology 199, 105424. doi:10.1016/j.apsoil.2024.105424

41. Egidi, E., Delgado-Baquerizo, M., Plett, J.M., Wang, J., Eldridge, D.J., Bardgett, R.D., Maestre, F.T., Singh, B.K., 2019. A few Ascomycota taxa dominate soil fungal communities worldwide. Nature Communications 10, 2369. doi:10.1038/s41467-019-10373-z

42. FAO, 2022. The state of the world’s land and water resources for food and agriculture 2021 – Systems at breaking point. Main report. FAO, Rome. doi:10.4060/cb9910en

43. Farrell, H.L., Léger, A., Breed, M.F., Gornish, E.S., 2020. Restoration, soil organisms, and soil processes: emerging approaches. Restoration Ecology 28, S307–S310. doi:10.1111/rec.13237

44. Fernandes, A.D., Macklaim, J.M., Linn, T.G., Reid, G., Gloor, G.B., 2013. ANOVA-Like Differential Expression (ALDEx) Analysis for Mixed Population RNA-Seq. PLoS ONE 8, e67019. doi:10.1371/journal.pone.0067019

45. Fierer, N., 2017. Embracing the unknown: disentangling the complexities of the soil microbiome. Nature Reviews Microbiology 15, 579–590. doi:10.1038/nrmicro.2017.87

46. Fierer, N., Jackson, J., 2005. Assessment of soil microbial community structure by use of taxon-specific quantitative PCR assays. Applied and Environmental Microbiology 71, 4117. doi:10.1128/AEM.71.7.4117

47. Foley, J.A., DeFries, R., Asner, G., Barford, C., Bonan, G., Carpenter, S., Chapin, F., Coe, M., DAily, G., Gibbs, H., Helkowski, J., Holloway, T., Howard, E., Kucharik, C., Monfreda, C., Patz, J., Prentice, I., Ramankutty, N., Snyder, P., 2005. Global Consequences of Land Use. Science 309, 570–574. doi:10.1126/science.1111772

48. Freeman, K.R., Martin, A.P., Karki, D., Lynch, R.C., Mitter, M.S., Meyer, A.F., Longcore, J.E., Simmons, D.R., Schmidt, S.K., 2009. Evidence that chytrids dominate fungal communities in high-elevation soils. Proceedings of the National Academy of Sciences 106, 18315– 18320. doi:10.1073/pnas.0907303106

49. Frey, S.D., 2019. Mycorrhizal Fungi as Mediators of Soil Organic Matter Dynamics. Annual Review of Ecology, Evolution, and Systematics 50, 237–259. doi:10.1146/annurev-ecolsys-110617-062331

50. Geisen, S., Bandow, C., Römbke, J., Bonkowski, M., 2014. Soil water availability strongly alters the community composition of soil protists. Pedobiologia 57, 205–213. doi:10.1016/j.pedobi.2014.10.001

51. Geisen, S., Koller, R., Hünninghaus, M., Dumack, K., Urich, T., Bonkowski, M., 2016. The soil food web revisited: Diverse and widespread mycophagous soil protists. Soil Biology and Biochemistry 94, 10–18. doi:10.1016/j.soilbio.2015.11.010

52. Geisen, S., Mitchell, E.A.D., Adl, S., Bonkowski, M., Dunthorn, M., Ekelund, F., Fernández, L.D., Jousset, A., Krashevska, V., Singer, D., Spiegel, F.W., Walochnik, J., Lara, E., 2018. Soil protists: A fertile frontier in soil biology research. FEMS Microbiology Reviews 42, 293–323. doi:10.1093/femsre/fuy006

53. Geisen, S., Mitchell, E.A.D., Wilkinson, D.M., Adl, S., Bonkowski, M., Brown, M.W., Fiore-Donno, A.M., Heger, T.J., Jassey, V.E.J., Krashevska, V., Lahr, D.J.G., Marcisz, K., Mulot, M., Payne, R., Singer, D., Anderson, O.R., Charman, D.J., Ekelund, F., Griffiths, B.S., Rønn, R., Smirnov, A., Bass, D., Belbahri, L., Berney, C., Blandenier, Q., Chatzinotas, A., Clarholm, M., Dunthorn, M., Feest, A., Fernández, L.D., Foissner, W., Fournier, B., Gentekaki, E., Hájek, M., Helder, J., Jousset, A., Koller, R., Kumar, S., La Terza, A., Lamentowicz, M., Mazei, Y., Santos, S.S., Seppey, C.V.W., Spiegel, F.W., Walochnik, J., Winding, A., Lara, E., 2017. Soil protistology rebooted: 30 fundamental questions to start with. Soil Biology and Biochemistry 111, 94–103. doi:10.1016/j.soilbio.2017.04.001

54. Geisen, S., Tveit, A.T., Clark, I.M., Richter, A., Svenning, M.M., Bonkowski, M., Urich, T., 2015. Metatranscriptomic census of active protists in soils. The ISME Journal 9, 2178– 2190. doi:10.1038/ismej.2015.30

55. Glaser, K., Kuppardt, A., Boenigk, J., Harms, H., Fetzer, I., Chatzinotas, A., 2015. The influence of environmental factors on protistan microorganisms in grassland soils along a land-use gradient. Science of the Total Environment 537, 33–42. doi:10.1016/j.scitotenv.2015.07.158

56. Gleason, F.H., Letcher, P.M., McGee, P.A., 2004. Some Chytridiomycota in soil recover from drying and high temperatures. Mycological Research 108, 583–589. doi:10.1017/S0953756204009736

57. Gloor, G.B., Macklaim, J.M., Pawlowsky-Glahn, V., Egozcue, J.J., 2017. Microbiome datasets are compositional: And this is not optional. Frontiers in Microbiology 8, 1–6. doi:10.3389/fmicb.2017.02224

58. Glücksman, E., Bell, T., Griffiths, R.I., Bass, D., 2010. Closely related protist strains have different grazing impacts on natural bacterial communities. Environmental Microbiology 12, 3105–3113. doi:10.1111/j.1462-2920.2010.02283.x

59. Goberna, M., Verdú, M., 2022. Cautionary notes on the use of co-occurrence networks in soil ecology 166.

60. Graham, E.B., Knelman, J.E., 2023. Implications of Soil Microbial Community Assembly for Ecosystem Restoration: Patterns, Process, and Potential. Microbial Ecology 85, 809–819. doi:10.1007/s00248-022-02155-w

61. Grossmann, L., Jensen, M., Heider, D., Jost, S., Glücksman, E., Hartikainen, H., Mahamdallie, S.S., Gardner, M., Hoffmann, D., Bass, D., Boenigk, J., 2016. Protistan community analysis: Key findings of a large-scale molecular sampling. ISME Journal 10, 2269–2279. doi:10.1038/ismej.2016.10

62. Guo, S., Xiong, W., Hang, X., Gao, Z., Jiao, Z., Liu, H., Mo, Y., Zhang, N., Kowalchuk, G.A., Li, R., Shen, Q., Geisen, S., 2021. Protists as main indicators and determinants of plant performance. Microbiome 9, 1–11. doi:10.1186/s40168-021-01025-w

63. Guo, Y., Hou, L., Zhang, Z., Zhang, J., Cheng, J., Wei, G., Lin, Y., 2019. Soil microbial diversity during 30 years of grassland restoration on the Loess Plateau, China: Tight linkages with plant diversity. Land Degradation and Development 30, 1172–1182. doi:10.1002/ldr.3300

64. Guo, Y., Xu, T., Cheng, J., Wei, G., Lin, Y., 2021. Above– and belowground biodiversity drives soil multifunctionality along a long-term grassland restoration chronosequence. Science of the Total Environment 772, 145010. doi:10.1016/j.scitotenv.2021.145010

65. Hanrahan-Tan, D.G., Lilje, O., Henderson, L., 2023a. Chytrids in Soil Environments: Unique Adaptations and Distributions. Encyclopedia 3, 642–664. doi:10.3390/encyclopedia3020046

66. Hanrahan-Tan, D.G., Lilje, O., Henderson, L., 2023b. Chytrids in Soil Environments: Unique Adaptations and Distributions. Encyclopedia 3, 642–664. doi:10.3390/encyclopedia3020046

67. Harris, J., 2009. Soil microbial communities and restoration ecology: Facilitators or followers? Science 325, 573–574. doi:10.1126/science.1172975

68. Herzberger, A.J., Duncan, D.S., Jackson, R.D., 2014. Bouncing back: Plant-associated soil microbes respond rapidly to prairie establishment. PLoS ONE 9, 1–14. doi:10.1371/journal.pone.0115775

69. Hess, S., Suthaus, A., 2022. The Vampyrellid Amoebae (Vampyrellida, Rhizaria). Protist 173, 125854. doi:10.1016/j.protis.2021.125854

70. Hu, P., Xiao, J., Zhang, W., Xiao, L., Yang, R., Xiao, D., Zhao, J., Wang, K., 2020. Response of soil microbial communities to natural and managed vegetation restoration in a subtropical karst region. Catena 195, 104849. doi:10.1016/j.catena.2020.104849

71. Hugerth, L.W., Muller, E.E.L., Hu, Y.O.O., Lebrun, L.A.M., Roume, H., Lundin, D., Wilmes, P., Andersson, A.F., 2014. Systematic design of 18S rRNA gene primers for determining eukaryotic diversity in microbial consortia. PLoS ONE 9, e95567. doi:10.1371/journal.pone.0095567

72. Jangid, K., Williams, M.A., Franzluebbers, A.J., Blair, J.M., Coleman, D.C., Whitman, W.B., 2010. Development of soil microbial communities during tallgrass prairie restoration. Soil Biology and Biochemistry 42, 302–312. doi:10.1016/j.soilbio.2009.11.008

73. Kabir, Z., 2005. Tillage or no-tillage: Impact on mycorrhizae. Canadian Journal of Plant Science 85, 23–29. doi:10.4141/P03-160

74. Kalam, S., Basu, A., Ahmad, I., Sayyed, R.Z., El-Enshasy, H.A., Dailin, D.J., Suriani, N.L., 2020. Recent Understanding of Soil Acidobacteria and Their Ecological Significance: A Critical Review. Frontiers in Microbiology 11. doi:10.3389/fmicb.2020.580024

75. Kemmerling, L.R., Rutkoski, C.E., Evans, S.E., Helms, J.A., Cordova-Ortiz, E.S., Smith, J.D., Vázquez Custodio, J.A., Vizza, C., Haddad, N.M., 2022. Prairie Strips and Lower Land Use Intensity Increase Biodiversity and Ecosystem Services. Frontiers in Ecology and Evolution 10, 1–18. doi:10.3389/fevo.2022.833170

76. Kielak, A.M., Barreto, C.C., Kowalchuk, G.A., van Veen, J.A., Kuramae, E.E., 2016. The Ecology of Acidobacteria: Moving beyond Genes and Genomes. Frontiers in Microbiology 7. doi:10.3389/fmicb.2016.00744

77. Kimmell, L.B., Fagan, J.M., Havrilla, C.A., 2023. Soil restoration increases soil health across global drylands: A meta-analysis. Journal of Applied Ecology 60, 1939–1951. doi:10.1111/1365-2664.14459

78. King, W.L., Kaminsky, L.M., Richards, S.C., Bradley, B.A., Kaye, J.P., Bell, T.H., 2022. Farm-scale differentiation of active microbial colonizers. ISME Communications 2, 1–8. doi:10.1038/s43705-022-00120-9

79. Kleijn, D., Bommarco, R., Fijen, T.P.M., Garibaldi, L.A., Potts, S.G., van der Putten, W.H., 2019. Ecological Intensification: Bridging the Gap between Science and Practice. Trends in Ecology and Evolution 34, 154–166. doi:10.1016/j.tree.2018.11.002

80. Kuramae, E.E., Yergeau, E., Wong, L.C., Pijl, A.S., Veen, J.A. Van, Kowalchuk, G.A., 2012. Soil characteristics more strongly influence soil bacterial communities than land-use type. FEMS Microbiology Ecology 79, 12–24. doi:10.1111/j.1574-6941.2011.01192.x

81. Kurtz, Z., Mueller, C., Miraldi, E., Bonneau, R., 2023. _SpiecEasi: Sparse Inverse Covariance for Ecological Statistical Inference_.

82. Kurtz, Z.D., Müller, C.L., Miraldi, E.R., Littman, D.R., Blaser, M.J., Bonneau, R.A., 2015. Sparse and Compositionally Robust Inference of Microbial Ecological Networks. PLOS Computational Biology 11, e1004226. doi:10.1371/journal.pcbi.1004226

83. Labouyrie, M., Ballabio, C., Romero, F., Panagos, P., Jones, A., Schmid, M.W., Mikryukov, V., Dulya, O., Tedersoo, L., Bahram, M., Lugato, E., van der Heijden, M.G.A., Orgiazzi, A., 2023. Patterns in soil microbial diversity across Europe. Nature Communications 14. doi:10.1038/s41467-023-37937-4

84. Lamb, A., Green, R., Bateman, I., Broadmeadow, M., Bruce, T., Burney, J., Carey, P., Chadwick, D., Crane, E., Field, R., Goulding, K., Griffiths, H., Hastings, A., Kasoar, T., Kindred, D., Phalan, B., Pickett, J., Smith, P., Wall, E., Ermgassen, E.K.H.J.Z., Balmford, A., 2016. The potential for land sparing to offset greenhouse gas emissions from agriculture. Nature Climate Change 6, 488–492. doi:10.1038/nclimate2910

85. Leff, J.W., Bardgett, R.D., Wilkinson, A., Jackson, B.G., Pritchard, W.J., De Long, J.R., Oakley, S., Mason, K.E., Ostle, N.J., Johnson, D., Baggs, E.M., Fierer, N., 2018. Predicting the structure of soil communities from plant community taxonomy, phylogeny, and traits. ISME Journal 12, 1794–1805. doi:10.1038/s41396-018-0089-x

86. Lekberg, Y., Arnillas, C.A., Borer, E.T., Bullington, L.S., Fierer, N., Kennedy, P.G., Leff, J.W., Luis, A.D., Seabloom, E.W., Henning, J.A., 2021. Nitrogen and phosphorus fertilization consistently favor pathogenic over mutualistic fungi in grassland soils. Nature Communications 12, 1–8. doi:10.1038/s41467-021-23605-y

87. Li, C., Veum, K.S., Goyne, K.W., Nunes, M.R., Acosta-Martinez, V., 2021. A chronosequence of soil health under tallgrass prairie reconstruction. Applied Soil Ecology 164. doi:10.1016/j.apsoil.2021.103939

88. Li, Y., Li, Z., Cui, S., Jagadamma, S., Zhang, Q., 2019. Residue retention and minimum tillage improve physical environment of the soil in croplands: A global meta-analysis. Soil and Tillage Research 194, 104292. doi:10.1016/j.still.2019.06.009

89. Lin, H., Peddada, S. Das, 2020. Analysis of compositions of microbiomes with bias correction. Nature Communications 11, 3514. doi:10.1038/s41467-020-17041-7

90. Lozano, Y.M., Hortal, S., Armas, C., Pugnaire, F.I., 2014. Interactions among soil, plants, and microorganisms drive secondary succession in a dry environment. Soil Biology and Biochemistry 78, 298–306. doi:10.1016/j.soilbio.2014.08.007

91. Ma, L., Zhou, G., Zhang, J., Jia, Z., Zou, H., Chen, L., Zhang, C., Ma, D., Han, C., Duan, Y., 2024. Long-term conservation tillage enhances microbial carbon use efficiency by altering multitrophic interactions in soil. Science of the Total Environment 915, 170018. doi:10.1016/j.scitotenv.2024.170018

92. MacColl, K., Tosi, M., Chagnon, P.-L., MacDougall, A.S., Dunfield, K.E., Maherali, H., 2024. Prairie restoration promotes the abundance and diversity of mutualistic arbuscular mycorrhizal fungi. Ecological Applications In press. doi:10.1002/eap.2981

93. MacDougall, A.S., Esch, E., Dolezal, A., Kamm, C., Carroll, O., Tosi, M., MacColl, K., Nessel, M., Wilcox, A., Ellsworth, L., Mazzorato, A., Noble, D., Pavusa, M., Ramirez, S., Arce, B., Gutgesell, M., McCann, K., Fraser, E., Fryxell, J., Gilvesy, B., Balpataky, K., Levison, J., Biswas, A., Dunfield, K., Rooney, N., Maherali, H., Husband, B., Steinke, D., DeWaard, J., Ali, G., Prosser, R., Young, A., Sulik, J., Harvey, E., Campbell, M., 2024. Ecosystem services on retired marginal farmland. Frontiers in Ecology and the Environment (under rev.

94. Mackelprang, R., Grube, A.M., Lamendella, R., Jesus, E. da C., Copeland, A., Liang, C., Jackson, R.D., Rice, C.W., Kapucija, S., Parsa, B., Tringe, S.G., Tiedje, J.M., Jansson, J.K., 2018. Microbial community structure and functional potential in cultivated and native tallgrass prairie soils of the midwestern United States. Frontiers in Microbiology 9, 1–15. doi:10.3389/fmicb.2018.01775

95. Maharning, A.R., Mills, A.A.S., Adl, S.M., 2009. Soil community changes during secondary succession to naturalized grasslands 41, 137–147. doi:10.1016/j.apsoil.2008.11.003

96. Malard, L.A., Mod, H.K., Guex, N., Broennimann, O., Yashiro, E., Lara, E., Mitchell, E.A.D., Niculita-Hirzel, H., Guisan, A., 2022. Comparative analysis of diversity and environmental niches of soil bacterial, archaeal, fungal and protist communities reveal niche divergences along environmental gradients in the Alps. Soil Biology and Biochemistry 169, 108674. doi:10.1016/j.soilbio.2022.108674

97. Mao, L., Pan, J., Jiang, S., Shi, G., Qin, M., Zhao, Z., Zhang, Q., An, L., Feng, H., Liu, Y., 2019. Arbuscular mycorrhizal fungal community recovers faster than plant community in historically disturbed Tibetan grasslands. Soil Biology and Biochemistry 134, 131–141. doi:10.1016/j.soilbio.2019.03.026

98. Mariotte, P., Mehrabi, Z., Bezemer, T.M., De Deyn, G.B., Kulmatiski, A., Drigo, B., Veen, G.F. (Ciska), van der Heijden, M.G.A., Kardol, P., 2018. Plant–soil feedback: bridging natural and agricultural sciences. Trends in Ecology and Evolution 33, 129–142. doi:10.1016/j.tree.2017.11.005

99. Martin, M., 2011. Cutadapt removes adapter sequences from high-throughput sequencing reads. EMBnet.Journal 17, 10. doi:10.14806/ej.17.1.200

100. Matamala, R., Jastrow, J.D., Miller, R.M., Garten, C.T., 2008. Temporal changes in C and N stocks of restored prairie: Implications for C sequestration strategies. Ecological Applications 18, 1470–1488. doi:10.1890/07-1609.1

101. Matchado, M.S., Lauber, M., Reitmeier, S., Kacprowski, T., Baumbach, J., Haller, D., List, M., Matchado, M.S., Lauber, M., Reitmeier, S., Kacprowski, T., Baumbach, J., Haller, D., List, M., Steffi, M., Lauber, M., Reitmeier, S., Kacprowski, T., Baumbach, J., Haller, D., List, M., 2021. Network analysis methods for studying microbial communities: A mini review. Computational and Structural Biotechnology Journal 19, 2687–2698. doi:10.1016/j.csbj.2021.05.001

102. Mau, R.L., Hayer, M., Purcell, A.M., Geisen, S., Hungate, B.A., Schwartz, E., 2024. Measurements of soil protist richness and community composition are influenced by primer pair, annealing temperature, and bioinformatics choices. Applied and Environmental Microbiology 90, 1–16. doi:10.1128/aem.00800-24

103. Mazel, F., Malard, L., Niculita-Hirzel, H., Yashiro, E., Mod, H.K., Mitchell, E.A.D., Singer, D., Buri, A., Pinto, E., Guex, N., Lara, E., Guisan, A., 2022. Soil protist function varies with elevation in the Swiss Alps. Environmental Microbiology 24, 1689–1702. doi:10.1111/1462-2920.15686

104. Mazzorato, A.C.M., Esch, E.H., MacDougall, A.S., 2022. Prospects for soil carbon storage on recently retired marginal farmland. Science of the Total Environment 806, 150738. doi:10.1016/j.scitotenv.2021.150738

105. Megonigal, J.P., Hines, M.E., Visscher, P.T., 2003. Anaerobic Metabolism: Linkages to Trace Gases and Aerobic Processes, in: Treatise on Geochemistry. Elsevier, pp. 317–424. doi:10.1016/B0-08-043751-6/08132-9

106. Meinshausen, N., Bühlmann, P., 2006. High-dimensional graphs and variable selection with the Lasso. The Annals of Statistics 34. doi:10.1214/009053606000000281

107. Montecchia, M.S., Tosi, M., Soria, M.A., Vogrig, J.A., Sydorenko, O., Correa, O.S., 2015. Pyrosequencing reveals changes in soil bacterial communities after conversion of Yungas forests to agriculture. PLoS ONE 10, e0119426. doi:10.1371/journal.pone.0119426

108. Morriën, E., Hannula, S.E., Snoek, L.B., Helmsing, N.R., Zweers, H., De Hollander, M., Soto, R.L., Bouffaud, M.L., Buée, M., Dimmers, W., Duyts, H., Geisen, S., Girlanda, M., Griffiths, R.I., Jørgensen, H.B., Jensen, J., Plassart, P., Redecker, D., Schmelz, R.M., Schmidt, O., Thomson, B.C., Tisserant, E., Uroz, S., Winding, A., Bailey, M.J., Bonkowski, M., Faber, J.H., Martin, F., Lemanceau, P., De Boer, W., Van Veen, J.A., Van Der Putten, W.H., 2017. Soil networks become more connected and take up more carbon as nature restoration progresses. Nature Communications 8. doi:10.1038/ncomms14349

109. Nearing, J.T., Douglas, G.M., Hayes, M.G., MacDonald, J., Desai, D.K., Allward, N., Jones, C.M.A., Wright, R.J., Dhanani, A.S., Comeau, A.M., Langille, M.G.I., 2022. Microbiome differential abundance methods produce different results across 38 datasets. Nature Communications 13, 1–16. doi:10.1038/s41467-022-28034-z

110. Noble, D.T., MacDougall, A.S., Levison, J., 2023. Impacts of soil, climate, and phenology on retention of dissolved agricultural nutrients by permanent-cover buffers. Science of The Total Environment 860, 160532. doi:10.1016/j.scitotenv.2022.160532

111. Obregon, D., Fonseca de Souza, L., Mendes, L.W., de Moraes, M.T., Tosi, M., Venturini, A.M., Meyer, K.M., Barbosa de Camargo, P., Bohannan, B.J.M., Mazza Rodrigues, J.L., Dunfield, K.E., Tsai, S.M., 2023. Shifts in functional traits and interactions patterns of soil methane-cycling communities following forest-to-pasture conversion in the Amazon Basin. Molecular Ecology 32, 3257–3275. doi:10.1111/mec.16912

112. Oksanen, J., Simpson, G., Blanchet, F., Kindt, R., Legendre, P., Minchin, P., O’Hara, R., Solymos, P., Stevens, M., Szoecs, E., Wagner, H., Barbour, M., Bedward, M., Bolker, B., Borcard, D., Carvalho, G., Chirico, M., DeCaceres, M., Durand, S., Evangelista, H., FitzJohn, R., Friendly, M., Furneaux, B., Hannigan, G., Hill, M., Lahti, L., McGlinn, D., Ouellette, M., RibeiroCunha, E., Smith, T., Stier, A., TerBraak, C., Weedon, J., 2022. vegan: Community Ecology Package.

113. Oliverio, A.M., Geisen, S., Delgado-Baquerizo, M., Maestre, F.T., Turner, B.L., Fierer, N., 2020. The global-scale distributions of soil protists and their contributions to belowground systems. Science Advances 6, 1–11. doi:10.1126/sciadv.aax8787

114. Pedregosa, F., Varoquaux, G., Gramfort, A., Michel, V., Thirion, B., Grisel, O., Blondel, M., Prettenhofer, P., Weiss, R., Dubourg, V., Vanderplas, J., Passos, A., Cournapeau, D., Brucher, M., Perrot, M., Duchesnay, E., 2011. Scikit-learn: machine learning in python. Journal of Machine Learning Research 12, 2825–2830.

115. Peng, Z., Qian, X., Liu, Y., Li, X., Gao, H., An, Y., Qi, J., Jiang, L., Zhang, Y., Chen, S., Pan, H., Chen, B., Liang, C., van der Heijden, M.G.A., Wei, G., Jiao, S., 2024. Land conversion to agriculture induces taxonomic homogenization of soil microbial communities globally. Nature Communications 15, 1–13. doi:10.1038/s41467-024-47348-8

116. Pinheiro, J., Bates, D., DebRoy, S., Sarkar, D., R Core Team, 2018. nlme: Linear and Nonlinear Mixed Effects Models.

117. Põlme, S., Abarenkov, K., Henrik Nilsson, R., Lindahl, B.D., Clemmensen, K.E., Kauserud, H., Nguyen, N., Kjøller, R., Bates, S.T., Baldrian, P., Frøslev, T.G., Adojaan, K., Vizzini, A., Suija, A., Pfister, D., Baral, H.-O., Järv, H., Madrid, H., Nordén, J., Liu, J.-K., Pawlowska, J., Põldmaa, K., Pärtel, K., Runnel, K., Hansen, K., Larsson, K.-H., Hyde, K.D., Sandoval-Denis, M., Smith, M.E., Toome-Heller, M., Wijayawardene, N.N., Menolli, N., Reynolds, N.K., Drenkhan, R., Maharachchikumbura, S.S.N., Gibertoni, T.B., Læssøe, T., Davis, W., Tokarev, Y., Corrales, A., Soares, A.M., Agan, A., Machado, A.R., Argüelles-Moyao, A., Detheridge, A., de Meiras-Ottoni, A., Verbeken, A., Dutta, A.K., Cui, B.-K., Pradeep, C.K., Marín, C., Stanton, D., Gohar, D., Wanasinghe, D.N., Otsing, E., Aslani, F., Griffith, G.W., Lumbsch, T.H., Grossart, H.-P., Masigol, H., Timling, I., Hiiesalu, I., Oja, J., Kupagme, J.Y., Geml, J., Alvarez-Manjarrez, J., Ilves, K., Loit, K., Adamson, K., Nara, K., Küngas, K., Rojas-Jimenez, K., Bitenieks, K., Irinyi, L., Nagy, L.G., Soonvald, L., Zhou, L.-W., Wagner, L., Aime, M.C., Öpik, M., Mujica, M.I., Metsoja, M., Ryberg, M., Vasar, M., Murata, M., Nelsen, M.P., Cleary, M., Samarakoon, M.C., Doilom, M., Bahram, M., Hagh-Doust, N., Dulya, O., Johnston, P., Kohout, P., Chen, Q., Tian, Q., Nandi, R., Amiri, R., Perera, R.H., dos Santos Chikowski, R., Mendes-Alvarenga, R.L., Garibay-Orijel, R., Gielen, R., Phookamsak, R., Jayawardena, R.S., Rahimlou, S., Karunarathna, S.C., Tibpromma, S., Brown, S.P., Sepp, S.-K., Mundra, S., Luo, Z.-H., Bose, T., Vahter, T., Netherway, T., Yang, T., May, T., Varga, T., Li, W., Coimbra, V.R.M., de Oliveira, V.R.T., de Lima, V.X., Mikryukov, V.S., Lu, Y., Matsuda, Y., Miyamoto, Y., Kõljalg, U., Tedersoo, L., 2020. FungalTraits: a user-friendly traits database of fungi and fungus-like stramenopiles. Fungal Diversity 105, 1–16. doi:10.1007/s13225-020-00466-2

118. R Core Team, 2022. R: A language and environment for statistical computing.

119. Robinson, J.M., Hodgson, R., Krauss, S.L., Liddicoat, C., Malik, A.A., Martin, B.C., Mohr, J.J., Moreno-Mateos, D., Muñoz-Rojas, M., Peddle, S.D., Breed, M.F., 2023. Opportunities and challenges for microbiomics in ecosystem restoration. Trends in Ecology and Evolution 38, 1189–1202. doi:10.1016/j.tree.2023.07.009

120. Romdhane, S., Spor, A., Banerjee, S., Breuil, M.C., Bru, D., Chabbi, A., Hallin, S., van der Heijden, M.G.A., Saghai, A., Philippot, L., 2022. Land-use intensification differentially affects bacterial, fungal and protist communities and decreases microbiome network complexity. Environmental Microbiomes 17, 1–15. doi:10.1186/s40793-021-00396-9

121. Rosenzweig, S.T., Carson, M.A., Baer, S.G., Blair, J.M., 2016. Changes in soil properties, microbial biomass, and fluxes of C and N in soil following post-agricultural grassland restoration. Applied Soil Ecology 100, 186–194. doi:10.1016/j.apsoil.2016.01.001

122. Ruhnau, B., 2000. Eigenvector-centrality — a node-centrality ? Social Networks 22, 357–365.

123. Sáez-Sandino, T., Delgado-Baquerizo, M., Egidi, E., Singh, B.K., 2023. New microbial tools to boost restoration and soil organic matter. Microbial Biotechnology 16, 2019–2025. doi:10.1111/1751-7915.14325

124. Santos, S.S., Schöler, A., Nielsen, T.K., Hansen, L.H., Schloter, M., Winding, A., 2020. Land use as a driver for protist community structure in soils under agricultural use across Europe. Science of the Total Environment 717. doi:10.1016/j.scitotenv.2020.137228

125. Schulz, G., Schneider, D., Brinkmann, N., Edy, N., Daniel, R., Polle, A., Scheu, S., Krashevska, V., 2019. Changes in trophic groups of protists with conversion of rainforest into rubber and oil palm plantations. Frontiers in Microbiology 10, 1–14. doi:10.3389/fmicb.2019.00240

126. Seshadri, R., Kyrpides, N.C., Ivanova, N.N., 2023. Comparative Genomics Using the Integrated Microbial Genomes and Microbiomes (IMG/M) System: A Deinococcus Use Case. Journal of the Indian Institute of Science 103, 877–890. doi:10.1007/s41745-023-00368-7

127. Sha, G., Chen, Y., Wei, T., Guo, X., Yu, H., Jiang, S., Xin, P., Ren, K., 2023a. Responses of soil microbial communities to vegetation restoration on the Loess Plateau of China: A meta-analysis. Applied Soil Ecology 189, 104910. doi:10.1016/j.apsoil.2023.104910

128. Sha, G., Chen, Y., Wei, T., Guo, X., Yu, H., Jiang, S., Xin, P., Ren, K., 2023b. Responses of soil microbial communities to vegetation restoration on the Loess Plateau of China: A meta-analysis. Applied Soil Ecology 189, 104910. doi:10.1016/j.apsoil.2023.104910

129. Shen, C., Wang, J., He, J.Z., Yu, F.H., Ge, Y., 2021. Plant Diversity Enhances Soil Fungal Diversity and Microbial Resistance to Plant Invasion. Applied and Environmental Microbiology 87, 1–15. doi:10.1128/AEM.00251-21

130. Shu, X., Liu, W., Hu, Y., Xia, L., Fan, K., Zhang, Y., Zhang, Y., Zhou, W., 2023. Ecosystem multifunctionality and soil microbial communities in response to ecological restoration in an alpine degraded grassland. Frontiers in Plant Science 14, 1–12. doi:10.3389/fpls.2023.1173962

131. Smith, R.S., Shiel, R.S., Bardgett, R.D., Millward, D., Corkhill, P., Rolph, G., Hobbs, P.J., Peacock, S., 2003. Soil microbial community, fertility, vegetation and diversity as targets in the restoration management of a meadow grassland. Journal of Applied Ecology 40, 51–64. doi:10.1046/j.1365-2664.2003.00780.x

132. Song, L., 2023. Toward Understanding Microbial Ecology to Restore a Degraded Ecosystem. International Journal of Environmental Research and Public Health 20. doi:10.3390/ijerph20054647

133. Strickland, M.S., Callaham, M.A., Gardiner, E.S., Stanturf, J.A., Leff, J.W., Fierer, N., Bradford, M.A., 2017. Response of soil microbial community composition and function to a bottomland forest restoration intensity gradient. Applied Soil Ecology 119, 317–326. doi:10.1016/j.apsoil.2017.07.008

134. Tedersoo, L., Bahram, M., Põlme, S., Kõljalg, U., Yorou, N.S., Wijesundera, R., Ruiz, L.V., Vasco-Palacios, A.M., Thu, P.Q., Suija, A., Smith, M.E., Sharp, C., Saluveer, E., Saitta, A., Rosas, M., Riit, T., Ratkowsky, D., Pritsch, K., Põldmaa, K., Piepenbring, M., Phosri, C., Peterson, M., Parts, K., Pärtel, K., Otsing, E., Nouhra, E., Njouonkou, A.L., Nilsson, R.H., Morgado, L.N., Mayor, J., May, T.W., Majuakim, L., Lodge, D.J., Lee, S.S., Larsson, K.-H., Kohout, P., Hosaka, K., Hiiesalu, I., Henkel, T.W., Harend, H., Guo, L., Greslebin, A., Grelet, G., Geml, J., Gates, G., Dunstan, W., Dunk, C., Drenkhan, R., Dearnaley, J., De Kesel, A., Dang, T., Chen, X., Buegger, F., Brearley, F.Q., Bonito, G., Anslan, S., Abell, S., Abarenkov, K., 2014. Global diversity and geography of soil fungi. Science 346, 1052– 1053. doi:10.1126/science.1256688

135. Tosi, M., Drummelsmith, J., Obregón, D., Chahal, I., Van Eerd, L.L., Dunfield, K.E., 2022. Cover crop-driven shifts in soil microbial communities could modulate early tomato biomass via plant-soil feedbacks. Scientific Reports 12, 1–13. doi:10.1038/s41598-022-11845-x

136. Trivedi, P., Delgado-Baquerizo, M., Anderson, I.C., Singh, B.K., 2016. Response of Soil Properties and Microbial Communities to Agriculture: Implications for Primary Productivity and Soil Health Indicators. Frontiers in Plant Science 7, 1–13. doi:10.3389/fpls.2016.00990

137. Vainio, E.J., Hantula, J., 2000. Direct analysis of wood-inhabiting fungi using denaturing gradient gel electrophoresis of amplified ribosomal DNA. Mycological Research 104, 927– 936. doi:10.1017/S0953756200002471

138. van der Bij, A.U., Pawlett, M., Harris, J.A., Ritz, K., van Diggelen, R., 2017. Soil microbial community assembly precedes vegetation development after drastic techniques to mitigate effects of nitrogen deposition. Biological Conservation 212, 476–483. doi:10.1016/j.biocon.2016.09.008

139. Vaulot, D., Mahé, F., Bass, D., Geisen, S., 2022. pr2-primer: An 18S rRNA primer database for protists. Molecular Ecology Resources 22, 168–179. doi:10.1111⁄1755-0998.13465

140. Verbruggen, E., Rillig, M.C., Wehner, J., Hegglin, D., Wittwer, R., van der Heijden, M.G.A., 2014. Sebacinales, but not total root associated fungal communities, are affected by land-use intensity. New Phytologist 203, 1036–1040. doi:10.1111/nph.12884

141. Wagg, C., Schlaeppi, K., Banerjee, S., Kuramae, E.E., van der Heijden, M.G.A., 2019. Fungal-bacterial diversity and microbiome complexity predict ecosystem functioning. Nature Communications 10, 1–10. doi:10.1038/s41467-019-12798-y

142. Wang, L., Li, J., Zhang, S., Huang, Y., Ouyang, Z., Mai, Z., 2024. Biological soil crust elicits microbial community and extracellular polymeric substances restructuring to reduce the soil erosion on tropical island, South China Sea. Marine Environmental Research 197, 106449. doi:10.1016/j.marenvres.2024.106449

143. Wang, M.Y., Hu, L.B., Wang, W.H., Liu, S.T., Li, M., Liu, R.J., 2009. Influence of long-term fixed fertilization on diversity of arbuscular mycorrhizal fungi. Pedosphere 19, 663–672.

144. Wezel, A., Casagrande, M., Celette, F., Vian, J.F., Ferrer, A., Peigné, J., 2014. Agroecological practices for sustainable agriculture. A review. Agronomy for Sustainable Development 34, 1–20. doi:10.1007/s13593-013-0180-7

145. Xiong, W., Jousset, A., Li, R., Delgado-Baquerizo, M., Bahram, M., Logares, R., Wilden, B., de Groot, G.A., Amacker, N., Kowalchuk, G.A., Shen, Q., Geisen, S., 2021. A global overview of the trophic structure within microbiomes across ecosystems. Environment International 151, 106438. doi:10.1016/j.envint.2021.106438

146. Xu, H., Chen, C., Pang, Z., Zhang, G., Wu, J., Kan, H., 2022. Short-Term Vegetation Restoration Enhances the Complexity of Soil Fungal Network and Decreased the Complexity of Bacterial Network. Journal of Fungi 8. doi:10.3390/jof8111122

147. Yang, T., Lupwayi, N., Marc, S.A., Siddique, K.H.M., Bainard, L.D., 2021. Anthropogenic drivers of soil microbial communities and impacts on soil biological functions in agroecosystems. Global Ecology and Conservation 27, e01521. doi:10.1016/j.gecco.2021.e01521

148. Yang, Y., Dou, Y., Wang, B., Xue, Z., Wang, Y., An, S., Chang, S.X., 2023. Deciphering factors driving soil microbial life-history strategies in restored grasslands. IMeta 2, 1–21. doi:10.1002/imt2.66

149. Yang, Y., Hobbie, S.E., Hernandez, R.R., Fargione, J., Grodsky, S.M., Tilman, D., Zhu, Y.G., Luo, Y., Smith, T.M., Jungers, J.M., Yang, M., Chen, W.Q., 2020. Restoring Abandoned Farmland to Mitigate Climate Change on a Full Earth. One Earth 3, 176–186. doi:10.1016/j.oneear.2020.07.019

150. Youssef, N.H., Farag, I.F., Rinke, C., Hallam, S.J., Woyke, T., Elshahed, M.S., 2015. In Silico Analysis of the Metabolic Potential and Niche Specialization of Candidate Phylum “Latescibacteria” (WS3). PLOS ONE 10, e0127499. doi:10.1371/journal.pone.0127499

151. Zhao, Z., He, J., Quan, Z., Wu, C., Sheng, R., Zhang, L., 2020. Fertilization changes soil microbiome functioning, especially phagotrophic protists. Soil Biology and Biochemistry 148, 107863. doi:10.1016/j.soilbio.2020.107863

152. Zhao, Z.B., He, J.Z., Geisen, S., Han, L.L., Wang, J.T., Shen, J.P., Wei, W.X., Fang, Y.T., Li, P.P., Zhang, L.M., 2019. Protist communities are more sensitive to nitrogen fertilization than other microorganisms in diverse agricultural soils. Microbiome 7, 1–16. doi:10.1186/s40168-019-0647-0

