## Supplementary material for "Complex responses of soil prokaryotes, fungi and protists to prairie restoration on retired agricultural lands"

### Supplementary methods

*Detailed description of protist sequencing data bioinformatics*

Protist data was processed by previously concatenating or joining paired-end reads to maximize the recovery of reads and taxa (Dacey & Chain, 2021). Even though the 18S rRNA fragment obtained with the 616*F/1132R primers has been previously analyzed with merged paired-end reads (Oliverio et al., 2020), this approach is not optimal. The reason is that this fragment size can surpass the Illumina MiSeq PE300 limit for several protist groups, including some common soil taxa (*e.g.*, Amoebozoa, Excavata and Opisthokonta) (Vaulot et al., 2022). Thus, those reads end up being excluded because they do not overlap and are not able to merge. Similar concerns were recently raised by Mau et al. (2024).

In order to recover the highest number of target reads and taxa, we compared the efficacy of four different approaches for processing our reads: forward reads only, reverse reads only, merged paired-end reads and concatenated paired-end reads (*i.e.*, joined without overlap). As found by Dacey and Chain (2021) with lower quality 16S rRNA sequences, the best results were obtained by concatenating previously trimmed reads. The concatenating procedure was carried out by adapting a Python script built and provided in their publication (Dacey & Chain, 2021). Among the options provided by the authors, concatenating length-trimmed reads recovered more protist reads and taxa than concatenating quality-trimmed reads (both with the custom Python script) or than merging and concatenating with PANDAseq. Briefly, cutadapt was used to trim forward and reverse primers, and to length-trim or truncate forward and reverse reads to a set length (150 bp for forward reads and 180 bp for reverse reads). Truncation length was chosen to remove low quality regions at the end of forward and reverse reads, as well as to avoid introducing redundant data in the case of reads which were long enough to merge.

Trimmed and concatenated reads were then imported into QIIME2 and then input into q2-dada2 (Callahan et al., 2016) for denoising, dereplication and chimera filtering of reads. After this step, we obtained a total of 656,512 reads and 9645 ASVs before removing non-target sequences. Taxonomy was assigned using q2-feature-classifier classify-sklearn (Bokulich, Dillon, et al., 2018) with a taxonomic classifier trained using the $PR^2$ (Protist Ribosomal Reference) database v.5.0.0 (Vaulot et al., 2022). $PR^2$ reads were trimmed with the forward and reverse primers used in this study. These reads were then quality-filtered (*i.e.*, removing those with 5+ ambiguous bases and/or with homopolymers of 8+ bases) using q2-rescript cull-seqs and dereplicated using q2-rescript dereplicate in mode uniq (*i.e.*, retaining all sequences with unique taxonomic classifications) (Robeson et al., 2021). Trimmed, quality-filtered and dereplicated 18S rRNA reads from $PR^2$ were used to train the taxonomic classifier using q2-feature-classifier fit-classifier-naive-bayes (Bokulich, Kaehler, et al., 2018; Pedregosa et al., 2011). Non-target reads, only found in 16S rRNA (*i.e.*, chloroplast and mitochondria) and 18S rRNA sequencing, were removed before carrying out downstream analyses.

### Supplementary figures

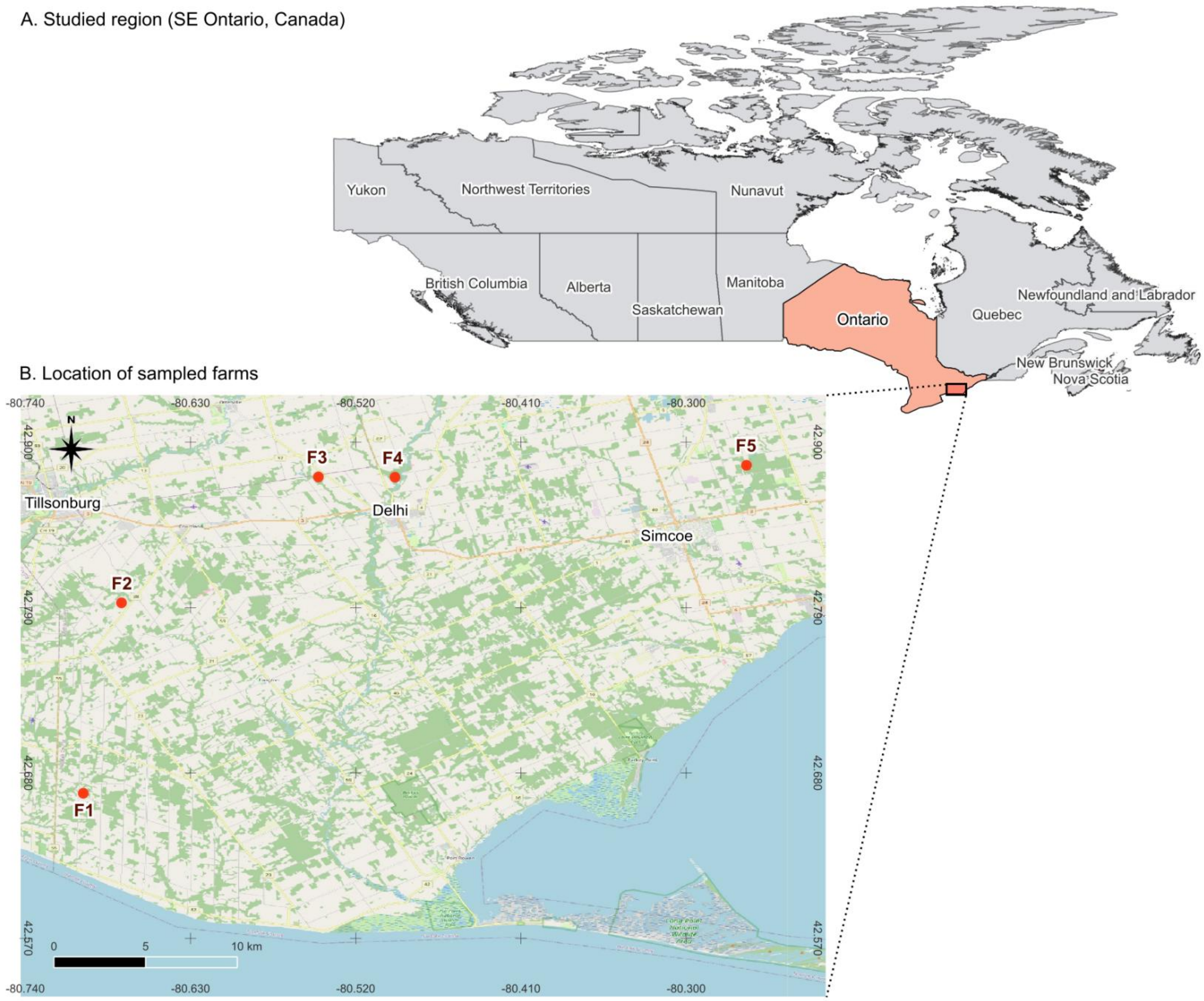


**Figure S1.** Map showing the location of the studied area in the SE of the province of Ontario, Canada (A) and the location of the sampled farms within this area (B). Maps done with QGIS v. 3.34 and compiled with Inkscape v. 1.3.2.

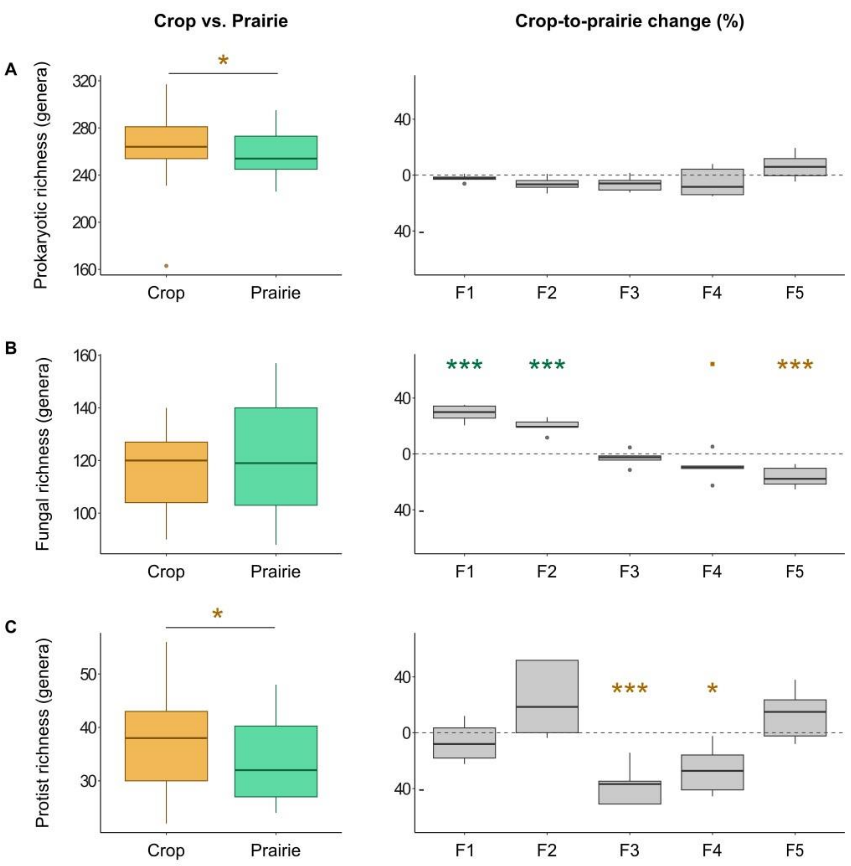


**Figure S2.** Response of soil microbial taxonomic richness to prairie restoration. Average value for all farms (left) and percentage change from crop to prairie by farm (right). *Crop-to-prairie change was calculated from each prairie plot relative to the crop site average of 5 plots (positive=higher in prairie, negative=higher in crop). In average value plots, asterisks represent significant changes based on ANOVA, whereas in percentage change plots, asterisks represent significant differences between prairie and crop soils based on Tukey contrasts (alpha=0.05).*

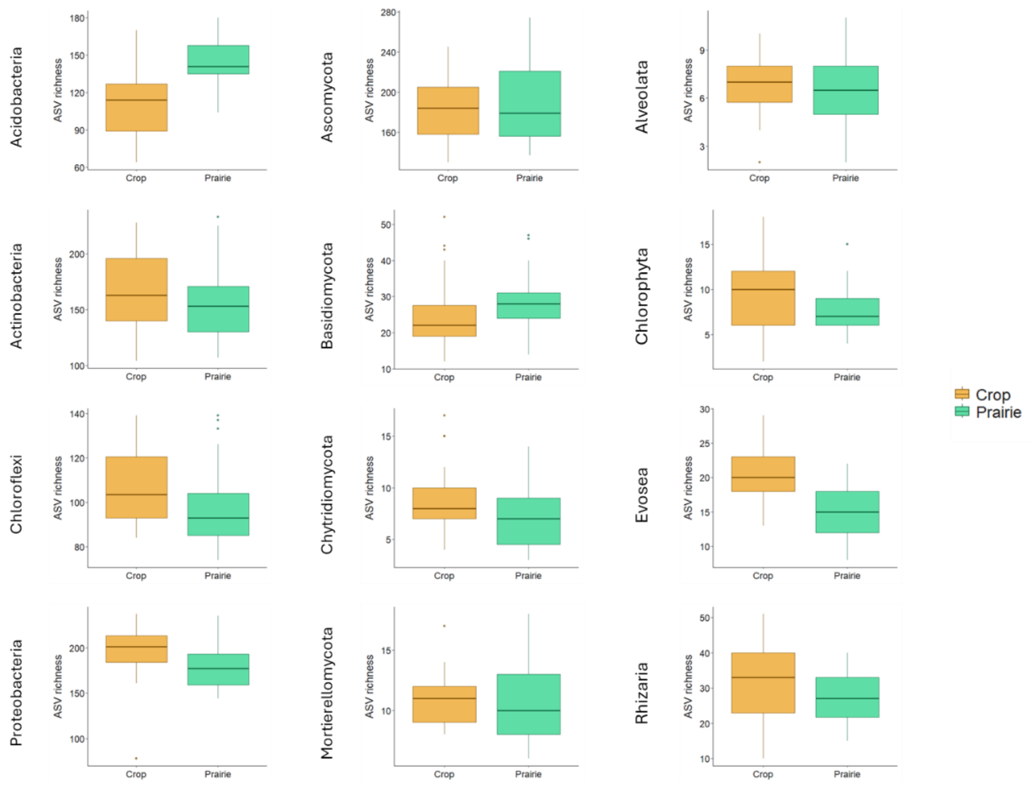


**Figure S3.** Response of soil genetic microbial richness to prairie restoration within the most abundant soil microbial phyla.

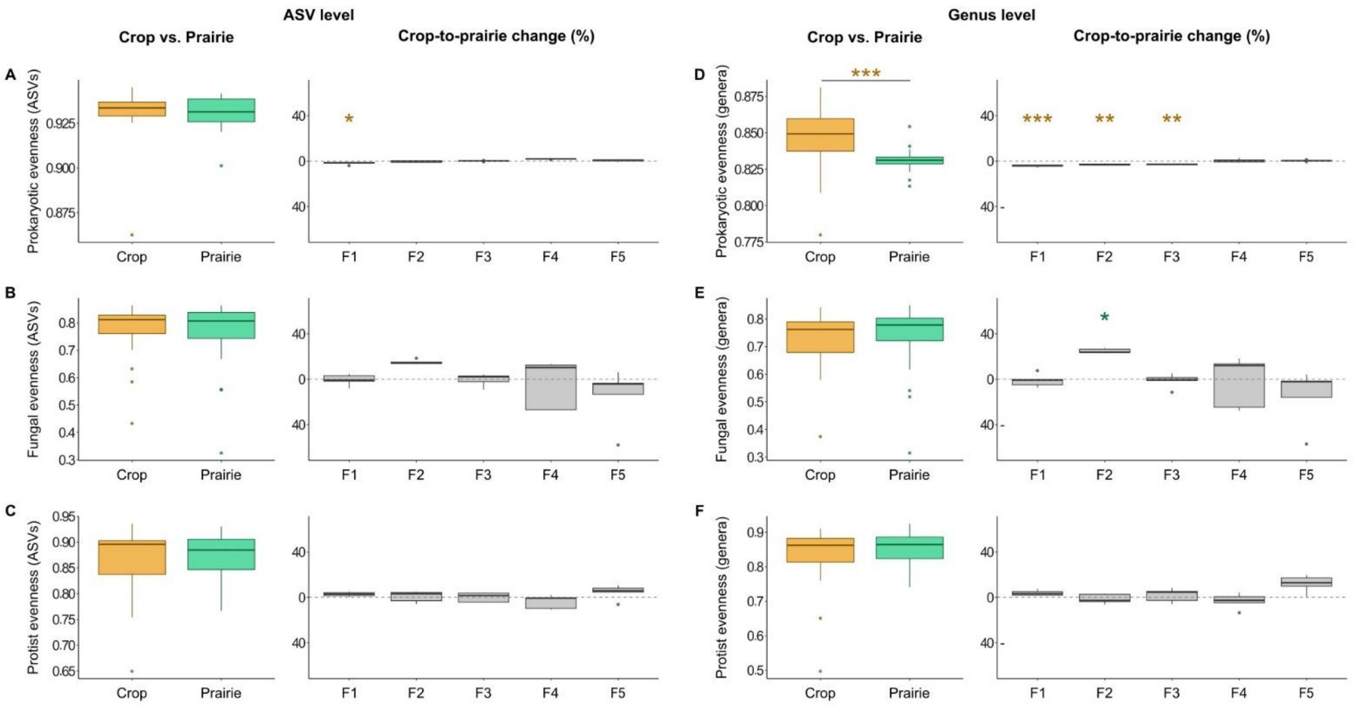


**Figure S4.** Response of measured soil genetic and taxonomic microbial evenness to prairie restoration (ASV and genus level, respectively). Average value for all farms (A, C, E, G, I, K) and percentage change from crop to prairie by farm (B, D, F, H, J, L). *Crop-to-prairie change was calculated from each prairie plot relative to the crop site average of 5 plots (positive=higher in prairie, negative=higher in crop). In average value plots, asterisks represent significant changes based on ANOVA, whereas in percentage change plots, asterisks represent significant differences between prairie and crop soils based on Tukey contrasts (alpha=0.05).*

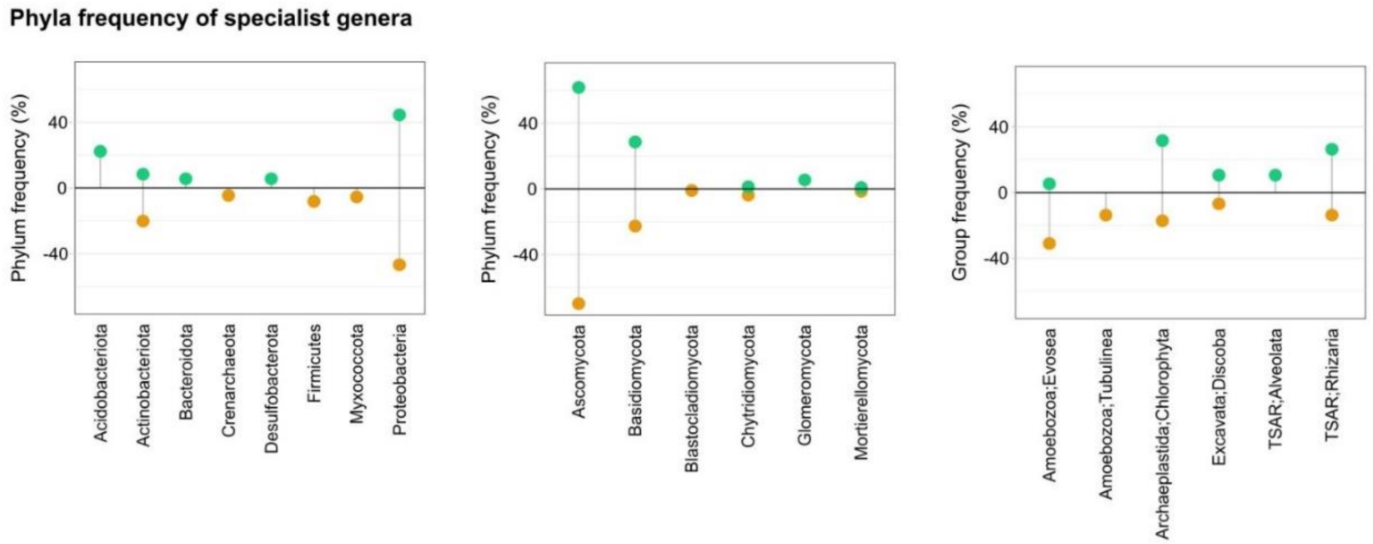


**Figure S5.** Frequency of phyla or major protist groups detected among crop or prairie "specialists" from the analysis of specialist-generalist genera based on the multinomial species classification method (CLAM test).

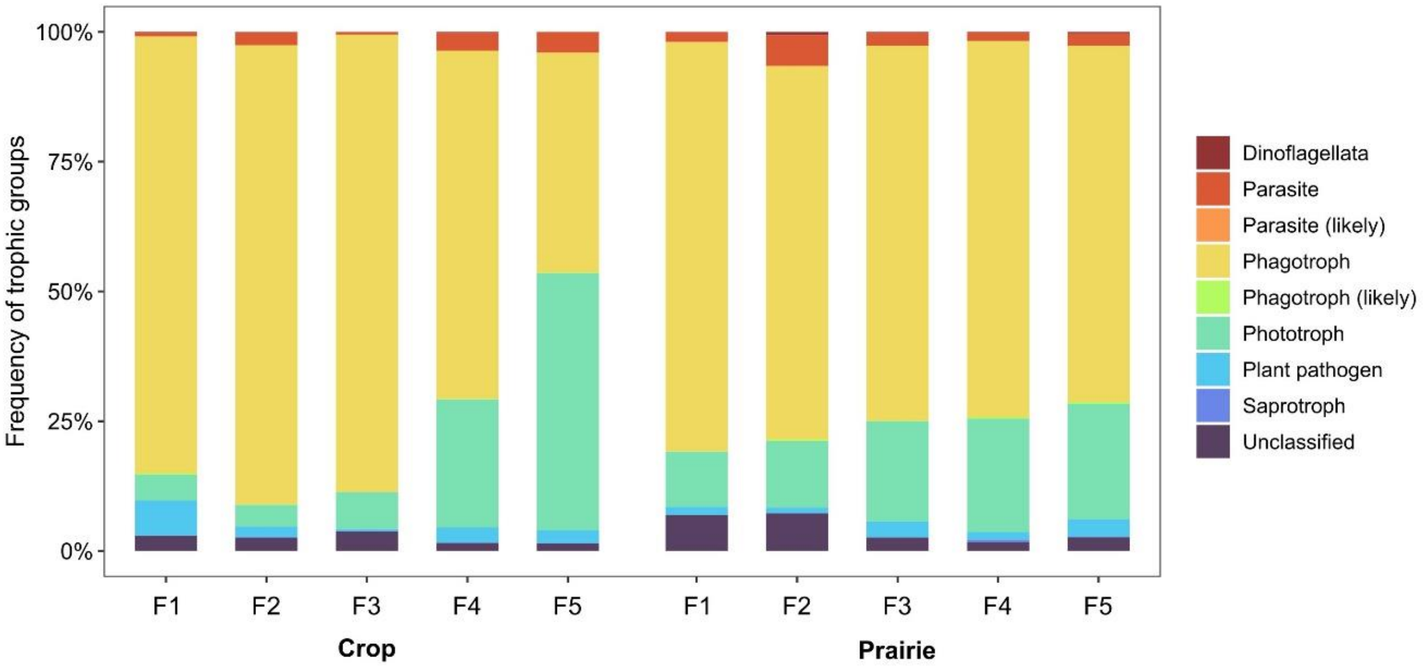


**Figure S6.** Frequency of soil protist trophic groups in each farm and land use (crop and prairie), evidencing the stronger farm gradient within crop soils. ALDEx2 and ANCOM-BC detected a change in the relative abundance of phototrophs across farms (increase from F1 to F5).

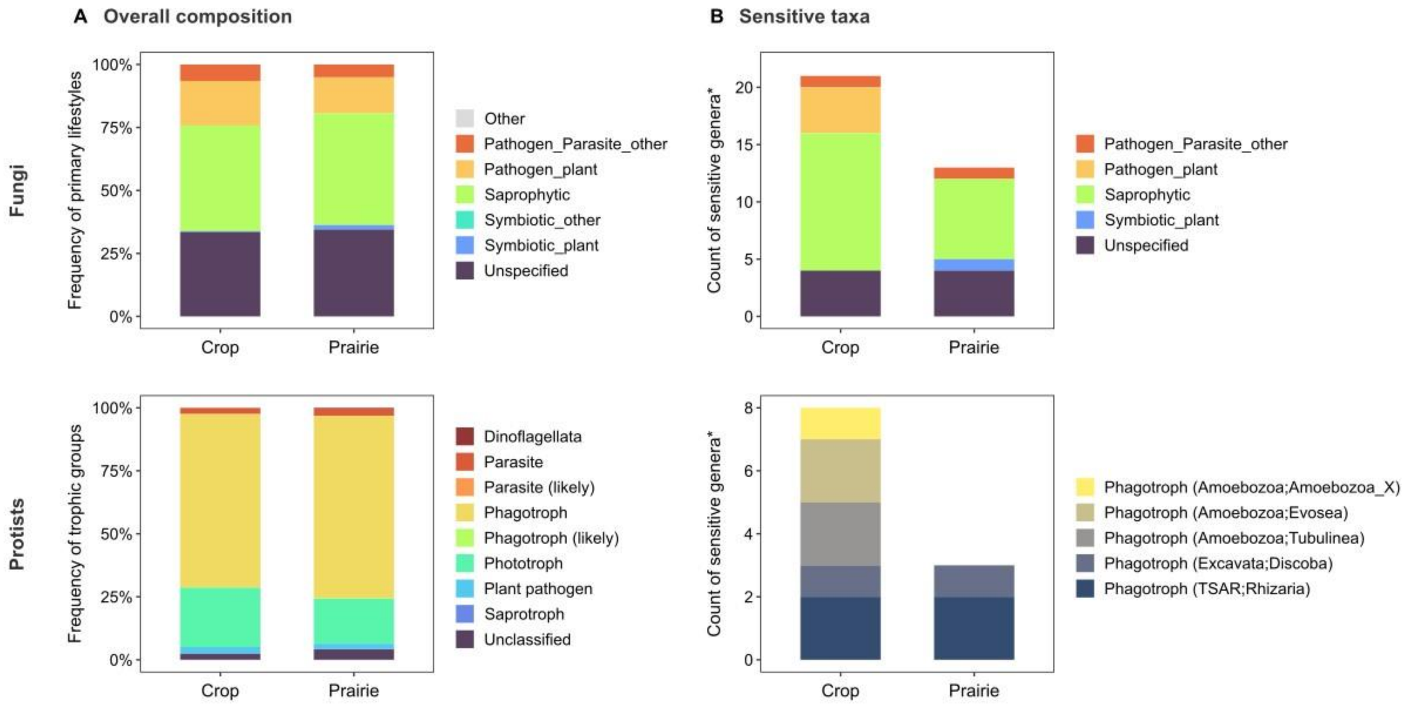


**Figure S7.** Changes in predicted soil fungal and protist functional groups between crop soils and adjacent restored prairie soils. A) Overall composition showing frequency of groups within each land use. B) Functional groups among sensitive genera detected in differential abundance analysis (see Figure 5).

### Supplementary tables

**Table S1.** ANOVA results for soil physicochemical properties, microbial abundance from qPCR and alpha diversity metrics (richness, Pielou's evenness and Shannon Entropy) from DNA sequencing.

| | | Use (numDF=1) | | | Farm (numDF=4) | | | Use:Farm (numDF=4) | | |
|---|---|---|---|---|---|---|---|---|---|---|
| | | F-value | p-value | | F-value | p-value | | F-value | p-value | |
| Physicochemical | Sand | 0.13 | 0.720 | | 37.73 | **<.0001** | *** | 3.20 | **0.023** | * |
| | Silt | 0.21 | 0.653 | | 28.45 | **<.0001** | *** | 3.42 | **0.017** | * |
| | Clay | 0.01 | 0.910 | | 60.93 | **<.0001** | *** | 6.44 | **0.000** | *** |
| | pH | 10.72 | **0.002** | ** | 15.73 | **<.0001** | *** | 11.24 | **<.0001** | *** |
| | OM | 9.62 | **0.004** | ** | 33.02 | **<.0001** | *** | 14.28 | **<.0001** | *** |
| | Mineral N | 73.87 | **<.0001** | *** | 29.19 | **<.0001** | *** | 26.63 | **<.0001** | *** |
| | Ext. P | 3.13 | 0.085 | | 33.80 | **<.0001** | *** | 36.23 | **<.0001** | *** |
| | C:P | 41.03 | **<.0001** | *** | 83.67 | **<.0001** | *** | 107.51 | **<.0001** | *** |
| Abundance (qPCR) | Bacteria | 1.34 | 0.253 | | 6.31 | **0.001** | ** | 4.44 | **0.005** | ** |
| | Fungi | 5.44 | **0.025** | * | 7.51 | **0.000** | *** | 1.04 | 0.398 | |
| | F:B ratio | 9.89 | **0.003** | ** | 1.49 | 0.2232 | | 1.02 | 0.4064 | |
| Richness (ASV level) | Prokaryotes | 1.27 | 0.266 | | 3.45 | **0.016** | * | 0.38 | 0.821 | |
| | Fungi | 2.35 | 0.133 | | 14.73 | **<.0001** | *** | 10.45 | **<.0001** | *** |
| | Protists | 8.69 | **0.005** | ** | 2.80 | **0.039** | * | 7.17 | **0.000** | *** |
| Richness (genus level) | Prokaryotes | 4.40 | **0.042** | * | 5.69 | **0.001** | ** | 0.64 | 0.638 | |
| | Fungi | 2.63 | 0.113 | | 22.93 | **<.0001** | *** | 15.37 | **<.0001** | *** |
| | Protists | 4.63 | **0.038** | * | 3.17 | **0.024** | ** | 5.99 | **0.001** | ** |
| Evenness (ASV level) | Prokaryotes | 0.40 | 0.539 | | 1.70 | 0.168 | | 3.40 | **0.017** | * |
| | Fungi | 0.01 | 0.938 | | 1.98 | 0.117 | | 0.94 | 0.450 | |
| | Protists | 0.13 | 0.725 | | 1.35 | 0.270 | | 0.71 | 0.593 | |
| Evenness (genus level) | Prokaryotes | 17.90 | **0.000** | *** | 0.67 | 0.619 | | 4.56 | **0.004** | ** |
| | Fungi | 0.00 | 0.980 | | 2.19 | 0.088 | | 1.75 | 0.158 | |
| | Protists | 0.32 | 0.574 | | 1.48 | 0.227 | | 1.02 | 0.408 | |
| Shannon (ASV level) | Prokaryotes | 0.00 | 0.951 | | 12.56 | **<.0001** | *** | 1.06 | 0.390 | |
| | Fungi | 0.58 | 0.451 | | 3.34 | **0.019** | * | 1.69 | 0.172 | |
| | Protists | 6.30 | **0.016** | * | 0.90 | 0.471 | | 5.67 | **0.001** | ** |
| Shannon (genus level) | Prokaryotes | 20.20 | **0.000** | *** | 6.20 | **0.001** | ** | 5.50 | **0.001** | ** |
| | Fungi | 0.65 | 0.425 | | 4.12 | **0.007** | ** | 2.51 | 0.057 | |
| | Protists | 1.51 | 0.227 | | 1.78 | 0.152 | | 4.02 | **0.008** | ** |

Degrees of freedom: use=1, farm=4, use:farm=4

C:P was log transformed because of its highly skewed distribution.

Bacterial and fungal abundance analyzed as log copies per gram dry soil.

Alpha diversity metrics were analyzed at the amplicon sequence variant (ASV) and genus level.

**Table S2.** Number of total reads, amplicon sequence variants (ASVs), genera and phyla recovered before and after filtering very rare ASVs (*i.e.*, with less than 10 total reads and/or present in less than 2 sites).

| | **Prokaryotes** | | | **Fungi** | | | **Protists** | | |
|---|---|---|---|---|---|---|---|---|---|
| | **No filter** | **Filter** | **% Lost** | **No filter** | **Filter** | **% Lost** | **No filter** | **Filter** | **% Lost** |
| Reads | 1,197,342 | 1,075,138 | 10 | 2,612,111 | 2,325,256 | 11 | 83,591 | 63,178 | 24 |
| ASVs | 17,725 | 6,002 | 66 | 5,411 | 1,765 | 67 | 2,225 | 528 | 76 |
| Genera | 1090 | 677 | 38 | 813 | 469 | 42 | 376 | 180 | 52 |
| Phyla* | 46 | 38 | 17 | 16 | 11 | 31 | 18 | 14 | 22 |

* Note about protists (level 3)
Unidentified were removed for phyla counts (*e.g.*, unid bacteria or fungi)
Filtering removed those ASVs present in only one sample and with less than 10 reads (so pretty relaxed)

**Table S3.** PERMANOVA results showing land use and farm effects on microbial composition. Results are shown for a qualitative (Jaccard) and quantitative (Bray-Curtis) metric, both at the ASV and genus level.

| | | | Prokaryotes[a] | | | Fungi | | | Protists | | |
|---|---|---|---|---|---|---|---|---|---|---|---|
| | | | $R^2$ | F | P | $R^2$ | F | P | $R^2$ | F | P |
| ASV | Jaccard | Use | 0.05 | 2.80 | 0.001*** | 0.06 | 3.94 | 0.001*** | 0.05 | 2.72 | 0.001*** |
| | | Farm | 0.18 | 2.80 | 0.001*** | 0.21 | 3.57 | 0.001*** | 0.15 | 2.18 | 0.001*** |
| | | Use:Farm | 0.12 | 1.82 | 0.001*** | 0.14 | 2.45 | 0.001*** | 0.12 | 1.67 | 0.001*** |
| | | Residual | 0.65 | | | 0.59 | | | 0.68 | | |
| | Bray-Curtis | Use | 0.07 | 5.54 | 0.001*** | 0.07 | 5.28 | 0.001*** | 0.07 | 4.80 | 0.001*** |
| | | Farm | 0.27 | 5.08 | 0.001*** | 0.22 | 4.04 | 0.001*** | 0.19 | 3.15 | 0.001*** |
| | | Use:Farm | 0.14 | 2.74 | 0.001*** | 0.17 | 3.06 | 0.001*** | 0.15 | 2.40 | 0.001*** |
| | | Residual | 0.52 | | | 0.54 | | | 0.59 | | |
| Genus | Jaccard | Use | 0.11 | 8.65 | 0.001*** | 0.07 | 4.41 | 0.001*** | 0.08 | 4.61 | 0.001*** |
| | | Farm | 0.24 | 4.90 | 0.001*** | 0.17 | 2.88 | 0.001*** | 0.15 | 2.39 | 0.001*** |
| | | Use:Farm | 0.16 | 3.16 | 0.001*** | 0.16 | 2.58 | 0.001*** | 0.14 | 2.15 | 0.001*** |
| | | Residual | 0.49 | | | 0.60 | | | 0.63 | | |
| | Bray-Curtis | Use | 0.13 | 12.09 | 0.001*** | 0.09 | 6.94 | 0.001*** | 0.11 | 7.65 | 0.001*** |
| | | Farm | 0.28 | 6.80 | 0.001*** | 0.21 | 4.09 | 0.001*** | 0.19 | 3.45 | 0.001*** |
| | | Use:Farm | 0.17 | 4.00 | 0.001*** | 0.18 | 3.49 | 0.001*** | 0.16 | 2.94 | 0.001*** |
| | | Residual | 0.42 | | | 0.52 | | | 0.54 | | |

[a] Betadisper test evidenced higher dispersion in crop than prairie prokaryotic community composition (alpha=0.05) for both features (ASV/genus) and distances (Jaccard/Bray-Curtis). For ASV tests, also higher dispersion in F4 than other farms. No other significant differences in beta dispersion were detected.

P-value codes: *** <.001, ** <.01, * <.05, · <.010

**Table S4.** PERMANOVA results by farm (ASV level)

| | | **Jaccard** | | | | **Bray-Curtis** | | | |
|---|---|---|---|---|---|---|---|---|---|
| | **Farm** | **Betadisper** | **R-squared** | **F** | **Pr(>F)** | **Betadisper** | **R-squared** | **F** | **Pr(>F)** |
| **Prokaryotes** | F1 | n.s. | 0.23 | 2.43 | 0.005 ** | crop>prairie** | 0.36 | 4.51 | 0.005 ** |
| | F2 | n.s. | 0.19 | 1.94 | 0.005 ** | n.s. | 0.27 | 2.89 | 0.005 ** |
| | F3 | n.s. | 0.23 | 2.36 | 0.005 ** | n.s. | 0.33 | 3.89 | 0.005 ** |
| | F4 | n.s. | 0.15 | 1.38 | 0.018 * | n.s. | 0.18 | 1.71 | 0.024 * |
| | F5 | n.s. | 0.20 | 1.97 | 0.005 ** | n.s. | 0.31 | 3.51 | 0.005 ** |
| **Fungi** | F1 | n.s. | 0.23 | 2.44 | 0.005 ** | n.s. | 0.25 | 2.64 | 0.005 ** |
| | F2 | n.s. | 0.27 | 3.03 | 0.005 ** | n.s. | 0.37 | 4.72 | 0.005 ** |
| | F3 | n.s. | 0.27 | 2.99 | 0.005 ** | n.s. | 0.33 | 4.00 | 0.005 ** |
| | F4 | prairie>crop* | 0.21 | 2.12 | 0.005 ** | prairie>crop* | 0.26 | 2.86 | 0.005 ** |
| | F5 | n.s. | 0.29 | 3.19 | 0.005 ** | n.s. | 0.30 | 3.43 | 0.005 ** |
| **Protists** | F1 | n.s. | 0.20 | 1.72 | 0.006 ** | n.s. | 0.22 | 2.01 | 0.010 ** |
| | F2 | n.s. | 0.20 | 2.02 | 0.005 ** | n.s. | 0.26 | 2.78 | 0.005 ** |
| | F3 | n.s. | 0.21 | 2.09 | 0.005 ** | n.s. | 0.33 | 3.86 | 0.005 ** |
| | F4 | n.s. | 0.19 | 1.82 | 0.005 ** | n.s. | 0.29 | 3.30 | 0.005 ** |
| | F5 | n.s. | 0.18 | 1.77 | 0.005 ** | n.s. | 0.24 | 2.58 | 0.005 ** |

**Table S5.** Species turnover and nestedness values calculated by comparing microbial species presence/absence data between crop field and restored prairie plots at each farm site, accompanied by comparisons with a null expectation

| | | Turnover | | Nestedness | | | |
|---|---|---|---|---|---|---|---|
| | Farm | Obs. | Exp. | Obs. | Exp. | SES turnover* | P-value |
| Prokaryotes | F1 | 0.990 | 0.985 | 0.010 | 0.015 | 4.291 | <0.001 |
| | F2 | 0.973 | 0.974 | 0.027 | 0.026 | -0.557 | 0.566 |
| | F3 | 0.984 | 0.977 | 0.016 | 0.023 | 3.392 | <0.001 |
| | F4 | 0.981 | 0.977 | 0.019 | 0.023 | 3.003 | <0.001 |
| | F5 | 0.964 | 0.963 | 0.036 | 0.037 | 0.288 | 0.942 |
| Fungi | F1 | 0.951 | 0.952 | 0.049 | 0.048 | -0.466 | 0.522 |
| | F2 | 0.985 | 0.979 | 0.015 | 0.021 | 3.760 | <0.001 |
| | F3 | 0.986 | 0.980 | 0.014 | 0.020 | 3.636 | <0.001 |
| | F4 | 0.972 | 0.969 | 0.028 | 0.031 | 1.312 | 0.052 |
| | F5 | 0.976 | 0.970 | 0.024 | 0.030 | 2.698 | <0.001 |
| Protists | F1 | 0.986 | 0.980 | 0.014 | 0.020 | 2.144 | 0.002 |
| | F2 | 0.977 | 0.972 | 0.023 | 0.028 | 1.799 | 0.026 |
| | F3 | 0.953 | 0.951 | 0.047 | 0.049 | 0.307 | 0.864 |
| | F4 | 0.978 | 0.973 | 0.022 | 0.027 | 1.601 | 0.034 |
| | F5 | 0.949 | 0.953 | 0.051 | 0.047 | -0.846 | 0.320 |

Mean species turnover and nestedness were calculated based on pairwise comparisons between all pairs of crop field and restored prairies samples.

Obs.: observed values (Obs.), Exp.: expected value based on 999 randomly generated scenarios, SES: standardized effect size for each observation [(Obs. – Exp.) / $SD_{Exp.}$], P: which represents the position of the observed value within the null distribution.

* Turnover and nestedness are reciprocal: SES nestedness = –(SES turnover)

**Table S6.** Differential abundance for prokaryotes

| | | ANCOM-BC | | | ALDEx2 | | |
|---|---|---|---|---|---|---|---|
| Higher in | Taxonomic id. | LFC | SE | W | Effect | CLR crop | CLR prairie |
| Crop | Archaea;Crenarchaeota;Nitrososphaeria;Nitrososphaerales;Nitrososphaeraceae;Candidatus_Nitrocosmicus | -0.644 | 0.161 | -4.014 | -0.70 | 6.91 | 6.03 |
| | Bacteria;Deinococcota;Deinococci;Deinococcales;Deinococcaceae;Deinococcus | -1.353 | 0.297 | -4.559 | -0.69 | 2.11 | -1.51 |
| | Bacteria;Proteobacteria;Gammaproteobacteria;Burkholderiales;Nitrosomonadaceae;Nitrosospira | -2.690 | 0.372 | -7.233 | -1.13 | 4.68 | -1.81 |
| | Bacteria;Proteobacteria;Gammaproteobacteria;Xanthomonadales;Xanthomonadaceae;Luteimonas | -2.582 | 0.390 | -6.627 | -0.98 | 4.77 | -1.41 |
| Prairie | Bacteria;Acidobacteriota;Blastocatellia;11-24;11-24;11-24 | 1.007 | 0.147 | 6.860 | 1.23 | 5.78 | 7.34 |
| | Bacteria;Acidobacteriota;Blastocatellia;Blastocatellales;Blastocatellaceae;Aridibacter | 2.131 | 0.366 | 5.824 | 0.81 | -0.24 | 4.65 |
| | Bacteria;Acidobacteriota;Blastocatellia;Blastocatellales;Blastocatellaceae;Blastocatellaceae | 1.815 | 0.450 | 4.031 | 0.68 | -1.75 | 3.54 |
| | Bacteria;Acidobacteriota;Blastocatellia;Pyrinomonadales;Pyrinomonadaceae;RB41 | 1.111 | 0.118 | 9.436 | 1.74 | 7.63 | 9.23 |
| | Bacteria;Acidobacteriota;Vicinamibacteria;Subgroup_17;Subgroup_17;Subgroup_17 | 0.596 | 0.127 | 4.679 | 0.78 | 6.73 | 7.53 |
| | Bacteria;Acidobacteriota;Vicinamibacteria;Vicinamibacterales;uncultured;uncultured | 0.594 | 0.066 | 9.031 | 1.54 | 9.45 | 10.35 |
| | Bacteria;Actinobacteriota;Actinobacteria;Streptosporangiales;Thermomonosporaceae;Actinocorallia | 1.316 | 0.294 | 4.468 | 0.63 | 1.93 | 3.87 |
| | Bacteria;Chloroflexi;Anaerolineae;SBR1031;SBR1031;SBR1031 | 0.787 | 0.170 | 4.630 | 0.72 | 5.60 | 6.44 |
| | Bacteria;Chloroflexi;Chloroflexia;Chloroflexales;Roseiflexaceae;uncultured | 0.923 | 0.162 | 5.678 | 0.99 | 5.37 | 6.65 |
| | Bacteria;Myxococcota;Polyangia;Haliangiales;Haliangiaceae;Haliangium | 0.452 | 0.089 | 5.076 | 0.80 | 6.02 | 6.63 |
| | Bacteria;Planctomycetota;Phycisphaerae;S-70;S-70;S-70 | 1.209 | 0.301 | 4.020 | 0.49 | -0.59 | 2.58 |
| | Bacteria;Planctomycetota;Planctomycetes;Gemmatales;Gemmataceae;uncultured | 0.704 | 0.098 | 7.150 | 1.22 | 6.34 | 7.23 |
| | Bacteria;Planctomycetota;Planctomycetes;Pirellulales;Pirellulaceae;Pir4_lineage | 0.290 | 0.060 | 4.818 | 0.72 | 7.11 | 7.54 |
| | Bacteria;Proteobacteria;Alphaproteobacteria;Rhizobiales;Hyphomicrobiaceae;Pedomicrobium | 0.520 | 0.130 | 3.993 | 0.63 | 5.65 | 6.42 |
| | Bacteria;Proteobacteria;Alphaproteobacteria;Rhizobiales;Rhizobiales_Incertae_Sedis;Nordella | 0.757 | 0.145 | 5.206 | 0.80 | 5.00 | 5.84 |
| | Bacteria;Proteobacteria;Alphaproteobacteria;Rhizobiales;Xanthobacteraceae;uncultured | 0.533 | 0.133 | 3.999 | 0.62 | 6.89 | 7.53 |
| | Bacteria;Proteobacteria;Gammaproteobacteria;Burkholderiales;Comamonadaceae;__ | 0.495 | 0.108 | 4.597 | 0.79 | 6.76 | 7.48 |
| | Bacteria;Proteobacteria;Gammaproteobacteria;Burkholderiales;Comamonadaceae;Rhizobacter | 2.525 | 0.402 | 6.275 | 0.98 | -1.25 | 4.91 |
| | Bacteria;Proteobacteria;Gammaproteobacteria;Burkholderiales;Nitrosomonadaceae;Ellin6067 | 0.564 | 0.142 | 3.983 | 0.71 | 5.73 | 6.64 |
| | Bacteria;Proteobacteria;Gammaproteobacteria;Pseudomonadales;Pseudomonadaceae;Pseudomonas | 1.387 | 0.316 | 4.390 | 0.88 | 5.46 | 7.53 |

**Table S7.** Differentially abundant soil fungal taxa and their predicted lifestyle

| | | 1º lifestyle | | 2º lifestyle | | ANCOM-BC | | | ALDEx2 | | |
|---|---|---|---|---|---|---|---|---|---|---|---|
| Higher in | Taxonomic id. | T.mode | Guild | T.mode | Guild | LFC | SE | W | Effect | CLR crop | CLR prairie |
| Crop | Ascomycota;Dothideomycetes;Pleosporales;Phaeosphaeriaceae;Neosetophoma | Sap | litter saprotroph | Sym | foliar endophyte | -2.75 | 0.56 | -4.90 | -0.71 | 5.41 | -0.38 |
| | Ascomycota;Dothideomycetes;Pleosporales;Phaeosphaeriaceae;Ophiosphaerella | Pat | plant pathogen | Sap | litter saprotroph | -2.03 | 0.51 | -3.96 | -0.54 | 5.17 | 0.69 |
| | Ascomycota;Dothideomycetes;Pleosporales;Pleosporaceae;Bipolaris | Pat | plant pathogen | Sap | litter saprotroph | -2.39 | 0.51 | -4.72 | -0.66 | 1.31 | -1.97 |
| | Ascomycota;Dothideomycetes;Pleosporales;Sporormiaceae;unidentified | Uns | | Uns | | -2.54 | 0.56 | -4.56 | -0.67 | 4.50 | -1.70 |
| | Ascomycota;Leotiomycetes;Helotiales;Ploettnerulaceae;Cadophora | Sap | litter saprotroph | Pat | plant pathogen | -1.82 | 0.46 | -3.94 | -0.51 | 4.25 | -0.72 |
| | Ascomycota;Sordariomycetes;Glomerellales;Plectosphaerellaceae;_ | Uns | | Uns | | -0.95 | 0.22 | -4.36 | -0.56 | 10.62 | 9.70 |
| | Ascomycota;Sordariomycetes;Glomerellales;Plectosphaerellaceae;Plectosphaerella | Pat | plant pathogen | Sap | litter saprotroph | -3.10 | 0.57 | -5.46 | -0.76 | 7.99 | 3.20 |
| | Ascomycota;Sordariomycetes;Hypocreales;Nectriaceae;_ | Uns | | Uns | | -1.97 | 0.49 | -4.04 | -0.49 | 8.05 | 6.50 |
| | Ascomycota;Sordariomycetes;Hypocreales;Nectriaceae;Fusarium | Pat | plant pathogen | Sap | litter saprotroph | -0.89 | 0.16 | -5.61 | -0.69 | 12.24 | 11.48 |
| | Ascomycota;Sordariomycetes;Hypocreales;Nectriaceae;Fusicolla | Pat | mycoparasite | Sap | fungal decomposer | -1.73 | 0.46 | -3.77 | -0.50 | 8.15 | 6.92 |
| | Ascomycota;Sordariomycetes;Microascales;Microascaceae;Enterocarpus | Sap | dung saprotroph | Sap | animal decomposer | -3.00 | 0.64 | -4.72 | -0.75 | 6.49 | -0.74 |
| | Ascomycota;Sordariomycetes;Microascales;Microascaceae;Kernia | Sap | dung saprotroph | Uns | | -1.86 | 0.42 | -4.44 | -0.54 | 0.18 | -2.01 |
| | Ascomycota;Sordariomycetes;Myrmecridiales;Myrmecridiaceae;Myrmecridium | Sap | soil saprotroph | Uns | | -2.37 | 0.63 | -3.76 | -0.58 | 6.88 | 1.59 |
| | Ascomycota;Sordariomycetes;Sordariales;Bombardiaceae;Ramophialophora | Sap | soil saprotroph | Uns | | -2.43 | 0.59 | -4.12 | -0.59 | 3.27 | -1.56 |
| | Ascomycota;Sordariomycetes;Sordariales;Podosporaceae;Cladorrhinum | Sap | unspecified saprotroph | Sym | foliar endophyte | -2.66 | 0.67 | -3.94 | -0.51 | 8.05 | 5.08 |
| | Basidiomycota;Agaricomycetes;Agaricales;Psathyrellaceae;Coprinellus | Sap | soil saprotroph | Uns | | -2.64 | 0.58 | -4.58 | -0.68 | 5.47 | -1.36 |
| | Basidiomycota;Agaricostilbomycetes;Agaricostilbales;Chionosphaeraceae;Kurtzmanomyces | Sap | unspecified saprotroph | Uns | | -2.18 | 0.55 | -3.98 | -0.57 | 4.56 | -1.30 |
| | Basidiomycota;Tremellomycetes;Cystofilobasidiales;Mrakiaceae;Mrakia | Sap | unspecified saprotroph | Uns | | -2.40 | 0.48 | -4.97 | -0.65 | 5.54 | 3.23 |
| | Basidiomycota;Tremellomycetes;Cystofilobasidiales;Mrakiaceae;Tausonia | Sap | soil saprotroph | Uns | | -2.88 | 0.62 | -4.65 | -0.73 | 6.90 | 0.35 |
| | Chytridiomycota;GS13;unidentified;unidentified;unidentified | Uns | | Uns | | -1.86 | 0.35 | -5.33 | -0.64 | 2.83 | -1.70 |
| | Chytridiomycota;Rhizophlyctidomycetes;Rhizophlyctidales;Rhizophlyctidaceae;Rhizophlyctis | Sap | litter saprotroph | Pat | algal parasite | -2.09 | 0.52 | -3.99 | -0.60 | 5.06 | -0.18 |
| Prairie | Ascomycota;Dothideomycetes;Mycosphaerellales;Extremaceae;Incertomyces | Sap | unspecified saprotroph | Other | rock-inhabiting | 2.81 | 0.54 | 5.20 | 0.90 | -1.49 | 5.96 |
| | Ascomycota;Dothideomycetes;Pleosporales;Cucurbitariaceae;Pyrenochaeta | Sap | wood saprotroph | Uns | | 4.68 | 0.39 | 11.93 | 2.17 | -1.77 | 7.75 |
| | Ascomycota;Dothideomycetes;Pleosporales;Didymosphaeriaceae;Paraphaeosphaeria | Sap | wood saprotroph | Uns | | 2.34 | 0.50 | 4.66 | 0.84 | -1.28 | 5.44 |
| | Ascomycota;Dothideomycetes;Pleosporales;Phaeosphaeriaceae;_ | Uns | | Uns | | 1.34 | 0.34 | 3.89 | 0.62 | -1.75 | 2.69 |
| | Ascomycota;Dothideomycetes;Pleosporales;Phaeosphaeriaceae;Wojnowiciella | Sap | litter saprotroph | Sap | wood saprotroph | 1.58 | 0.37 | 4.26 | 0.66 | -1.98 | 1.67 |
| | Ascomycota;Leotiomycetes;Helotiales;Helotiaceae;Allophylaria | Sap | litter saprotroph | Uns | | 2.74 | 0.70 | 3.93 | 0.65 | -1.20 | 7.59 |
| | Ascomycota;Orbiliomycetes;Orbiliales;_;_ | Uns | | Uns | | 1.99 | 0.41 | 4.84 | 0.81 | 3.01 | 6.63 |
| | Ascomycota;Pezizomycotina_cls_Incertae_sedis;unidentified;unidentified;unidentified | Uns | | Uns | | 4.75 | 0.51 | 9.39 | 1.72 | -0.96 | 9.00 |
| | Ascomycota;Sordariomycetes;Branch06;unidentified;unidentified | Uns | | Uns | | 2.42 | 0.47 | 5.15 | 0.97 | -0.54 | 5.37 |
| | Ascomycota;Sordariomycetes;Hypocreales;Ophiocordycipitaceae;Hirsutella | Pat | animal parasite | Sap | animal decomposer | 2.01 | 0.51 | 3.97 | 0.69 | 0.65 | 6.45 |
| | Basidiomycota;Agaricomycetes;Sebacinales;Serendipitaceae;Serendipita | Sym | root endophyte | Sap | soil saprotroph | 4.19 | 0.46 | 9.10 | 1.55 | -0.95 | 8.96 |
| | Basidiomycota;Tremellomycetes;Tremellales;Bulleribasidiaceae;Vishniacozyma | Sap | soil saprotroph | Uns | | 2.07 | 0.55 | 3.77 | 0.71 | -1.02 | 5.92 |
| | Mortierellomycota;Mortierellomycetes;Mortierellales;Mortierellaceae;Mortierella | Sap | soil saprotroph | Sym | root-associated | 0.99 | 0.26 | 3.88 | 0.91 | 8.80 | 10.23 |

**Table S8.** Differentially abundant soil protist taxa and their predicted trophic mode

| Higher in | Taxonomic id. | Trophic mode | ANCOM-BC | | | ALDEx2 | | |
|---|---|---|---|---|---|---|---|---|
| | | | LFC | SE | W | Effect | CLR crop | CLR prairie |
| Crop | Amoebozoa;Amoebozoa_X;Amoebozoa_XX;Lobosa-G1;Lobosa-G1_X;Lobosa-G1_XX;Lobosa-G1_XXX | Phagotroph | -1.25 | 0.31 | -4.00 | -0.59 | 3.96 | 0.22 |
| | Amoebozoa;Evosea;Evosea_X;Variosea;Fractovitellida;Schizoplasmodiidae;__ | Phagotroph | -1.71 | 0.28 | -6.10 | -0.81 | 6.84 | 4.83 |
| | Amoebozoa;Evosea;Evosea_X;Variosea;Variosea_X;Filamoebidae;Filamoeba | Phagotroph | -1.87 | 0.28 | -6.71 | -1.07 | 7.02 | 5.38 |
| | Amoebozoa;Tubulinea;Tubulinea_X;Echinamoebida;Echinamoebida_X;Echinamoebidae;Echinamoeba | Phagotroph | -2.10 | 0.29 | -7.25 | -1.01 | 4.87 | -0.60 |
| | Amoebozoa;Tubulinea;Tubulinea_X;Elardia;Leptomyxida;__;__ | Phagotroph | -1.26 | 0.28 | -4.46 | -0.63 | 2.82 | -1.05 |
| | Excavata;Discoba;Discoba_X;Heterolobosea;Acrasida;Acrasidae;Allovahlkampfia | Phagotroph | -1.83 | 0.34 | -5.43 | -0.82 | 5.10 | 0.77 |
| | TSAR;Rhizaria;Cercozoa;Filosa-Sarcomonadea;Glissomonadida;Glissomonadida_X;Glissomonadida_XX | Phagotroph | -1.48 | 0.32 | -4.61 | -0.60 | 3.95 | -0.11 |
| | TSAR;Rhizaria;Cercozoa;Filosa-Sarcomonadea;Glissomonadida;Sandonidae;Neoheteromita | Phagotroph | -1.70 | 0.43 | -3.95 | -0.63 | 5.02 | -0.45 |
| Prairie | Excavata;Discoba;Euglenozoa;Kinetoplastea;__;__;__ | Phagotroph | 0.97 | 0.22 | 4.46 | 0.59 | -1.14 | 2.16 |
| | TSAR;Rhizaria;Cercozoa;Endomyxa;Vampyrellida;sm27-lineage;sm27-lineage_X | Phagotroph | 1.82 | 0.44 | 4.18 | 0.63 | -0.56 | 4.95 |
| | TSAR;Rhizaria;Cercozoa;Endomyxa;Vampyrellida;Vampyrellidae;Vampyrella | Phagotroph | 1.89 | 0.36 | 5.29 | 0.77 | 3.59 | 6.82 |

**Table S8.** Influential nodes based on degree and eigenvector centrality

| Use | Group | Short Id. | Full Id. | Degree | Eigencentrality |
|---|---|---|---|---|---|
| Crop | Prokaryotes | *Nonomuraea* | p_Actinobacteriota;c_Actinobacteria;o_Streptosporangiales;f_Streptosporangiaceae;g_Nonomuraea | 12 | 1.00 |
| | | *Blastococcus* | p_Actinobacteriota;c_Actinobacteria;o_Frankiales;f_Geodermatophilaceae;g_Blastococcus | 9 | 0.58 |
| | | *Rhodanobacter* | p_Proteobacteria;c_Gammaproteobacteria;o_Xanthomonadales;f_Rhodanobacteraceae;g_Rhodanobacter | 9 | 0.41 |
| | | Vicinamibacteraceae | p_Acidobacteriota;c_Vicinamibacteria;o_Vicinamibacterales;f_Vicinamibacteraceae;_ | 8 | 0.70 |
| | | OM190 | p_Planctomycetota;c_OM190;o_OM190;f_OM190;g_OM190 | 8 | 0.59 |
| | | Steroidobacteraceae | p_Proteobacteria;c_Gammaproteobacteria;o_Steroidobacterales;f_Steroidobacteraceae;_ | 8 | 0.55 |
| | | *Gaiella* | p_Actinobacteriota;c_Thermoleophilia;o_Gaiellales;f_Gaiellaceae;g_Gaiella | 8 | 0.48 |
| | | Sutterellaceae | p_Proteobacteria;c_Gammaproteobacteria;o_Burkholderiales;f_Sutterellaceae;g_uncultured | 7 | 0.60 |
| | | *Streptomyces* | p_Actinobacteriota;c_Actinobacteria;o_Streptomycetales;f_Streptomycetaceae;g_Streptomyces | 7 | 0.56 |
| | | *Geodermatophilus* | p_Actinobacteriota;c_Actinobacteria;o_Frankiales;f_Geodermatophilaceae;g_Geodermatophilus | 7 | 0.55 |
| | | AD3 | p_Chloroflexi;c_AD3;o_AD3;f_AD3;g_AD3 | 7 | 0.52 |
| | | Subgroup_22 | p_Acidobacteriota;c_Subgroup_22;o_Subgroup_22;f_Subgroup_22;g_Subgroup_22 | 7 | 0.43 |
| | | *Mesorhizobium* | p_Proteobacteria;c_Alphaproteobacteria;o_Rhizobiales;f_Rhizobiaceae;g_Mesorhizobium | 7 | 0.41 |
| | | Micropepsaceae | p_Proteobacteria;c_Alphaproteobacteria;o_Micropepsales;f_Micropepsaceae;g_uncultured | 7 | 0.40 |
| | | *Rubellimicrobium* | p_Proteobacteria;c_Alphaproteobacteria;o_Rhodobacterales;f_Rhodobacteraceae;g_Rubellimicrobium | 7 | 0.19 |
| | | *Nocardioides* | p_Actinobacteriota;c_Actinobacteria;o_Propionibacteriales;f_Nocardioidaceae;g_Nocardioides | 7 | 0.14 |
| | | *Candidatus_Xiphinematobacter* | p_Verrucomicrobiota;c_Verrucomicrobiae;o_Chthoniobacterales;f_Xiphinematobacteraceae;g_Candidatus_Xiphinematobacter | 6 | 0.55 |
| | Fungi | *Aaosphaeria* | p_Ascomycota;c_Dothideomycetes;o_Pleosporales;f_Pleosporales_fam_Incertae_sedis;g_Aaosphaeria | 12 | 0.59 |
| | | *Staphylotrichum* | p_Ascomycota;c_Sordariomycetes;o_Sordariales;f_Sordariales_fam_Incertae_sedis;g_Staphylotrichum | 8 | 0.41 |
| | | *Pleotrichocladium* | p_Ascomycota;c_Dothideomycetes;o_Pleosporales;f_Melanommataceae;g_Pleotrichocladium | 7 | 0.53 |
| | | *Thelonectria* | p_Ascomycota;c_Sordariomycetes;o_Hypocreales;f_Nectriaceae;g_Thelonectria | 7 | 0.31 |
| | | *Pyrenochaetopsis* | p_Ascomycota;c_Dothideomycetes;o_Pleosporales;f_Cucurbitariaceae;g_Pyrenochaetopsis | 7 | 0.28 |
| | Protists | WIM80-lineage_X | Amoebozoa;Evosea;Evosea_X;Variosea;Flamellidae;WIM80-lineage;WIM80-lineage_X | 12 | 0.74 |
| | | *Chlamydomonas* | Archaeplastida;Chlorophyta;Chlorophyta_X;Chlorophyceae;Chlamydomonadales;Chlamydomonadales_X;Chlamydomonas | 7 | 0.34 |
| Prairie | Prokaryotes | AD3 | p_Chloroflexi;c_AD3;o_AD3;f_AD3;g_AD3 | 13 | 1.00 |
| | | *Burkholderia_Caballeronia_Paraburkholderia* | p_Proteobacteria;c_Gammaproteobacteria;o_Burkholderiales;f_Burkholderiaceae;g_Burkholderia-Caballeronia-Paraburkholderia | 11 | 0.65 |
| | | Amb_16S_1323 | p_Proteobacteria;c_Alphaproteobacteria;o_Rhizobiales;f_Amb-16S-1323;g_Amb-16S-1323 | 9 | 0.75 |
| | | *Blastococcus* | p_Actinobacteriota;c_Actinobacteria;o_Frankiales;f_Geodermatophilaceae;g_Blastococcus | 9 | 0.46 |
| | | AKAU4049 | p_Gemmatimonadota;c_AKAU4049;o_AKAU4049;f_AKAU4049;g_AKAU4049 | 8 | 0.71 |
| | | *Oryzihumus* | p_Actinobacteriota;c_Actinobacteria;o_Micrococcales;f_Intrasporangiaceae;g_Oryzihumus | 8 | 0.19 |
| | | *Rubrobacter* | p_Actinobacteriota;c_Rubrobacteria;o_Rubrobacterales;f_Rubrobacteriaceae;g_Rubrobacter | 7 | 0.67 |
| | | *Candidatus_Udaeobacter* | p_Verrucomicrobiota;c_Verrucomicrobiae;o_Chthoniobacterales;f_Chthoniobacteraceae;g_Candidatus_Udaeobacter | 7 | 0.48 |
| | | Alphaproteobacteria | p_Proteobacteria;c_Alphaproteobacteria;_;_;_ | 7 | 0.40 |
| | | *Dactylosporangium* | p_Actinobacteriota;c_Actinobacteria;o_Micromonosporales;f_Micromonosporaceae;g_Dactylosporangium | 7 | 0.35 |
| | | *Acidothermus* | p_Actinobacteriota;c_Actinobacteria;o_Frankiales;f_Acidothermaceae;g_Acidothermus | 7 | 0.30 |
| | Protists | sm27-lineage_X | TSAR;Rhizaria;Cercozoa;Endomyxa;Vampyrellida;sm27-lineage;sm27-lineage_X | 7 | 0.50 |

All prokaryotes were bacteria.
Shaded taxa were found in both crop and prairie networks.

**References**


Bokulich, N. A., Dillon, M., Bolyen, E., Kaehler, B., Huttley, G., & Caporaso, J. (2018). Q2-Sample-Classifier: Machine-Learning Tools for Microbiome Classification and Regression. *Journal of Open Source Software*, *3*(30), 934. https://doi.org/10.21105/joss.00934

Bokulich, N. A., Kaehler, B. D., Rideout, J. R., Dillon, M., Bolyen, E., Knight, R., Huttley, G. A., & Gregory Caporaso, J. (2018). Optimizing taxonomic classification of marker-gene amplicon sequences with QIIME 2's q2-feature-classifier plugin. *Microbiome*, *6*(1), 90. https://doi.org/10.1186/s40168-018-0470-z

Callahan, B. J., McMurdie, P. J., Rosen, M. J., Han, A. W., Johnson, A. J. A., & Holmes, S. P. (2016). DADA2: High-resolution sample inference from Illumina amplicon data. *Nature Methods*, *13*(7), 581–583. https://doi.org/10.1038/nmeth.3869

Dacey, D. P., & Chain, F. J. J. (2021). Concatenation of paired-end reads improves taxonomic classification of amplicons for profiling microbial communities. *BMC Bioinformatics*, *22*, 493. https://doi.org/10.1186/s12859-021-04410-2

Oliverio, A. M., Geisen, S., Delgado-baquerizo, M., Maestre, F. T., Turner, B. L., & Fierer, N. (2020). *The global-scale distributions of soil protists and their contributions to belowground systems*. *January*, 1–11.

Pedregosa, F., Varoquaux, G., Gramfort, A., Michel, V., Thirion, B., Grisel, O., Blondel, M., Prettenhofer, P., Weiss, R., Dubourg, V., Vanderplas, J., Passos, A., Cournapeau, D., Brucher, M., Perrot, M., & Duchesnay, E. (2011). Scikit-learn: machine learning in python. *Journal of Machine Learning Research*, *12*(Oct), 2825–2830.

Robeson, M. S., O'Rourke, D. R., Kaehler, B. D., Ziemski, M., Dillon, M. R., Foster, J. T., & Bokulich, N. A. (2021). RESCRIPt: Reproducible sequence taxonomy reference database management. *PLOS Computational Biology*, *17*(11), e1009581. https://doi.org/10.1371/journal.pcbi.1009581

Vaulot, D., Mahé, F., Bass, D., & Geisen, S. (2022). pr2-primer : An 18S rRNA primer database for protists. *Molecular Ecology Resources*, *22*(1), 168–179. https://doi.org/10.1111/1755-0998.13465
